# Simulating the growth of TAF15 inclusions in neuron soma

**DOI:** 10.1101/2024.07.14.603428

**Authors:** Andrey V. Kuznetsov

## Abstract

To the best of the author’s knowledge, this paper presents the first attempt to develop a mathematical model of the formation and growth of inclusions containing misfolded TATA-box binding protein associated factor 15 (TAF15). It has recently been shown that TAF15 inclusions are involved in approximately 10% of cases of frontotemporal lobar degeneration (FTLD). FTLD is the second most common neurodegenerative disease after Alzheimer’s disease (AD). It is characterized by a progressive loss of personality, behavioral changes, and a decline in language skills due to the degeneration of the frontal and anterior temporal lobes. The model simulates TAF15 monomer production, nucleation and autocatalytic growth of free TAF15 aggregates, and their deposition into TAF15 inclusions. The accuracy of the numerical solution of the model equations is validated by comparing it with analytical solutions available for limiting cases. Physiologically relevant parameter values were used to predict TAF15 inclusion growth. It is shown that the growth of TAF15 inclusions is influenced by two opposing mechanisms: the rate at which free TAF15 aggregates are deposited into inclusions and the rate of autocatalytic production of free TAF15 aggregates from monomers. A low deposition rate slows inclusion growth, while a high deposition rate hinders the autocatalytic production of new aggregates, thus also slowing inclusion growth. Consequently, the rate of inclusion growth is maximized at an intermediate deposition rate of free TAF15 aggregates into TAF15 inclusions.

## 1. Introduction

Frontotemporal lobar degeneration (FTLD) ranks as the second most prevalent neurodegenerative disease after Alzheimer’s disease (AD). Characterized by progressive behavioral and language deterioration, FTLD stems from degeneration in the frontal and anterior temporal lobes. It is now acknowledged as a major cause of dementia, particularly among those under 65. About 50% of FTLD cases feature neuronal inclusions of misfolded TDP-43, while 40% exhibit tau aggregates with amyloid structures, signifying filamentous protein assemblies stabilized by intermolecular β-sheets. A recent Nature publication, ref. [1], used cryo-electron microscopy to reveal that the remaining 10% of FTLD cases involve neuronal cytoplasmic inclusions composed of aggregated TATA-box binding protein associated factor 15 (TAF15). This discovery is significant because the protein implicated in these FTLD cases was previously believed to be protein fused in sarcoma (FUS). Consequently, ref. [1] identified TAF15 as a new protein capable of forming amyloid filaments linked to neurodegenerative disease [2].

TAF15 belongs to the ten-eleven translocation (TET) family of RNA-binding proteins, alongside FUS and protein Ewing’s sarcoma (EWS) [3]. These proteins play roles in gene expression regulation at transcriptional and post-transcriptional levels [4]. While primarily found in the nucleus, TET proteins shuttle between the nucleus and cytosol [5]. Furthermore, TAF15 can interact with a limited number of RNA granules in the cytoplasm [6].

Ref. [7] developed a model of TDP-43 inclusion body growth, exploring the change of its radius with time and extending prior studies on amyloid-β plaques [8-10] and Lewy bodies [11,12]. In ref. [13], a model was developed to simulate the diffusion of amyloid beta (Aβ) monomers, the generation of free Aβ aggregates via nucleation and autocatalytic mechanisms, and their deposition into senile plaques. The present paper extends the approach developed in ref. [13] to simulate the growth of TAF15 inclusions.

## 2. Materials and models

### 2.1. Model equations

The Finke-Watzky (F-W) model is a simplified two-step process simulating the formation of certain aggregates, which involves two pseudo-elementary reaction steps, nucleation and autocatalysis. Ref. [14] demonstrated the F-W model’s effectiveness in simulating the aggregation of various proteins. During nucleation, new aggregates are continuously formed, while in the autocatalysis step, existing aggregates catalyze their own production [14,15]. The two pseudo-elementary reaction steps are given by the following equations:

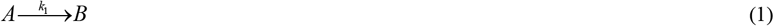

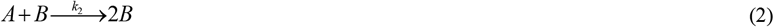

In this model, *A* represents a monomeric protein, while *B* represents a free (not deposited into TAF15 inclusions) amyloid-converted protein. The kinetic constants *k*_1_ and *k*_2_ denote the rates of nucleation and autocatalytic growth, respectively [14]. The primary nucleation process, described by Eq. (1), involves only monomers. In contrast, the secondary nucleation process, described by Eq. (2), involves both monomers and free pre-existing aggregates of the same protein [16].

The concentration of monomers is *C*_*A*_ , while the concentration of free aggregates is *C*_*B*_ . These aggregates include various forms of TAF15 oligomers, protofibrils, and fibrils [17]. TAF15 aggregates can be deposited into TAF15 inclusions, driving their growth. However, the minimalistic nature of the F-W model does not allow it to differentiate between the types and sizes of TAF15 aggregates. Typically, the F-W model is used to investigate the conversion of a given initial concentration of monomers into aggregates. In this work, a novel extension of the F-W model is proposed to address scenarios where TAF15 monomers are continuously generated in the soma (neuron body).

The formation of TAF15 inclusions is a multistep process involving the production of monomers, their conversion into free aggregates through nucleation and autocatalytic processes, and the deposition of these aggregates into growing inclusions. All these steps need to be modeled. To monitor the number of monomers, free aggregates, and deposited aggregates, the soma is chosen as the control volume (CV), as illustrated in Fig. 1. It is assumed that TAF15 monomers are produced at a constant rate, *q* , within the soma. Applying the conservation of TAF15 monomers in the soma results in the following equation:

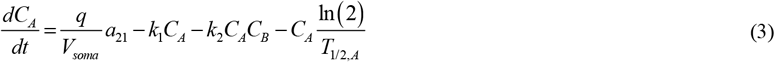

**Fig. 1.**
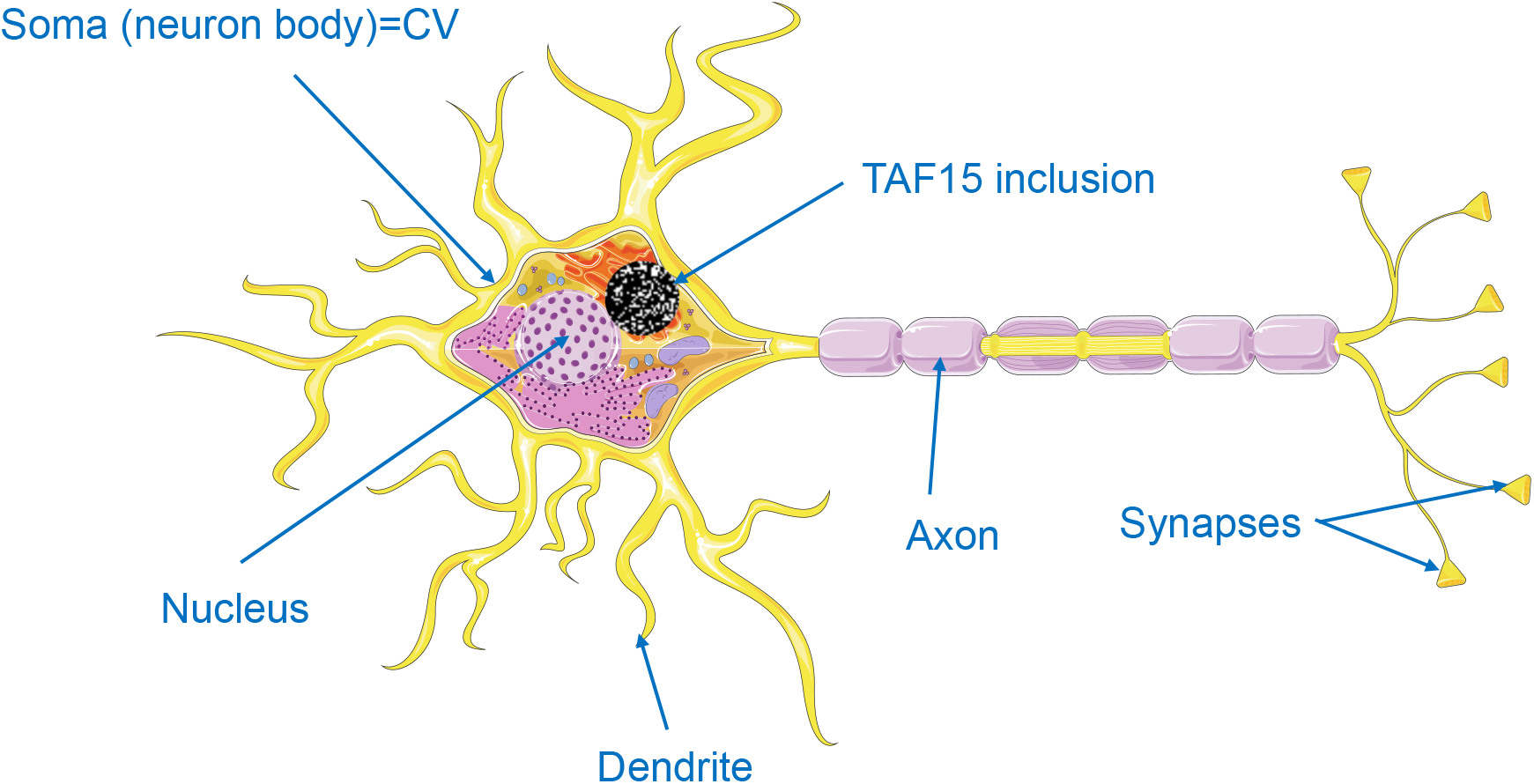
A diagram illustrating a neuron with a TAF15 inclusion situated in the soma. The inclusion in Fig. 1 is not depicted to scale. *Figure generated with the aid of servier medical art, licensed under a creative commons attribution 3*.*0 generic license. http://Smart.servier.com.*

The sole independent variable in the mathematical model is time, *t*. The dependent variables employed in the model are listed in Table 1, and the parameters used are summarized in Table 2. In Eq. (3), 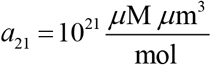 represents the conversion factor from 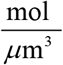 to *μ*M . The first term on the right-hand side of Eq. (3) accounts for the production rate of TAF15 monomers. The second term models the conversion rate of TAF15 monomers into aggregates by nucleation, while the third term describes the conversion rate by autocatalytic growth. The second and third terms are derived from the law of mass action to represent the reaction rates in Eqs. (1) and (2). The final term on the right-hand side of Eq. (3) represents the degradation of TAF15 monomers. Also, in Eq. (3) the volume of the soma is

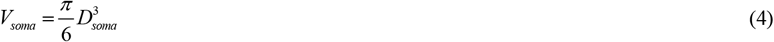

**Table 1.** Dependent variables used in the model.

| Symbol | Definition | Units |
| --- | --- | --- |
| $C_A$ | Molar concentration of TAF15 monomers | $\mu\text{M}$ |
| $C_B$ | Molar concentration of free TAF15 aggregates<br>(not deposited into inclusions) | $\mu\text{M}$ |
| $C_D$ | Molar concentration of TAF15 aggregates<br>deposited into TAF15 inclusions | $\mu\text{M}$ |
| $r_l$ | Radius of a TAF15 inclusion | $\mu\text{m}$ |

**Table 2.**
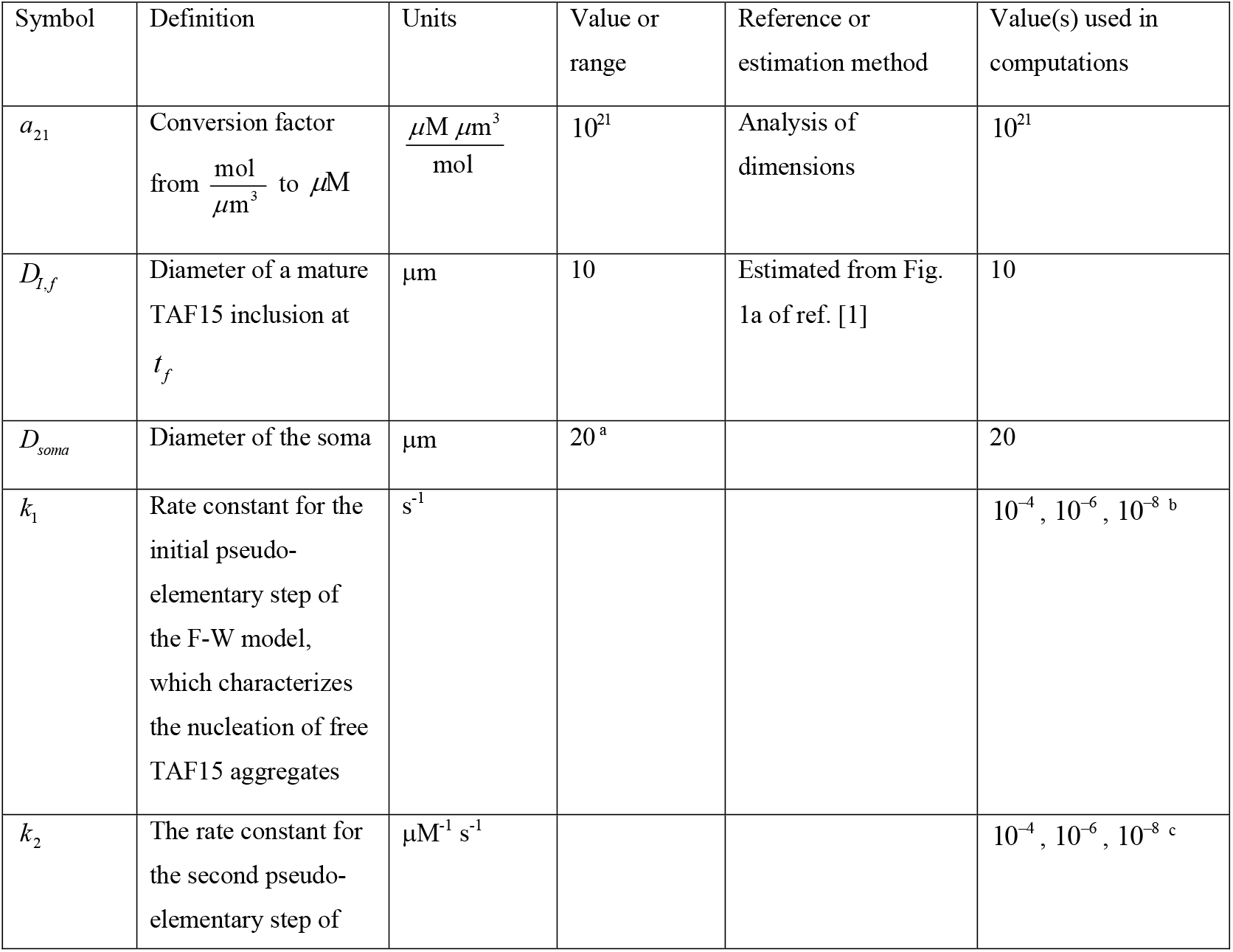
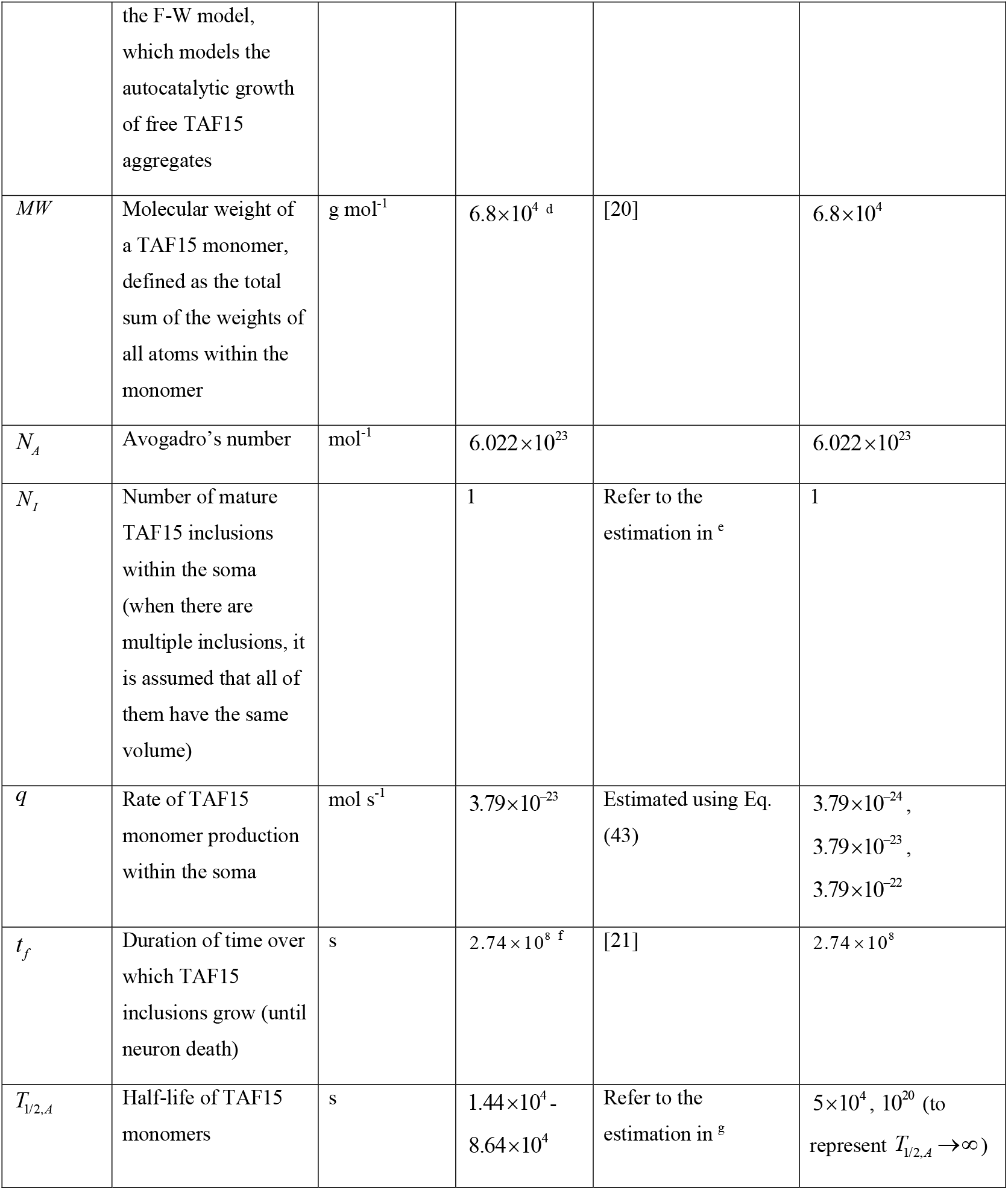
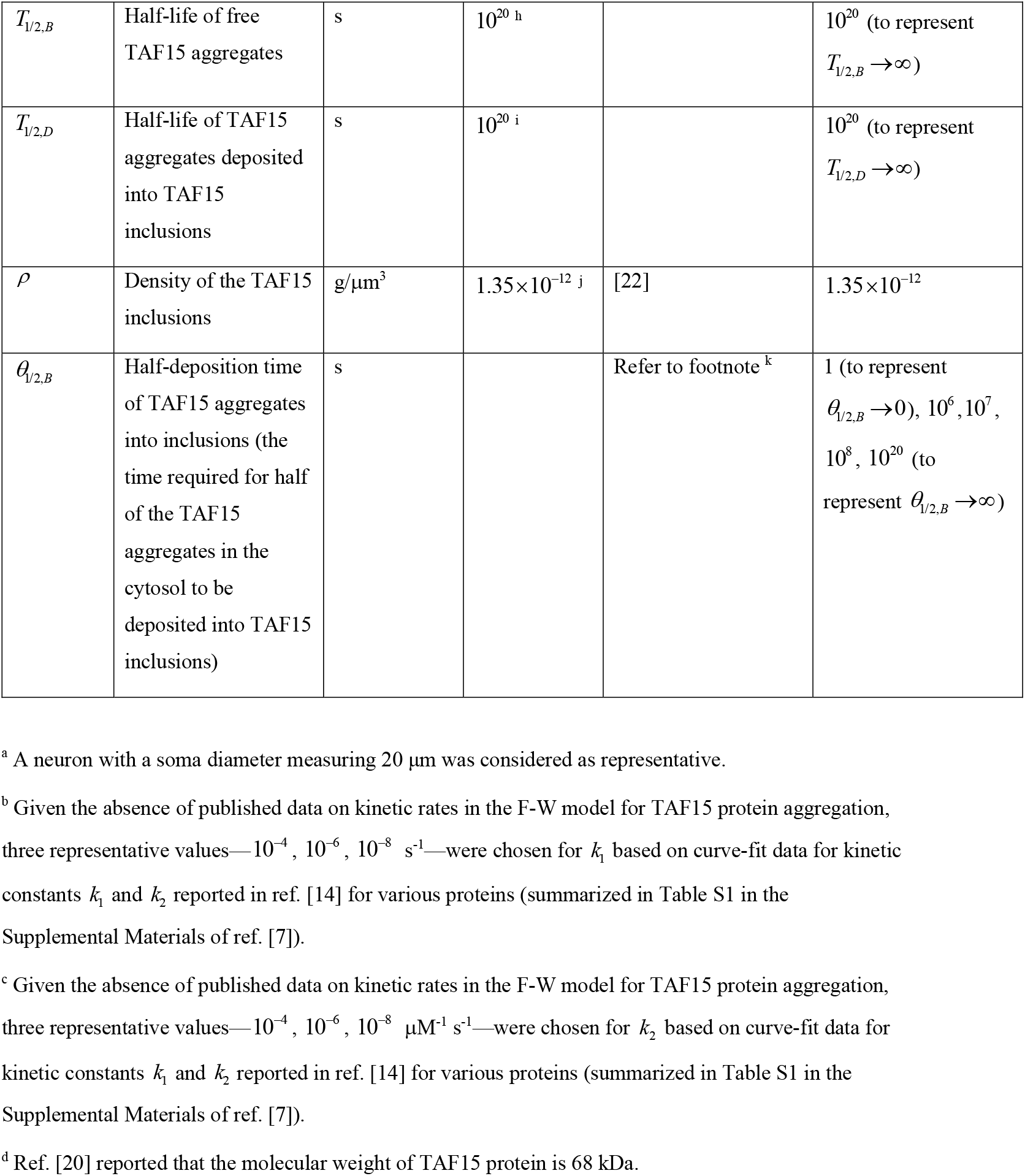
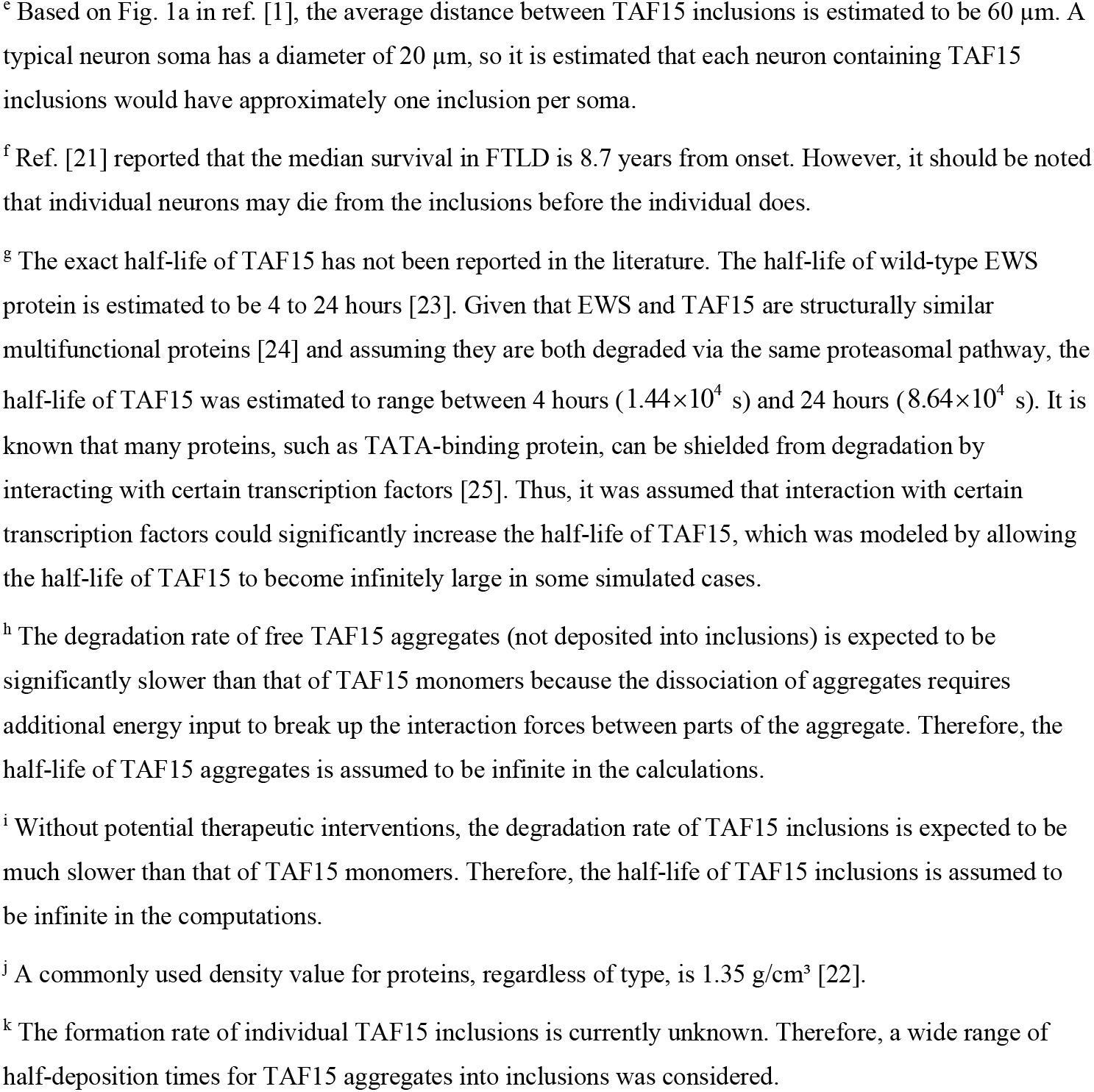
Parameters used in the model.

| Symbol | Definition | Units | Value or range | Reference or estimation method | Value(s) used in computations |
| --- | --- | --- | --- | --- | --- |
| $a_{21}$ | Conversion factor<br>from $\frac{\text{mol}}{\mu\text{m}^3}$ to $\mu\text{M}$ | $\frac{\mu\text{M } \mu\text{m}^3}{\text{mol}}$ | $10^{21}$ | Analysis of dimensions | $10^{21}$ |
| $D_{l,f}$ | Diameter of a mature TAF15 inclusion at $t_f$ | $\mu\text{m}$ | 10 | Estimated from Fig. 1a of ref. [1] | 10 |
| $D_{soma}$ | Diameter of the soma | $\mu\text{m}$ | $20^a$ | | 20 |
| $k_1$ | Rate constant for the initial pseudo-elementary step of the F-W model, which characterizes the nucleation of free TAF15 aggregates | $\text{s}^{-1}$ | | | $10^{-4}, 10^{-6}, 10^{-8}^b$ |
| $k_2$ | The rate constant for the second pseudo-elementary step of the F-W model,<br>which models the<br>autocatalytic growth<br>of free TAF15<br>aggregates | $\mu\text{M}^{-1} \text{s}^{-1}$ | | | $10^{-4}, 10^{-6}, 10^{-8}^c$ |
| $MW$ | Molecular weight of<br>a TAF15 monomer,<br>defined as the total<br>sum of the weights of<br>all atoms within the<br>monomer | $\text{g mol}^{-1}$ | $6.8 \times 10^4$ <sup>d</sup> | [20] | $6.8 \times 10^4$ |
| $N_A$ | Avogadro's number | $\text{mol}^{-1}$ | $6.022 \times 10^{23}$ | | $6.022 \times 10^{23}$ |
| $N_I$ | Number of mature<br>TAF15 inclusions<br>within the soma<br>(when there are<br>multiple inclusions, it<br>is assumed that all of<br>them have the same<br>volume) | | 1 | Refer to the<br>estimation in <sup>e</sup> | 1 |
| $q$ | Rate of TAF15<br>monomer production<br>within the soma | $\text{mol s}^{-1}$ | $3.79 \times 10^{-23}$ | Estimated using Eq.<br>(43) | $3.79 \times 10^{-24}$,<br>$3.79 \times 10^{-23}$,<br>$3.79 \times 10^{-22}$ |
| $t_f$ | Duration of time over<br>which TAF15<br>inclusions grow (until<br>neuron death) | s | $2.74 \times 10^8$ <sup>f</sup> | [21] | $2.74 \times 10^8$ |
| $T_{1/2,A}$ | Half-life of TAF15<br>monomers | s | $1.44 \times 10^4$ -<br>$8.64 \times 10^4$ | Refer to the<br>estimation in <sup>g</sup> | $5 \times 10^4$, $10^{20}$ (to<br>represent $T_{1/2,A} \rightarrow \infty$) |
| $T_{1/2,B}$ | Half-life of free TAF15 aggregates | s | $10^{20}$ <sup>h</sup> | | $10^{20}$ (to represent $T_{1/2,B} \rightarrow \infty$) |
| $T_{1/2,D}$ | Half-life of TAF15 aggregates deposited into TAF15 inclusions | s | $10^{20}$ <sup>i</sup> | | $10^{20}$ (to represent $T_{1/2,D} \rightarrow \infty$) |
| $\rho$ | Density of the TAF15 inclusions | $\text{g}/\mu\text{m}^3$ | $1.35 \times 10^{-12}$ <sup>j</sup> | [22] | $1.35 \times 10^{-12}$ |
| $\theta_{1/2,B}$ | Half-deposition time of TAF15 aggregates into inclusions (the time required for half of the TAF15 aggregates in the cytosol to be deposited into TAF15 inclusions) | s | | Refer to footnote <sup>k</sup> | 1 (to represent $\theta_{1/2,B} \rightarrow 0$), $10^6, 10^7, 10^8, 10^{20}$ (to represent $\theta_{1/2,B} \rightarrow \infty$) |

<sup>a</sup> A neuron with a soma diameter measuring 20 $\mu\text{m}$ was considered as representative.
<sup>b</sup> Given the absence of published data on kinetic rates in the F-W model for TAF15 protein aggregation, three representative values— $10^{-4}, 10^{-6}, 10^{-8} \text{ s}^{-1}$ —were chosen for $k_1$ based on curve-fit data for kinetic constants $k_1$ and $k_2$ reported in ref. [14] for various proteins (summarized in Table S1 in the Supplemental Materials of ref. [7]).
<sup>c</sup> Given the absence of published data on kinetic rates in the F-W model for TAF15 protein aggregation, three representative values— $10^{-4}, 10^{-6}, 10^{-8} \mu\text{M}^{-1} \text{ s}^{-1}$ —were chosen for $k_2$ based on curve-fit data for kinetic constants $k_1$ and $k_2$ reported in ref. [14] for various proteins (summarized in Table S1 in the Supplemental Materials of ref. [7]).
<sup>d</sup> Ref. [20] reported that the molecular weight of TAF15 protein is 68 kDa.
<sup>e</sup> Based on Fig. 1a in ref. [1], the average distance between TAF15 inclusions is estimated to be 60 $\mu\text{m}$ . A typical neuron soma has a diameter of 20 $\mu\text{m}$ , so it is estimated that each neuron containing TAF15 inclusions would have approximately one inclusion per soma.
<sup>f</sup> Ref. [21] reported that the median survival in FTLT is 8.7 years from onset. However, it should be noted that individual neurons may die from the inclusions before the individual does.
<sup>g</sup> The exact half-life of TAF15 has not been reported in the literature. The half-life of wild-type EWS protein is estimated to be 4 to 24 hours [23]. Given that EWS and TAF15 are structurally similar multifunctional proteins [24] and assuming they are both degraded via the same proteasomal pathway, the half-life of TAF15 was estimated to range between 4 hours ( $1.44 \times 10^4$ s) and 24 hours ( $8.64 \times 10^4$ s). It is known that many proteins, such as TATA-binding protein, can be shielded from degradation by interacting with certain transcription factors [25]. Thus, it was assumed that interaction with certain transcription factors could significantly increase the half-life of TAF15, which was modeled by allowing the half-life of TAF15 to become infinitely large in some simulated cases.
<sup>h</sup> The degradation rate of free TAF15 aggregates (not deposited into inclusions) is expected to be significantly slower than that of TAF15 monomers because the dissociation of aggregates requires additional energy input to break up the interaction forces between parts of the aggregate. Therefore, the half-life of TAF15 aggregates is assumed to be infinite in the calculations.
<sup>i</sup> Without potential therapeutic interventions, the degradation rate of TAF15 inclusions is expected to be much slower than that of TAF15 monomers. Therefore, the half-life of TAF15 inclusions is assumed to be infinite in the computations.
<sup>j</sup> A commonly used density value for proteins, regardless of type, is 1.35 g/cm<sup>3</sup> [22].
<sup>k</sup> The formation rate of individual TAF15 inclusions is currently unknown. Therefore, a wide range of half-deposition times for TAF15 aggregates into inclusions was considered.

By applying the principle of conservation to TAF15 aggregates within the soma, the following equation is obtained:

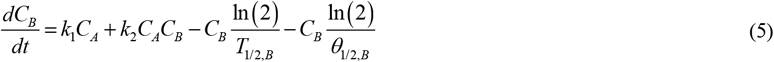

On the right side in Eq. (5), the first term accounts for the rate of aggregate production via nucleation, while the second term represents the rate of aggregate production through autocatalytic growth. These terms are of equal magnitude but opposite in sign to the second and third terms in Eq. (3), reflecting the F-W model’s principle that the aggregate production rate matches the monomer consumption rate. The third term in Eq. (5) models the degradation of TAF15 aggregates, which could become relevant with future therapies aimed at promoting TAF15 aggregate clearance to prevent or treat FTLD. The final term represents the deposition of TAF15 aggregates into TAF15 inclusions.

The formation of TAF15 inclusions from adhesive TAF15 fibrils is modeled by a method analogous to that used to simulate coagulation of colloidal suspensions [18]. It is assumed that free TAF15 aggregates (*B*) deposit into TAF15 inclusions, driving their growth. The concentration of these deposited aggregates is *C*_*D*_ . The conservation of deposited aggregates gives the following equation:

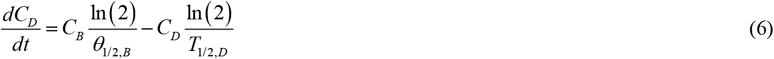

where *θ*_1/2,*B*_ is the half-deposition time of TAF15 aggregates into TAF15 inclusions, defined as the time required for half of the TAF15 aggregates in the control volume to be deposited into a TAF15 inclusion. The second term on the right-hand side of Eq. (6) represents the half-life of TAF15 aggregates within inclusions, accounting for potential future therapies aimed at clearing TAF15 inclusions [19].

The initial conditions are as follows:

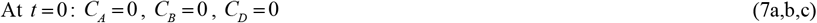

### 2.2. Dimensionless equations

The model expressed in Eqs. (3), (5), and (6) is reformulated in dimensionless form as follows:

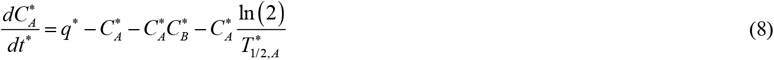

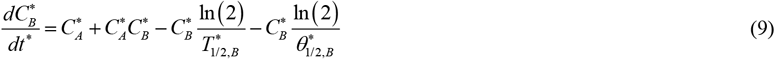

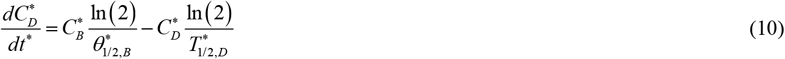

The initial conditions, given by Eq. (7), in dimensionless form can be expressed as follows:

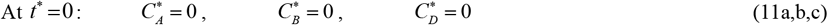

Table 3 defines the dimensionless independent variable, Table 4 summarizes the dimensionless dependent variables, and Table 5 provides the dimensionless parameters utilized in the model.

**Table 3.** Dimensionless independent variable used in the model.

| Symbol | Definition |
| --- | --- |
| $t^*$ | $t k_1$ |

**Table 4.**
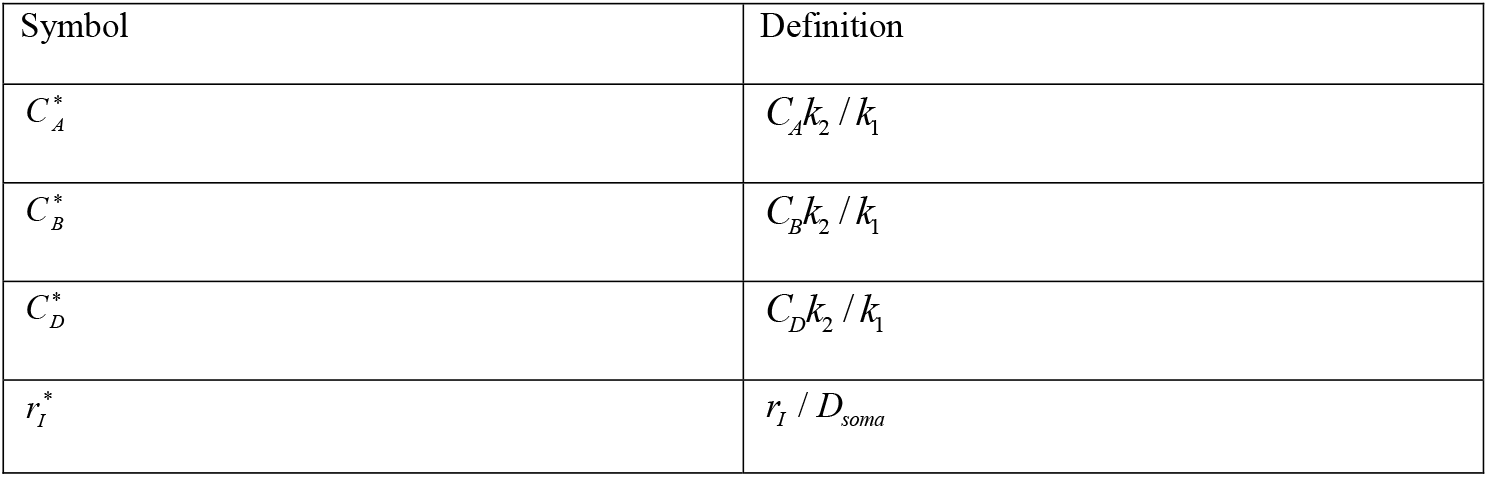
Dimensionless dependent variables used in the model.

**Table 5.** Dimensionless parameters used in the model.

| Symbol | Definition |
| --- | --- |
| $q^*$ | $\frac{q}{V_{soma}} a_{21} \frac{k_2}{(k_1)^2}$ |
| $t_f^*$ | $t_f k_1$ |
| $T_{1/2,A}^*$ | $T_{1/2,A} k_1$ |
| $T_{1/2,B}^*$ | $T_{1/2,B} k_1$ |
| $T_{1/2,D}^*$ | $T_{1/2,D} k_1$ |
| $\rho^*$ | $\frac{\rho}{MW} a_{21} \frac{k_2}{k_1}$ |
| $\theta_{1/2,B}^*$ | $\theta_{1/2,B} k_1$ |

### 2.3. The scenario where free TAF15 aggregates rapidly deposit into TAF15 inclusions, 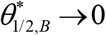

As 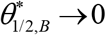, indicating immediate deposition of free TAF15 aggregates into inclusions, Eqs. (8)-(10) simplify to the following:

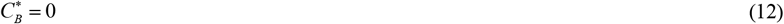

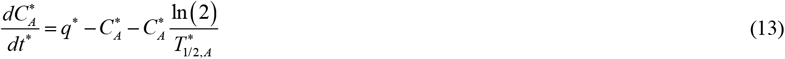

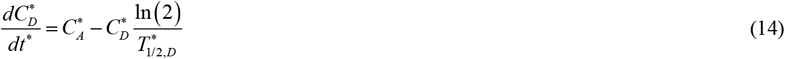

By adding Eqs. (13) and (14) for the case of 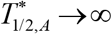 and 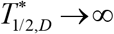 , the resulting equation is

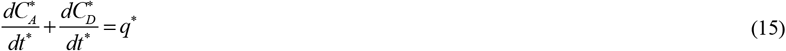

By integrating Eq. (15) with respect to time and applying the initial condition from Eq. (11a,c), the result is

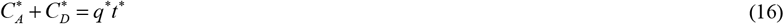

The increase in 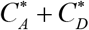 over time is due to the production of TAF15 monomers in the soma. By substituting 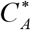 found from Eq. (16) into Eq. (14), the following is obtained:

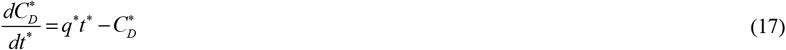

The exact solution of Eq. (17) with the initial condition given by Eq. (11c):

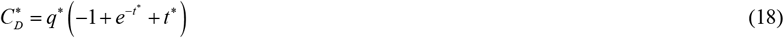

Rewriting Eq. (18) in terms of dimensional variables yields:

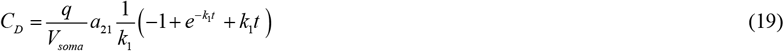

As *t*^∗^ → 0 , from Eq. (18) it follows that

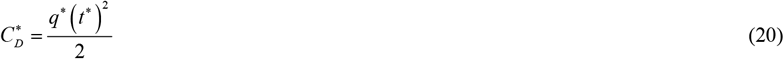

which shows that for small *t*^∗^ , the concentration of TAF15 aggregates deposited into inclusions increases proportionally to the square of time.

For large *t*^∗^ (as *t*^∗^ → ∞ ), from Eq. (18) it follows that

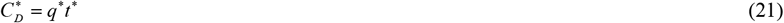

Returning to dimensional variables, Eq. (21) becomes

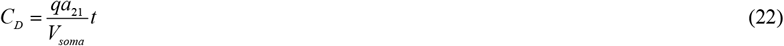

Then *C*_*A*_ can be calculated using Eqs. (16) and (18) as:

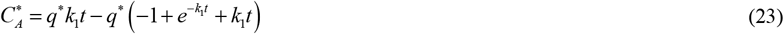

Rewriting Eq. (22) in terms of dimensional variables yields

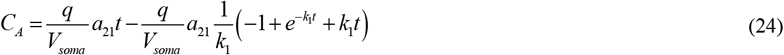

### 2.4. The scenario where the deposition rate of free TAF15 aggregates into TAF15 inclusions is slow, 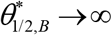

For the case of 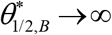, indicating no deposition of free TAF15 aggregates into inclusions, Eqs. (8)-(10) simplify to the following:

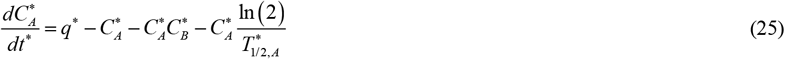

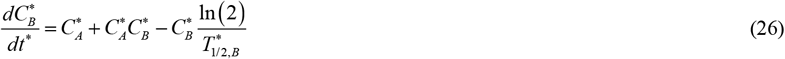

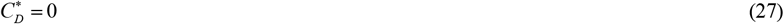

Eqs. (25) and (26) were solved using the initial conditions (11a,b) in dimensionless form, as detailed in ref. [8]. For completeness, analytical solutions for Eqs. (25) and (26) are provided in section S1 of the Supplementary Materials.

### 2.5. Simulation of TAF15 inclusion growth

The growth of TAF15 inclusions (refer to Fig. 1) is computed as follows. The number of TAF15 monomers incorporated into a single inclusion at any given time *t* can be described by the concentration of TAF15 monomers deposited into the inclusion, as detailed in ref. [26]:

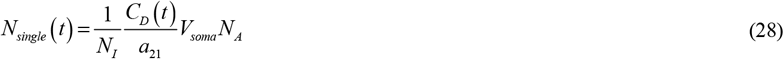

Here, *N*_*I*_ denotes the number of TAF15 inclusions in the soma, assuming that all inclusions have the same volume.

Following the method described in ref. [26] and presenting the number of monomers that was used to build a TAF15 inclusion through the volume of a single inclusion body, *V*_*single*_ , the following is obtained:

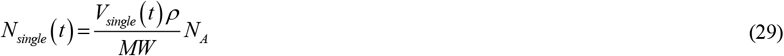

where *MW* represents the molecular weight of TAF15 monomers, calculated as the sum of the weights of all atoms within the monomer.

Equating the expressions on the right-hand sides of Eqs. (28) and (29), and solving for the volume of a single inclusion body, gives

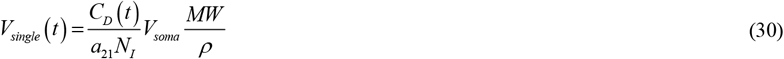

To derive the equation for the volume of a TAF15 inclusion at long times, Eq. (22) was substituted into Eq. (30), resulting in the following equation:

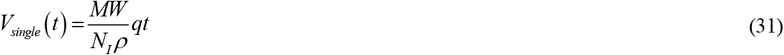

This equation suggests that the volume of a single inclusion body in the soma increases linearly with both the rate of TAF15 monomer generation and time.

TAF15 inclusions within the soma can vary in shape, from skein-like to rounded [27]. For simplicity, it was assumed that all inclusion bodies are spherical and of the same size. The volume of a spherical inclusion body is calculated by

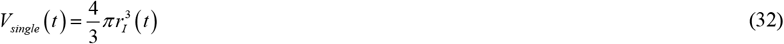

When equating the right-hand sides of Eqs. (30) and (32) and solving for the radius of the inclusion body, the following result emerges:

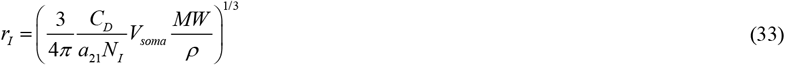

In its dimensionless form, Eq. (33) transforms into:

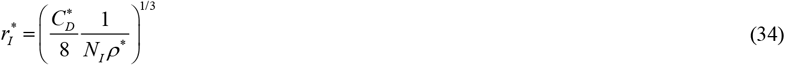

For rapid deposition rates of free TAF15 aggregates into TAF15 inclusions, that is 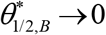, substituting Eq. (18) into Eq. (34) results in

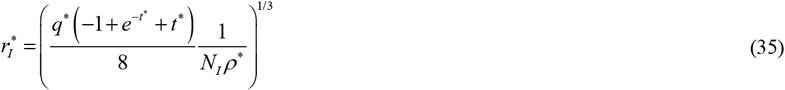

In dimensional variables Eq. (35) can be rewritten as:

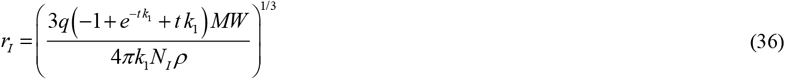

If *t*^∗^ → ∞ , Eq. (35) leads to

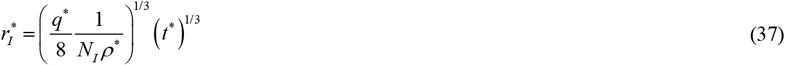

In dimensional variables, Eq. (37) can be expressed as:

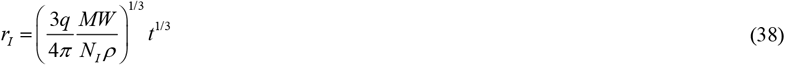

Eq. (38) presents the cube root hypothesis, initially formulated in ref. [12] for Lewy bodies. As *t*^∗^ → 0 , Eq. (35) simplifies to:

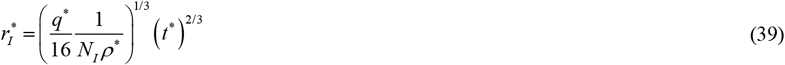

Eq. (35) can be used to estimate the rate at which TAF15 monomers are generated in the soma. If, after the growth period *t* _*f*_ , the radius of the inclusion body reaches the size *r*_*I* , *f*_ , then according to Eq. (35), *q* can be determined as:

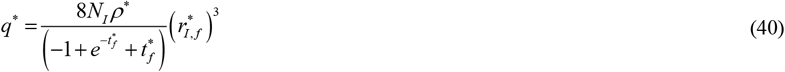

Returning to dimensional variables, Eq. (40) can be rewritten as:

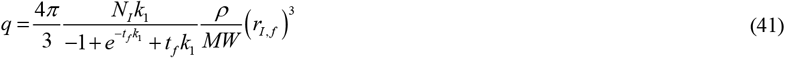

where *r*_*I, f*_ = *D*_*I, f*_ /2.

For 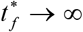, Eq. (40) simplifies to:

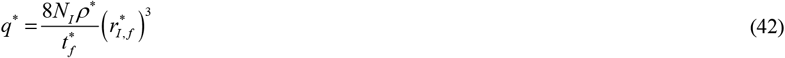

Rewriting Eq. (42) in dimensional variables yields:

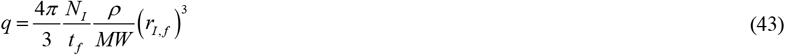

Eq. (43) is essential for estimating model parameters, as it allows the calculation of *q* when *r*_*I* , *f*_ and *t* are known. Using the Eq. (43), *q* was estimated to be 3.79×10^−23^ mol s^-1^, as reported in Table 2.

### 2.6. Sensitivity analysis

One of the goals of this study is to examine how sensitive the radius of mature TAF15 inclusions (at *t* = *t* _*f*_ ) is to various model parameters, such as the half-deposition time of TAF15 aggregates into inclusions, *θ*_1/2,*B*_ ; the rate constant describing the nucleation of TAF15 aggregates, *k*_1_ ; the rate constant describing the autocatalytic growth of TAF15 aggregates, *k*_2_ ; and the rate of TAF15 monomer production, *q* .

This required computing the local sensitivity coefficients, which denote the first-order partial derivatives of the observable (the radius of a mature TAF15 inclusion) with respect to parameters such as the half-deposition time of free TAF15 aggregates into inclusions. This procedure followed methods outlined in refs. [28-31]. For example, the sensitivity coefficient of *r*_*I* , *f*_ to *θ*_1/2,*B*_ was determined as follows:

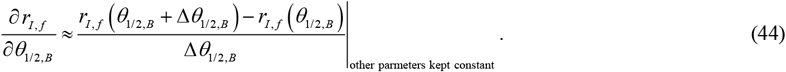

Here, Δ*θ*_1/2,*B*_ = 10^−3^*θ*_1/2,*B*_ is the step size. Various step sizes were used to assess the independence of the sensitivity coefficients from the step size.

The dimensionless relative sensitivity coefficients were calculated according to refs. [29,32], for instance,

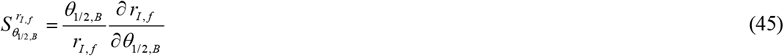

## 3. Results

Details of the numerical solution are provided in section S2 of the Supplementary Materials.

### 3.1. Validation of the numerical solution for the limiting cases of *θ*_1/2,*B*_ →0 and *θ*_1/2,*B*_ →∞

The numerical solution was validated by comparison to the analytical solution (given by Eqs. (12), (19), (24), and (36)) for the case of *T*_1/2,*A*_ →∞ , *T*_1/2,*B*_ →∞ , *T*_1/2,*D*_ →∞, and *θ*_1/2,*B*_ →0. Figs. S1 and S2 in the Supplementary Materials show a comparison for various values of *k*_1_, Figs. S3 and S4 illustrate comparisons for various values of *k*_2_, and Figs. S5 and S6 demonstrate comparisons for different values of *q* . All displayed cases show excellent agreement between the numerical and analytical solutions. The numerical solution was also validated by comparison to the analytical solution provided in Eqs. (S8), (S9), (27), and (S10) for the case of *T*_1/2,*A*_ →∞ , *T*_1/2,*B*_ →∞ , *T*_1/2,*D*_ →∞, and *θ*_1/2,*B*_ →∞. Figs. S7 and S8 in the Supplementary Materials present comparisons for various values of *k*_1_, Figs. S9 and S10 show comparisons for different values of *k*_2_, and Figs. S11 and S12 demonstrate comparisons for various values of *q* . All cases displayed show excellent agreement between the numerical and analytical solutions.

### 3.2. Analysis of model predictions for physiologically relevant parameter values

In the subsequent figures, physiologically relevant values are used for all parameters except for the one being varied. For the case of small *θ*_1/2,*B*_ all free TAF15 aggregates are immediately deposited into TAF15 inclusions. This results in a zero concentration of free TAF15 aggregates, *C*_*B*_ (see the line corresponding to *θ*_1/2,*B*_ = 1 s in Fig. 2b). The production of free TAF15 aggregates is driven by two mechanisms: nucleation and autocatalytic conversion (see Eq. (5)). Because for small values of *θ*_1/2,*B*_ there are no free TAF15 aggregates to catalyze the conversion of TAF15 monomers into aggregates, the production of TAF15 aggregates is solely driven by nucleation. Consequently, the growth rate of the concentration of TAF15 aggregates deposited into TAF15 inclusions, *C*_*D*_ , is slow, leading to a slow increase in the radius of the TAF15 inclusion (see the line corresponding to *θ*_1/2,*B*_ = 1 s in Fig. 3a and 3b).

**Fig. 2.**
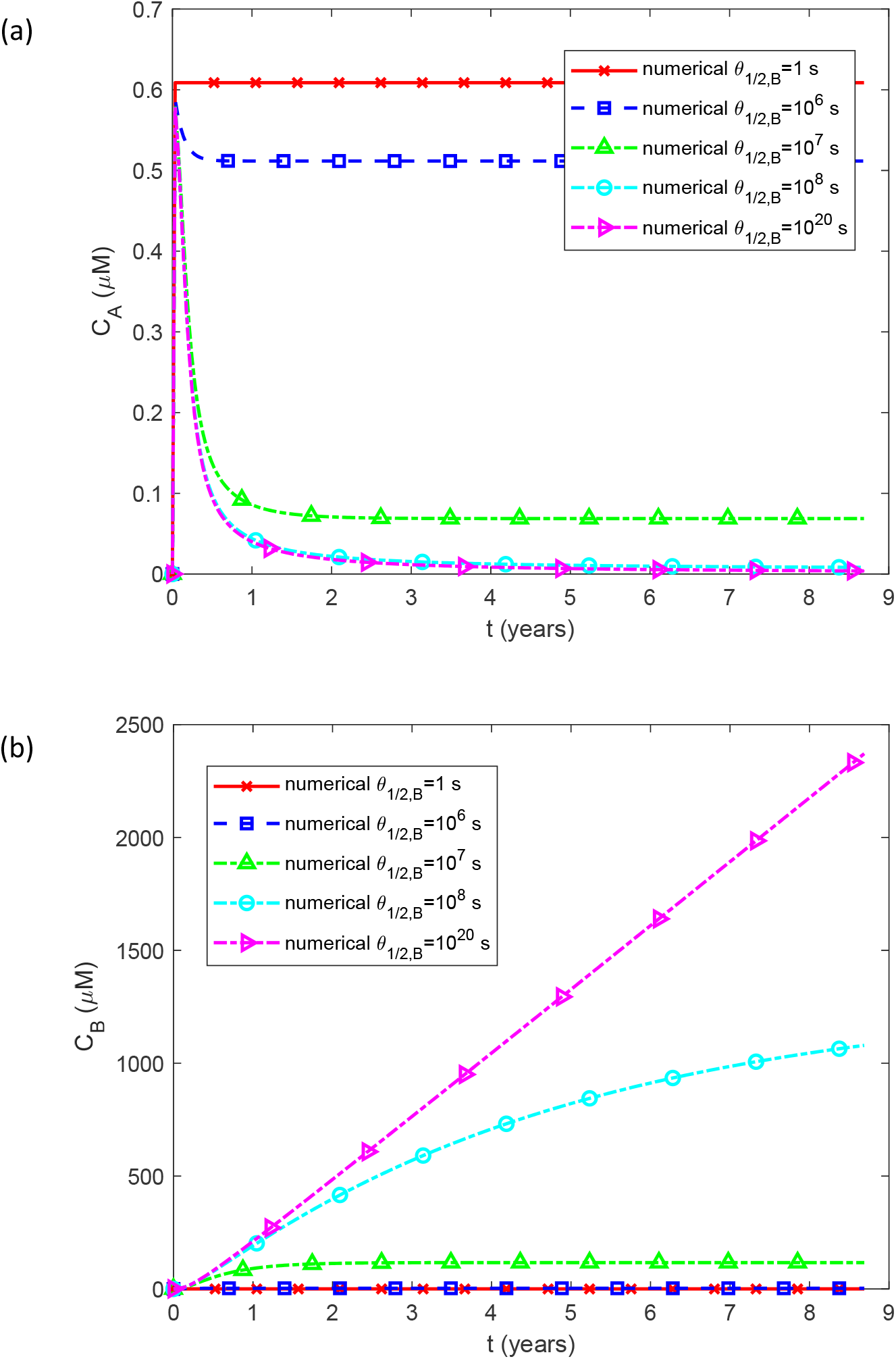
(a) The molar concentration of TAF15 monomers, *C*_*A*_ , plotted against time; and (b) the molar concentration of free TAF15 aggregates (not deposited into inclusions), *C*_*B*_ , plotted against time, for various values of *θ*_1/2,*B*_ . Except for *θ*_1/2,*B*_ , physiologically relevant parameter values are used: *k*_1_ = 10 ^−6^ s^-1^, *k*_2_ = 10^−6^ μM^-1^ s^-1^, *q* = 3.79×10^−23^ mol s^-1^, and *T*_1/2, *A*_ = 5×10^4^ s. Other parameters follow those specified in Table 2.

**Fig. 3.**
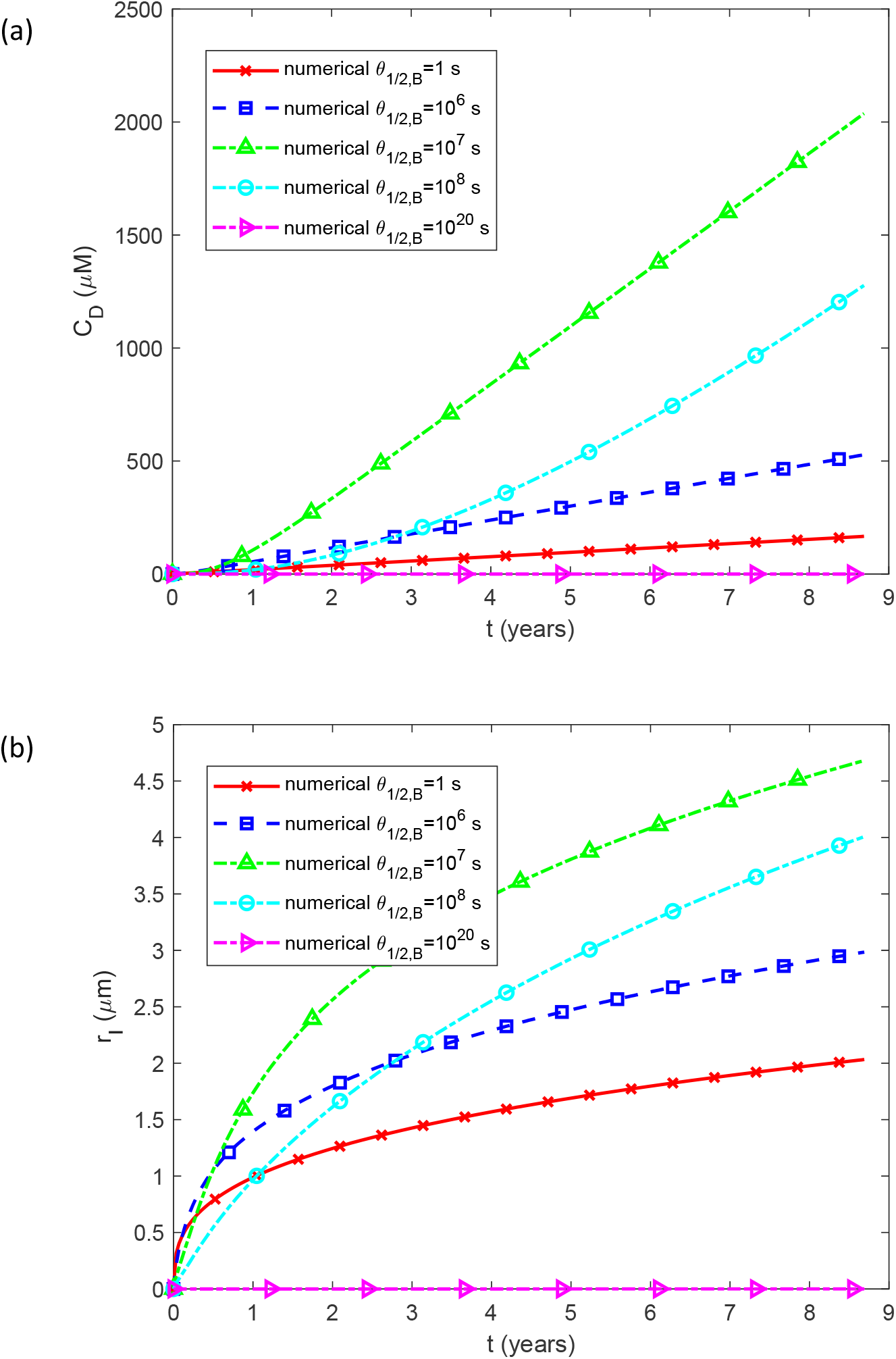
(a) The molar concentration of TAF15 aggregates deposited into TAF15 inclusions, *C*_*D*_ , plotted against time; and (b) the radius of a TAF15 inclusion, *r*_*I*_ , plotted against time, for various values of *θ*_1/2,*B*_ . Except for *θ*_1/2,*B*_ , physiologically relevant parameter values are used: *k*_1_ = 10 ^−6^ s^-1^, *k*_2_ = 10^−6^ μM^-1^ s^-1^, *q* = 3.79×10^−23^ mol s^-1^, and *T*_1/2, *A*_ = 5×10^4^ s. Other parameters are as specified in Table 2.

Conversely, if *θ*_1/2,*B*_ is large, free TAF15 aggregates do not deposit into inclusions. As *θ*_1/2,*B*_ →∞ , the rapid autocatalytic production of free TAF15 aggregates from monomers results in a linear increase in the free aggregate concentration with time (see the line corresponding to *θ*_1/2,*B*_ = 10^20^ s in Fig. 2b). This indicates that the growth of *C*_*B*_ in this scenario is limited by the rate of monomer production, *q* . The concentration of TAF15 monomers, *C*_*A*_ , quickly decreases over time because they are rapidly converted into free aggregates as *C*_B_ increases (see the line corresponding to *θ*_1/2,*B*_ = 10^20^ s in Fig. 2a).

For the case of *θ*_1/2,*B*_ →∞ , *C* _*D*_ and *r*_*I*_ are zero (see the lines corresponding to *θ*_1/2,*B*_ = 10^20^ in Fig. 3a and 3b). Consequently, the highest concentration of TAF15 aggregates deposited into TAF15 inclusions, *C*_*D*_ , and the largest radius of TAF15 inclusions, *r*_*I*_ , occur at an intermediate value of *θ*_1/2,*B*_ . This is confirmed in Fig. 3a and b, where the maximum values of *C* _*D*_ and *r* _*I*_ are observed for *θ*_1/2,*B*_ = 10^7^ s.

Increasing the kinetic constant *k*_1_ characterizing the nucleation rate of free TAF15 aggregates results in a more rapid rise in the concentration of free TAF15 aggregates, *C*_*B*_ (Fig. S13b). Consequently, there is a faster decline in the concentration of TAF15 monomers, *C*_*A*_ (Fig. S13a), due to their autocatalytic conversion into aggregates catalyzed by free aggregates, which occurs more rapidly with larger values of *C*_*B*_ . The accelerated increase in free aggregate concentration for higher *k*_1_ values leads to a quicker deposition of aggregates into inclusions, resulting in larger *C*_*D*_ (Fig. S14a) and faster growth of the inclusion radius, *r*_*I*_ (Fig. S14b).

The quantities *C*_*A*_ , *C*_*B*_ , *C*_*D*_ , and *r*_*I*_ exhibit greater sensitivity to variations in *k*_2_ compared to variations in *k*_1._ When *k*_2_ is small (*k*_2_ = 10^−8^ μM^-1^ s^-1^), the concentration of free aggregates, *C* _*B*_ (Fig. S15b), remains low. This results in minimal autocatalytic conversion of monomers into aggregates, leading to a very slow decrease in the monomer concentration, *C*_*A*_ (Fig. S15a). Conversely, with a large value of *k*_2_ (*k*_2_= 10^−4^ μM^-1^ s^-1^) the concentration of *C* _*B*_ increases rapidly (Fig. S15b), while *C* _*A*_ drops to zero almost immediately due to the high rate of autocatalytic conversion of monomers into aggregates (Fig. S15a).

The concentration of aggregates deposited into inclusions, *C*_*D*_ , changes almost linearly with time (Fig. S16a), suggesting that the growth of inclusions in this case is primarily driven by the constant rate of TAF15 monomer production (almost all monomers are converted into deposited aggregates, with only a few being degraded). The dependence of the radius of TAF15 inclusions, *r*_*I*_ , on time (Fig. S16b) is consistent with the cube root hypothesis expressed by Eq. (38).

The concentration of free TAF15 aggregates, *C*_*B*_ , increases with the production rate of TAF15 monomers, *q* (Fig. S17b). The TAF15 monomer concentration, *C*_*A*_ , is lowest when the TAF15 monomer production rate, *q* , is smallest (Fig. S17a). The concentration of aggregates deposited into inclusions, *C*_*D*_ , depends linearly on *q* and *t*, consistent with Eq. (22). The inclusion radius, *r* _*I*_ , is proportional to *q*^1/3^ and *t*^1/3^ , in agreement with Eq. (38) (see Fig. S18). The increase in the inclusion radius with a higher TAF15 monomer production rate aligns with findings reported in ref. [33], which indicated that increased TAF15 expression is detrimental to working memory in aged mice. Ref. [33] hypothesized that this could be due to the formation of amyloid filaments, attributed to TAF15, which may be harmful to the actin skeleton in dendritic spines.

To better understand the effect of *θ*_1/2,*B*_ , Figs. 4 and 5 plot the values of *C*_*A*_ , *C*_*B*_ , *C*_*D*_ , and *r*_*I*_ for various values of *k*_1_ after 8.7 years of growth (*t* = *t* _*f*_ ) vs *θ*_1/2,*B*_ . As *θ*_1/2,*B*_ → 0 , free TAF15 aggregates immediately deposit into TAF15 inclusions, resulting in a zero concentration of free TAF15 aggregates, *C*_*B, f*_ (Fig. 4b). Since the absence of free TAF15 aggregates leads to a zero rate of autocatalytic conversion of monomers into aggregates, the concentration of TAF15 monomers, *C*_*A, f*_ , is highest at small values of *θ*_1/2,*B*_ . The opposite situation occurs as *θ*_1/2,*B*_ →∞ . In this case, free TAF15 aggregates do not deposit into TAF15 inclusions, resulting in the highest concentration of free TAF15 aggregates, *C*_*B, f*_ (Fig. 4b). This high concentration of free TAF15 aggregates leads to a high rate of autocatalytic conversion of TAF15 monomers into free aggregates, resulting in an almost zero concentration of TAF15 monomers, *C*_*A, f*_ (Fig. 4a).

**Fig. 4.**
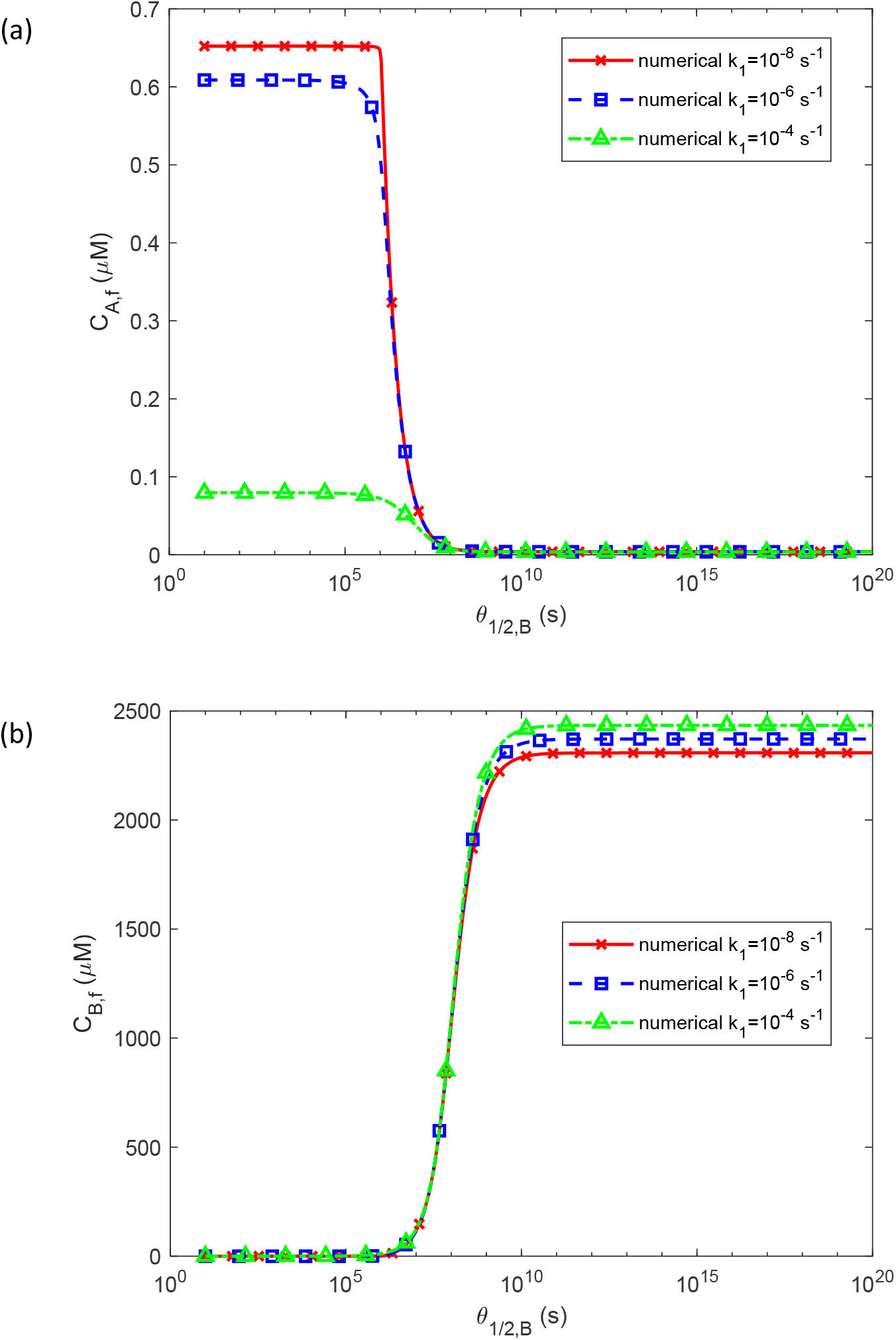
(a) The molar concentration of TAF15 monomers at *t* _*f*_ , *C*_*A, f*_ , plotted against *θ*_1/2,*B*_ ; and (b) the molar concentration of free TAF15 aggregates (not deposited into inclusions) at *t* _*f*_ , *C*_*B* , *f*_ , plotted against *θ*_1/2,*B*_ , for various values of *k*_1_ . Except for *θ*_1/2,*B*_ and *k*_1_ , physiologically relevant parameter values are used: *k*_2_ = 10^−6^ μM^-1^ s^-1^, *q* = 3.79×10^−23^ mol s^-1^, and *T*_1/2, *A*_ = 5×10^4^ s. Other parameters are as specified in Table 2.

**Fig. 5.**
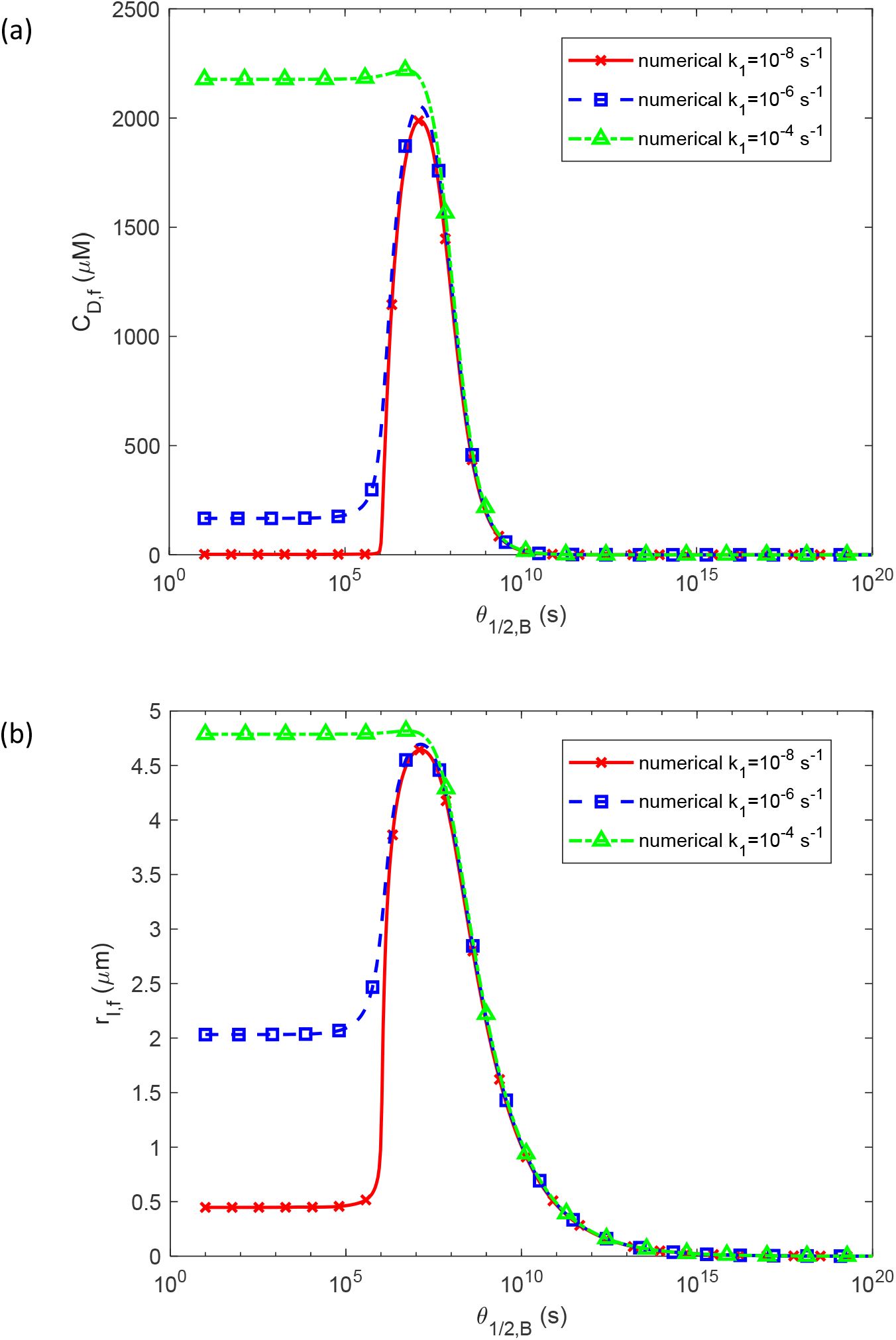
(a) The molar concentration of TAF15 aggregates deposited into TAF15 inclusions at *t* _*f*_ , *C*_*D* , *f*_ , plotted against *θ*_1/2,*B*_ ; and (b) the radius of a TAF15 inclusion at *t* _*f*_ , *r*_*I* , *f*_ , plotted against *θ*_1/2,*B*_ , for various values of *k*_1_ . Except for *θ*_1/2,*B*_ and *k*_1_ , physiologically relevant parameter values are used: *k*_2_ = 10^−6^ μM^-1^ s^-1^, *q* = 3.79×10^−23^ mol s^-1^, and *T*_1/2, *A*_ = 5×10^4^ s. Other parameter values are as specified in Table 2.

Thus, the lack of autocatalytic conversion and deposition leads to a small concentration of TAF15 aggregates deposited into inclusions. For this reason, the highest concentration of TAF15 aggregates deposited into inclusions, *C*_*D, f*_ , and the largest radius of TAF15 inclusions, *r*_*I* , *f*_ , occur at an intermediate value of *θ*_1/2,*B*_ , approximately 10^7^ s, as shown in Figs. 5a and 5b.

It is notable that for large values of *k*_1_ (10^−4^ s^-1^), *C* _*D, f*_ and *r* _*I* , *f*_ remain large even for small values of *θ*_1/2,*B*_ (Fig. 5). This occurs because as *θ*_1/2,*B*_ → 0 the concentration of free TAF15 aggregates is negligibly small (Fig. 4b), resulting in a negligible autocatalytic conversion rate from TAF15 monomers to aggregates. However, the high value of *k*_1_ compensates for the lack of autocatalytic conversion through a high nucleation rate.

The dependence of *C*_*B, f*_ on *θ*_1/2,*B*_ for various *k*_2_ (Fig. 6b) is similar to that for various *k*_1_ (Fig. 4b), but the effect of *k*_2_ variation is more pronounced, particularly for large values of *θ*_1/2,*B*_ (Fig. 6b). Since the concentration of TAF15 monomers depends on the rate of their conversion into free aggregates, the monomer concentration, *C*_*A, f*_ , strongly depends on *k*_2_ , especially for large values of *θ*_1/2,*B*_ (Fig. 6a).

**Fig. 6.**
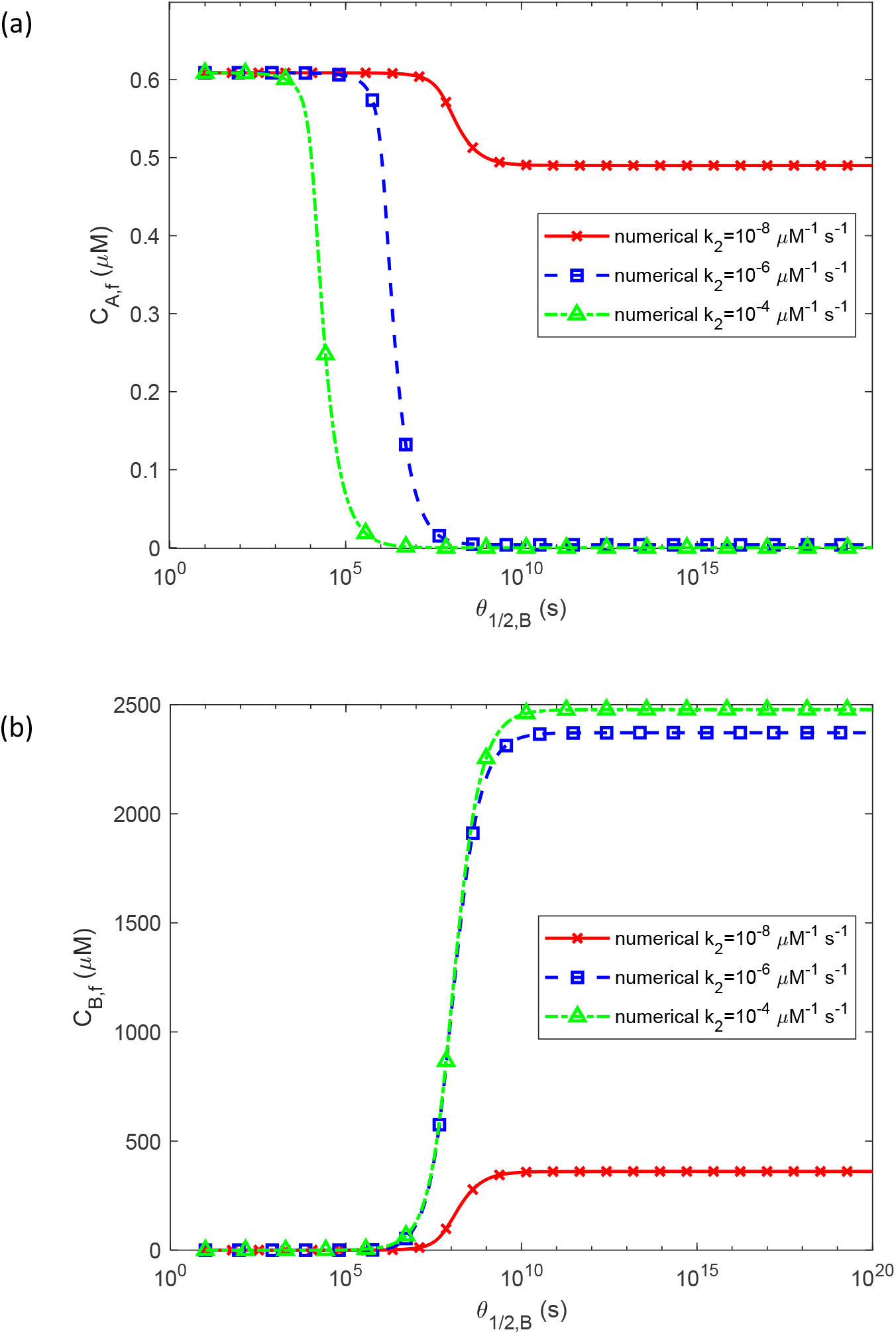
(a) The molar concentration of TAF15 monomers at *t* _*f*_ , *C*_*A, f*_ , plotted against *θ*_1/2,*B*_ ; and (b) the molar concentration of free TAF15 aggregates (not deposited into inclusions) at *t* _*f*_ , *C*_*B* , *f*_ , plotted against *θ*_1/2,*B*_ , for various values of *k*_2_ . Except for *θ*_1/2,*B*_ and *k*_2_ , physiologically relevant parameter values are used: *k*_1_ = 10 ^−6^ s^-1^, *q* = 3.79×10^−23^ mol s^-1^, and *T*_1/2, *A*_ = 5×10^4^ s. Other parameter values are as specified in Table 2.

For various values of *k*_2_ , the range of *θ*_1/2,*B*_ at which *C*_*D, f*_ and *r*_*I* , *f*_ take large values is broader than for various values of *k*_1_ (compare Fig. 7 to Fig. 5).

**Fig. 7.**
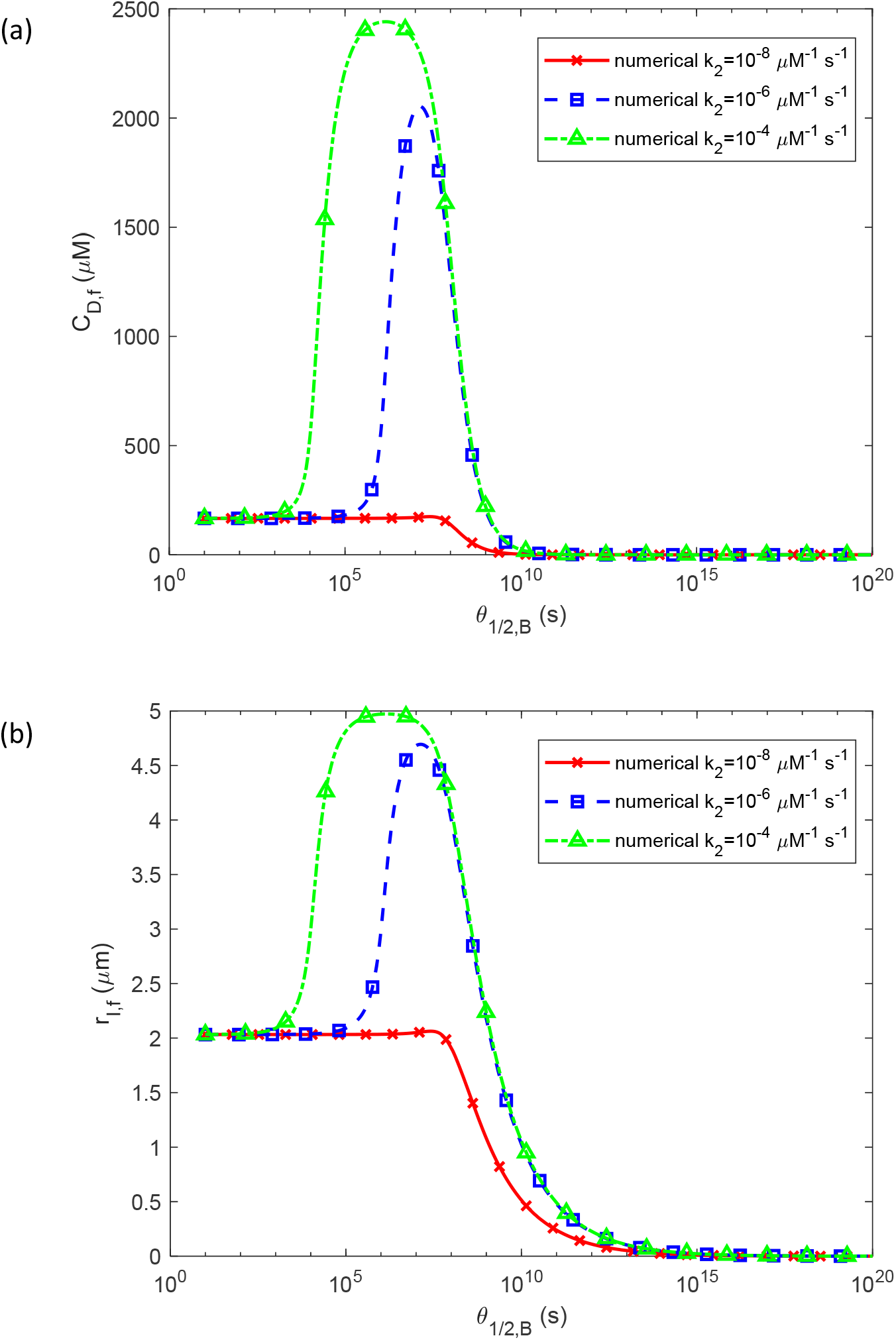
(a) The molar concentration of TAF15 aggregates deposited into TAF15 inclusions at *t* _*f*_ , *C*_*D* , *f*_ , plotted against *θ*_1/2,*B*_ ; and (b) the radius of a TAF15 inclusion at *t* _*f*_ , *r*_*I* , *f*_ , plotted against *θ*_1/2,*B*_ , for various values of *k*_2_ . Except for *θ*_1/2,*B*_ and *k*_2_ , physiologically relevant parameter values are used: *k*_1_ = 10 ^−6^ s^-1^, *q* = 3.79×10^−23^ mol s^-1^, and *T*_1/2, *A*_ = 5×10^4^ s. Other parameter values are as specified in Table 2.

The dependence of *C*_*B, f*_ on *θ*_1/2,*B*_ for various rates of TAF15 monomer production in the soma, *q* (Fig. 8b), is similar to that for various *k*_1_ (Fig. 4b), but variation in *q* affects *C*_*B, f*_ more strongly for large values of *θ*_1/2,*B*_ (Fig. 8b). This is because free TAF15 aggregates are directly produced by conversion from TAF15 monomers and changing the rate of monomer production by a factor of 10 (Fig. 8) significantly impacts the number of TAF15 monomers available for conversion. Similarly, variation in *q* strongly affects the concentration of TAF15 monomers, *C*_*A, f*_ , for small values of *θ*_1/2,*B*_ (Fig. 8a), when autocatalytic conversion from monomers into free aggregates is slow due to the low concentration of free TAF15 aggregates (Fig. 8b).

**Fig. 8.**
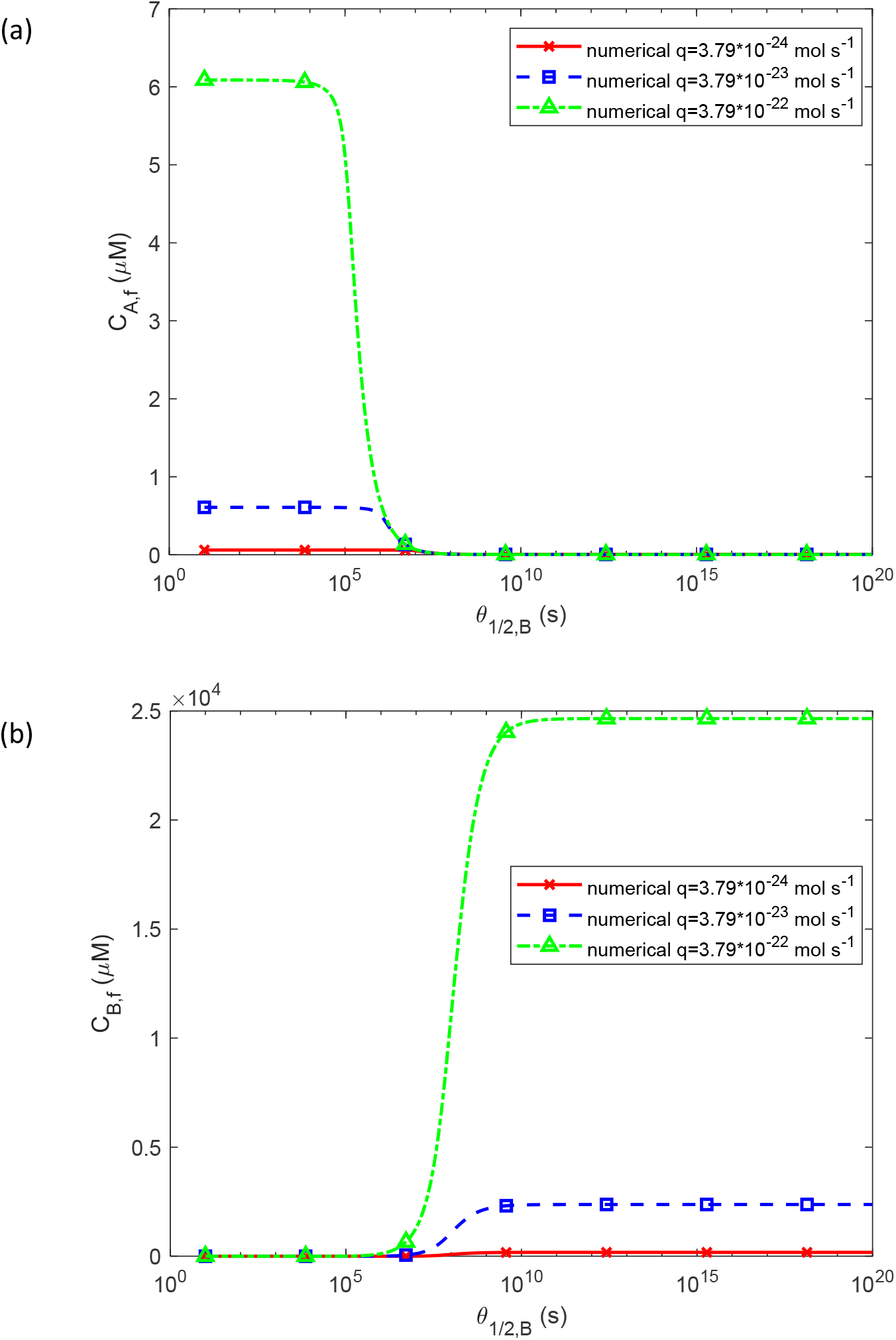
(a) The molar concentration of TAF15 monomers at *t* _*f*_ , *C*_*A, f*_ , plotted against *θ*_1/2,*B*_ ; and (b) the molar concentration of free TAF15 aggregates (not deposited into inclusions) at *t* _*f*_ , *C*_*B* , *f*_ , plotted against *θ*_1/2,*B*_ , for various values of *q* . Except for *θ*_1/2,*B*_ and *q* , physiologically relevant parameter values are used: *k*_1_ = 10 ^−6^ s^-1^, *k*_2_ = 10^−6^ μM^-1^ s^-1^, and *T*_1/2, *A*_ = 5×10^4^ s. Other parameter values are as specified in Table 2.

The value of *q* significantly influences the magnitude of the peaks of *C*_*D, f*_ and *r*_*I* , *f*_ , which occur at approximately *θ*_1/2,*B*_ = 10^7^ s (Fig. 9). A decrease in *q* causes a slight shift in the value of *θ*_1/2,*B*_ at which the peaks occur, displacing it to larger values (Fig. 9).

**Fig. 9.**
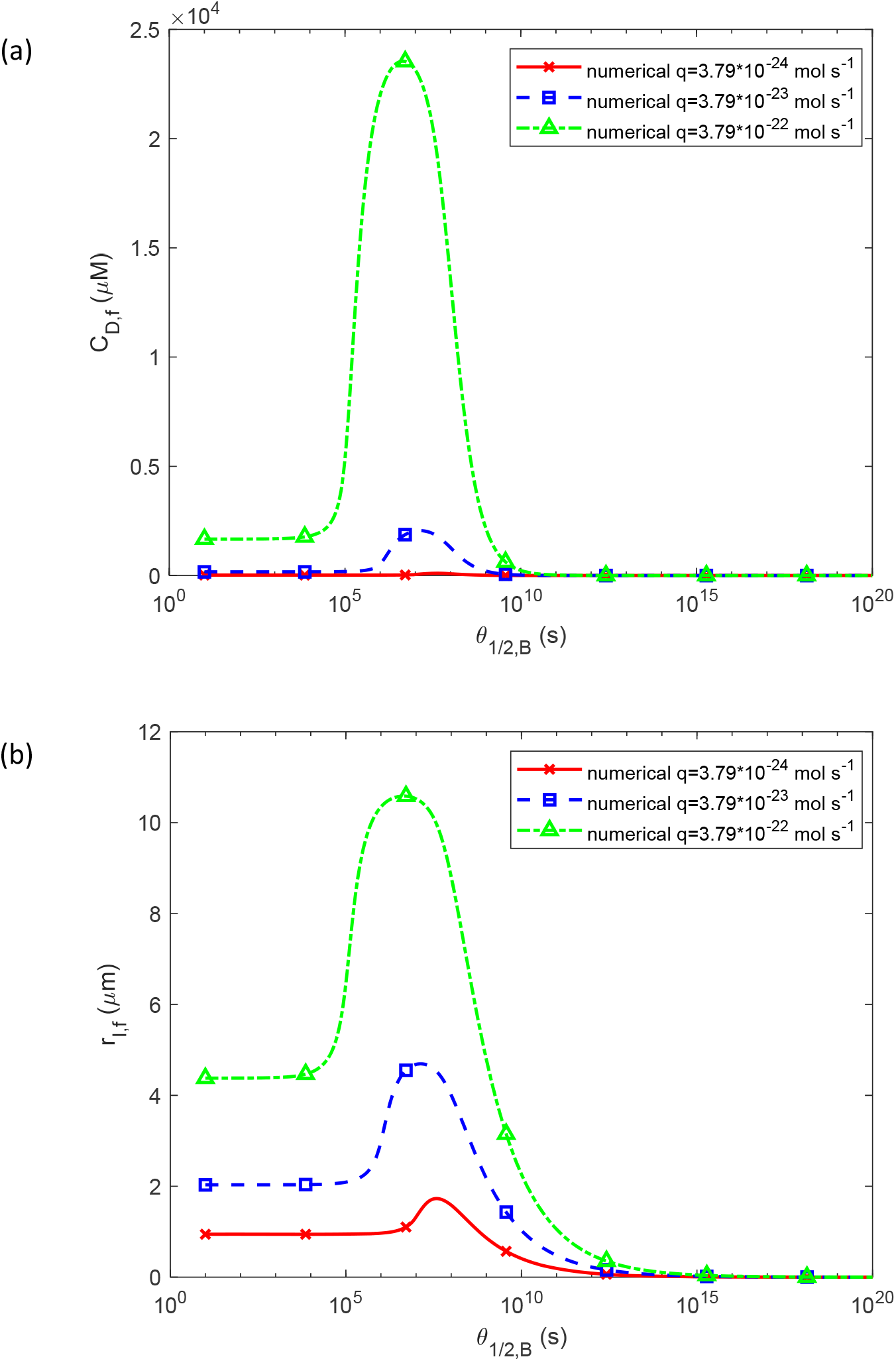
(a) The molar concentration of TAF15 aggregates deposited into TAF15 inclusions at *t* _*f*_ , *C*_*D* , *f*_ , plotted against *θ*_1/2,*B*_ ; and (b) the radius of a TAF15 inclusion at *t* _*f*_ , *r*_*I* , *f*_ , plotted against *θ*_1/2,*B*_ , for various values of *q* . Except for *θ*_1/2,*B*_ and *q* , physiologically relevant parameter values are used: *k*_1_ = 10 ^−6^ s^-1^, *k*_2_ = 10^−6^ μM^-1^ s^-1^, and *T*_1/2, *A*_ = 5×10^4^ s. Other parameter values are as specified in Table 2.

### 3.3. Analysis of the sensitivity of the radius of a mature TAF15 inclusion to various model parameters

The sensitivity of *r*_*I* , *f*_ to *θ*_1/2,*B*_, 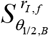 , peaks at approximately *θ*_1/2,*B*_ = 10^7^ s, independent of the value of *k*_1_ (Fig. 10a). When the value of *k*_2_ is changed, the peaks of 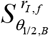 occur at different values of *θ*_1/2,*B*_ , depending on *k*_2_ . Interestingly, for the curve corresponding to *k*_2_ = 10^−8^ μM^-1^ s^-1^, the peak disappears completely (Fig. 10b). Similarly, when the rate of monomer production, *q* , is varied, the peaks of 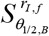 occur at different values of *θ*_1/2,*B*_ , depending on *q* . The peaks occur for all three values of *q* used in the computations but at different values of *θ*_1/2,*B*_ (Fig. 10c).

**Fig. 10.**
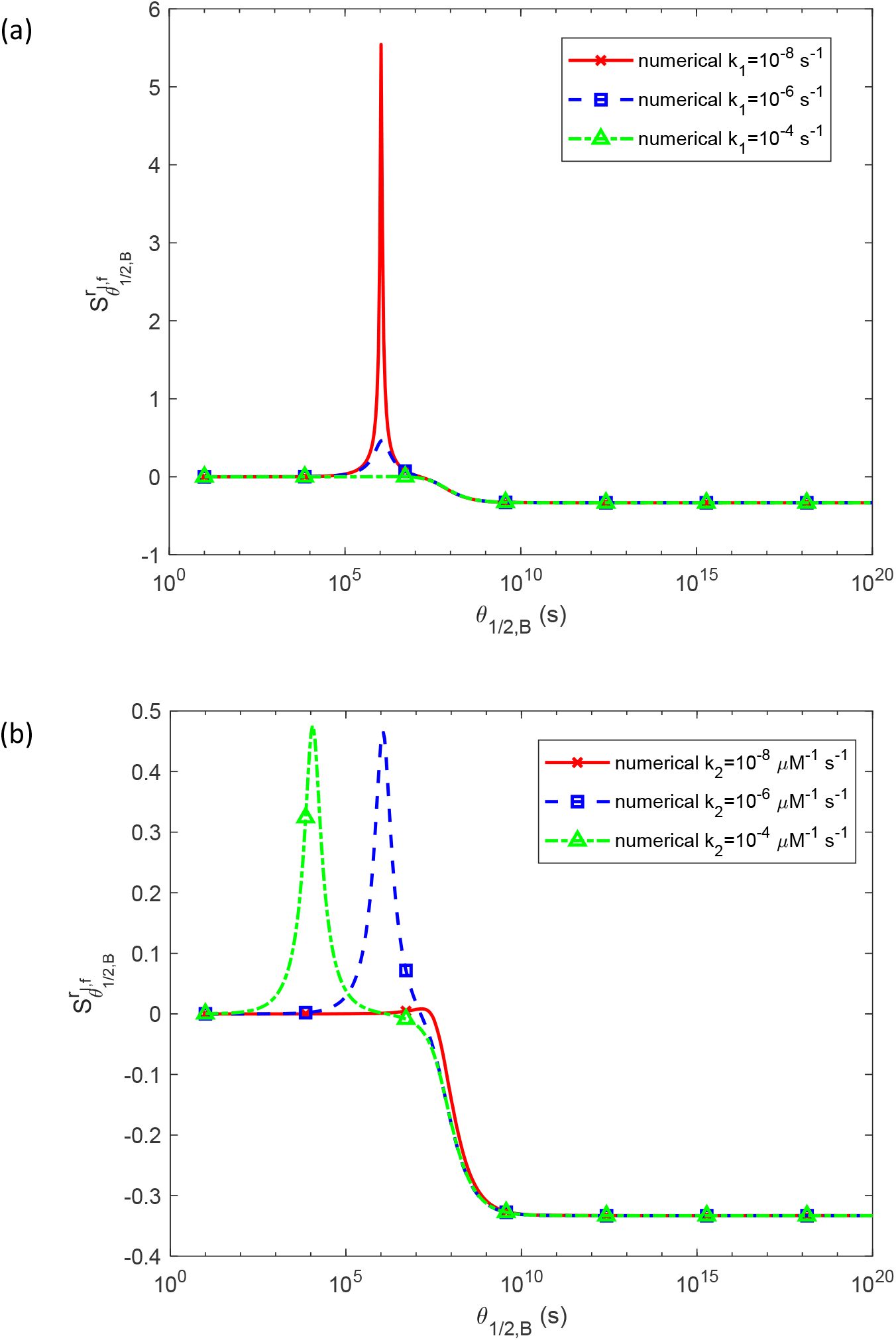
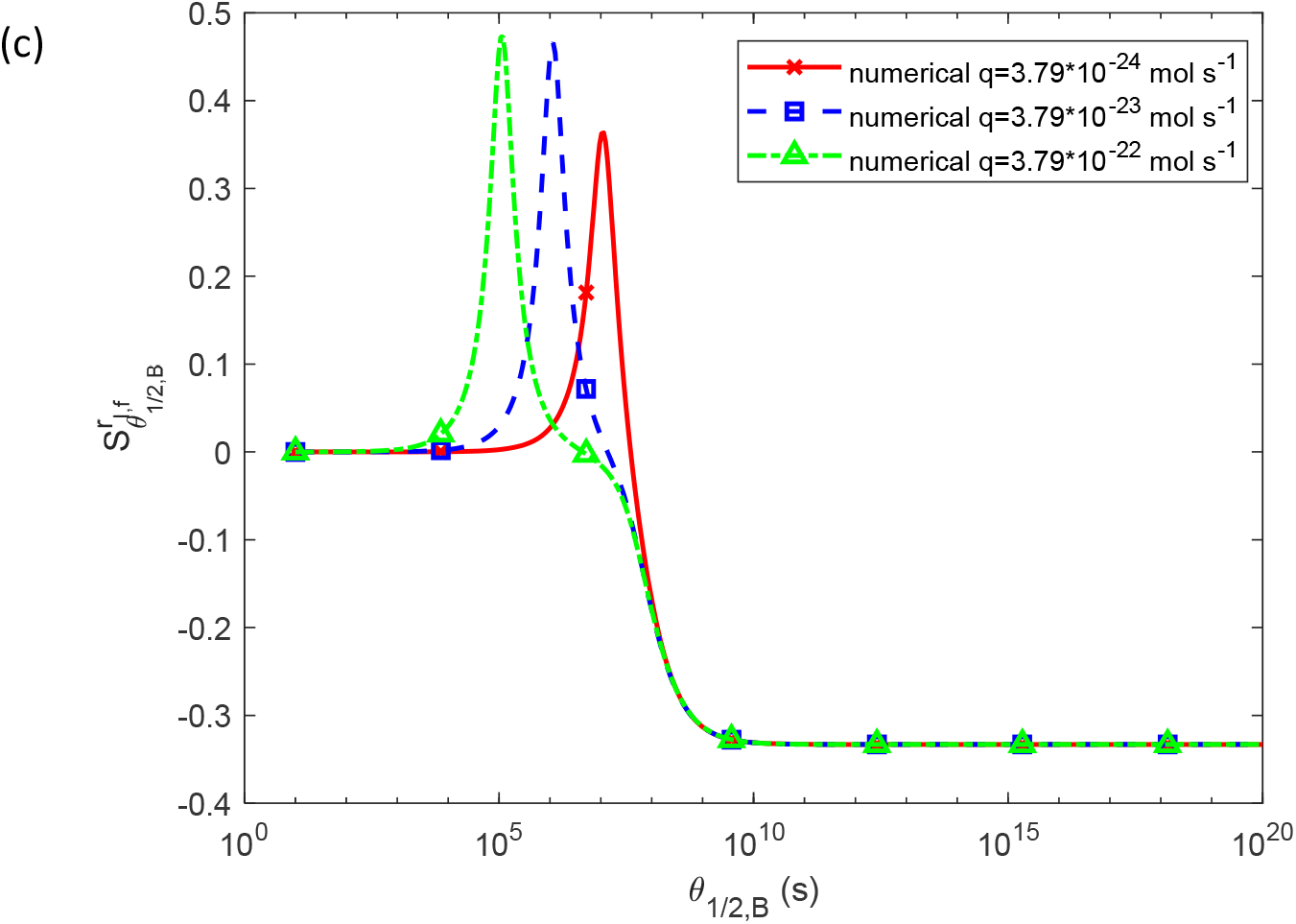
(a) The dimensionless sensitivity of the radius of a TAF15 inclusion at *t* _*f*_ , *r*_*I* , *f*_ , to the half-deposition time of TAF15 aggregates into inclusions, *θ*_1/2,*B*_,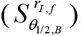 for various values of *k*_1_ . Except for *θ*_1/2,*B*_ and *k*_2_ , physiologically relevant parameter values are used: *k*_2_ = 10^−6^ μM^-1^ s^-1^, *q* = 3.79×10^−23^ mol s^-1^, and *T*_1/2, *A*_ = 5×10^4^ s. Other parameter values are as specified in Table 2. (b) The dimensionless sensitivity of *r*_*If*_ to *θ*_1/2,*B*_ 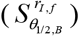 for various values of *k*_2_ . Except for *θ*_1/2,*B*_ and *k*_2_ , physiologically relevant parameter values are used: *k*_1_ = 10 ^−6^ s^-1^, *q* = 3.79×10^−23^ mol s^-1^, and *T*_1/2, *A*_ = 5×10^4^ s. Other parameter values are as specified in Table 2. (c) The dimensionless sensitivity of *r*_*If*_ to *θ*_1/2,*B*_ 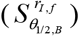 for various values of *q* . Except for *θ*_1/2,*B*_ and *q* , physiologically relevant parameter values are used: *k*_1_ = 10 ^−6^ s^-1^, *k*_2_ = 10^−6^ μM^-1^ s^-1^, and *T*_1/2, *A*_ = 5×10^4^ s. Other parameter values are as specified in Table 2.

The sensitivity of *r*_*I* , *f*_ to *k*_1_ , 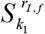 , drops to almost zero at approximately *θ*_1/2,*B*_ = 10^7^ s. This drop occurs because for large values of *θ*_1/2,*B*_ free TAF15 aggregates do not deposit into inclusions. Consequently, the inclusions do not grow, and their radii become independent of *k*_1_ . The value of *θ*_1/2,*B*_ at which this drop occurs is independent of *k*_1_ as well (Fig. S19a).

For small *θ*_1/2,*B*_ and value *k*_1_ (*k*_1_ = 10^−8^ s^-1^), 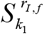 equals approximately 0.33 (Fig. 19a). This is consistent with the approximate solution obtained for *θ*_1/2,*B*_ → 0 . Indeed, according to Eq. (36),

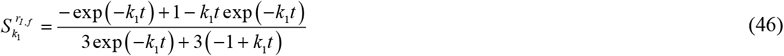

If *k*_1_ = 0 and *t* →∞ , then

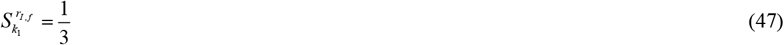

The sensitivity of *r* _*I* , *f*_ to *k*_2_ , 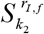 exhibits peaks at varying values of *θ*_1/2,*B*_ , dependent on *k*_2_ . Interestingly, when plotted for *k*_2_ = 10^−8^ μM^-1^ s^-1^, the peak disappears entirely, showing a monotonic increase of 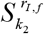 with *θ*_1/2,*B*_ (Fig. S19b). Peaks in the sensitivity of *r* _*I* , *f*_ to the rate of monomer production *q* , 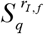 , also vary with *q* , appearing across different values of *θ*_1/2,*B*_ used in the study (Fig. S19c). For small values of *θ*_1/2,*B*_ , 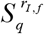 equals approximately 0.33, consistent with the result that follows from Eq. (36):

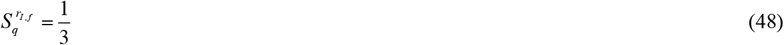

## 4. Discussion, limitations of the model, and future directions

A mathematical model simulating the formation and growth of TAF15 inclusions is developed. The model encompasses the production of TAF15 monomers, the nucleation and autocatalytic growth of free TAF15 aggregates, and the deposition of these aggregates into TAF15 inclusions. The numerical solution of the model’s equations is validated by comparing with analytical solutions obtained for two limiting cases, very fast and very slow deposition of free TAF15 aggregates into inclusion bodies, for infinitely large half-lives of TAF15 monomers and aggregates. The numerical solutions show an excellent agreement under these limiting cases.

Modeling under physiologically relevant conditions shows that TAF15 inclusion growth is influenced by both nucleation and autocatalytic conversion mechanisms. Optimal growth conditions occur at intermediate rates of free TAF15 aggregate deposition, striking a balance between the deposition rate of free TAF15 aggregates into the inclusion and the availability of these aggregates to catalyze the production of more aggregates. Sensitivity analysis indicates that inclusion size is highly responsive to variations in the kinetic constants for both nucleation and autocatalytic processes. Additionally, the model predicts a strong dependence of inclusion growth on the monomer production rate, a limiting factor under certain conditions.

In future research, the model should be extended to simulate TAF15 aggregation dynamics across interconnected neuronal networks [34]. This could elucidate how inclusion formation propagates and affects neuronal function in disease progression. The model should be calibrated by comparing its predictions with future experimental data so that it could guide therapeutic strategies toward finding effective treatments for FTLD. Additionally, it is important to extend the model to investigate how TAF15 interacts with other proteins implicated in neurodegeneration, exploring their potential synergistic effects on aggregation. In future research, it is crucial to conduct experiments that could either validate or refute the predictions of the developed model.

## Abbreviations

Aβ: amyloid beta
AD: Alzheimer’s disease
CV: control volume
EWS: Ewing’s sarcoma
F-W: Finke-Watzky
FTLD: Frontotemporal lobar degeneration
FUS: fused in sarcoma
TAF15: TATA-box binding protein associated factor 15
TDP-43: transactive response DNA binding protein of 43 kDa
TET: ten-eleven translocation

## Data accessibility

This article has no additional data.

## Authors’ contributions

AVK is the sole author of this paper.

## Supplemental Materials

### S1. Analytical solution of Eqs. (25) and (26) for the case where free TAF15 aggregates deposit slowly into TAF15 inclusions, 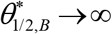 , and the half-lives of TAF15 monomers and aggregates are infinitely large, 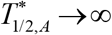 and 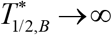

Adding Eqs. (25) and (26) gives

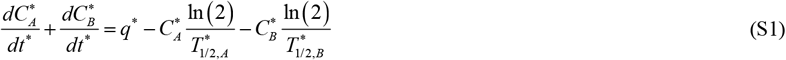

Assuming 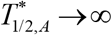 and 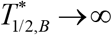 , and using the initial conditions provided in Eq. (11a,b), Eq. (S1) was integrated with respect to time, yielding

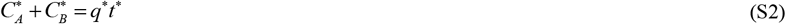

By substituting Eq. (S2) into Eq. (26) to eliminate 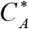 , the following equation is obtained:

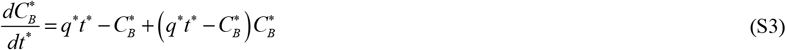

In ref. [11], a similar equation to Eq. (S3) was examined. To obtain the exact solution of Eq. (S3) with the initial condition provided by Eq. (11b), the DSolve function followed by the FullSimplify function in Mathematica 13.3 (Wolfram Research, Champaign, IL) was employed. The resulting solution is

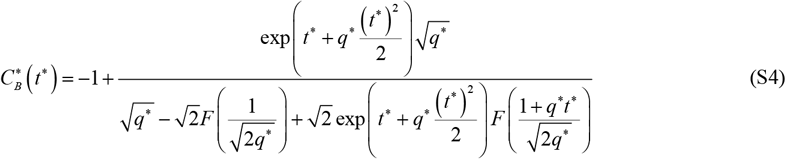

In this equation, *F* (*x* ) denotes Dawson’s integral:

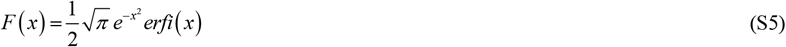

where *erfi* (*x*) is the imaginary error function.

Assuming *t*^∗^ → 0 , Eq. (S4) leads to

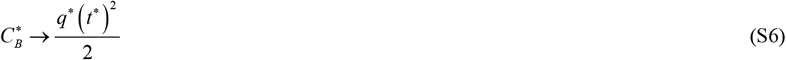

Then 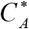 is found using Eq. (S2) as:

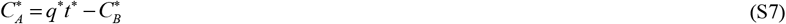

The exact solution given by Eqs. (S4) and (S7) is quite cumbersome. A more elegant approximate solution, also applicable under conditions where 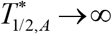 and 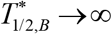 , is as follows:

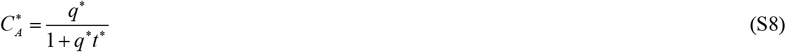

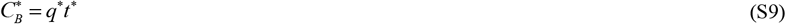

As *t*^∗^ →∞ , Eq. (S8) indicates that 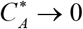 . Note that Eqs. (S8) and (S9) are suitable for larger time scales but become inapplicable as *t*^∗^ → 0 , as Eq. (S8) contradicts the initial condition specified in Eq. (11a).

In line with Eq. (27), in the scenario where 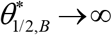 ,

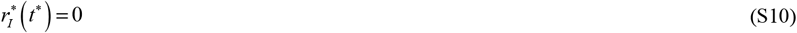

### S2. Numerical solution

Eqs. (3), (5), and (6) along with their initial conditions in (7) were solved numerically using MATLAB’s ODE45 solver (MATLAB R2020b, MathWorks, Natick, MA, USA), widely recognized as reliable. To ensure precise solutions, the error tolerance parameters RelTol and AbsTol were set to 1e-10.

### S3. Supplementary figures

**Fig. S1.**
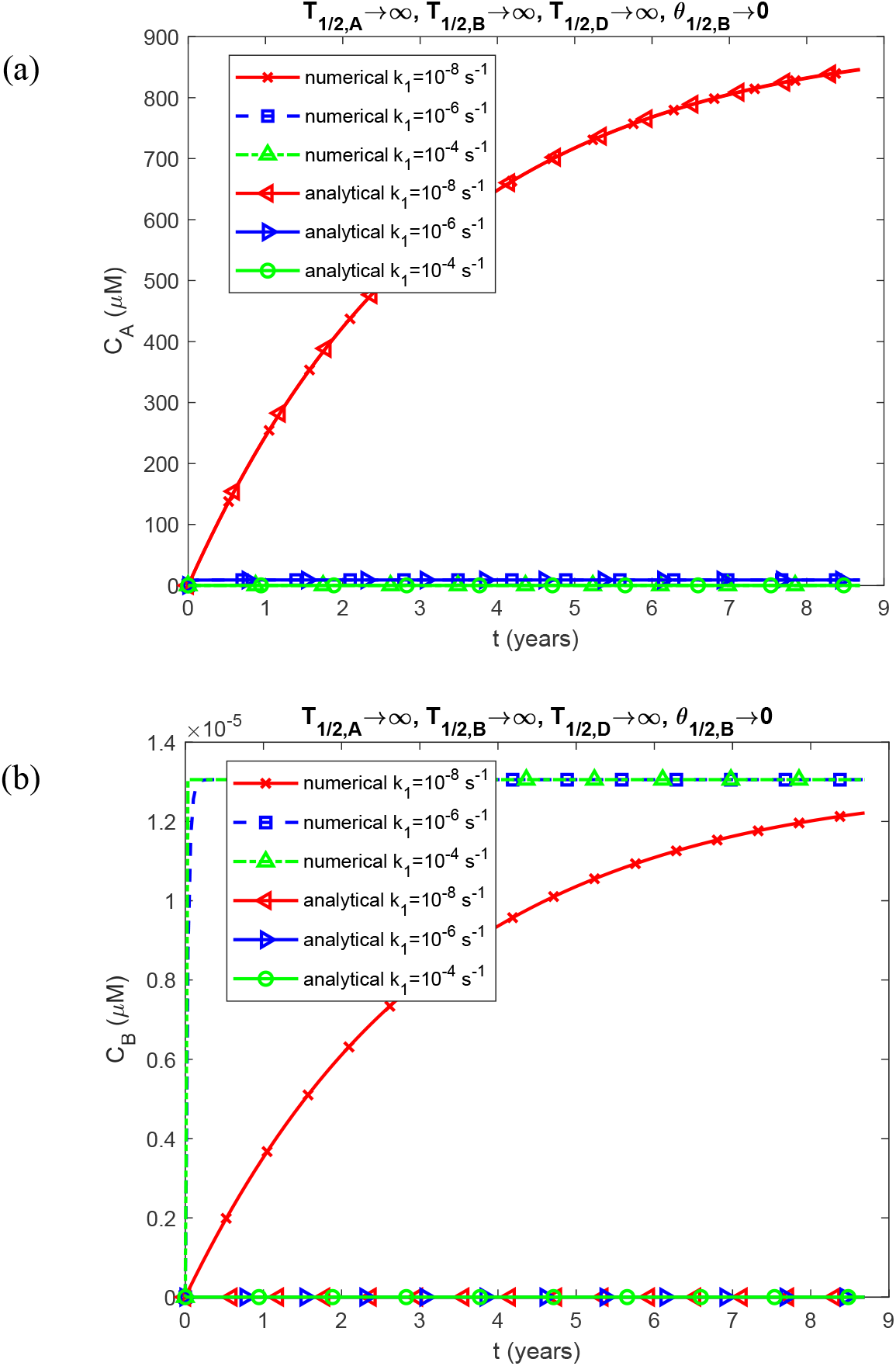
(a) The molar concentration of TAF15 monomers, *C*_*A*_ , plotted against time; and (b) the molar concentration of free TAF15 aggregates (not deposited into inclusions), *C*_*B*_ , plotted against time, for various values of *k*_1_ . The scenario with *T*_1/2,*A*_ →∞ , *T*_1/2,*B*_ →∞ , *T*_1/2,*D*_ →∞, and *θ*_1/2,*B*_ →0 is presented. The analytical solution derived from Eqs. (12), (19), (24), and (36) is utilized to validate the numerical solution shown in Fig. S1. *k*_2_ = 10^−6^ μM^-1^ s^-1^ and *q* = 3.79×10^−23^ mol s^-1^. Other parameters follow those specified in Table 2.

**Fig. S2.**
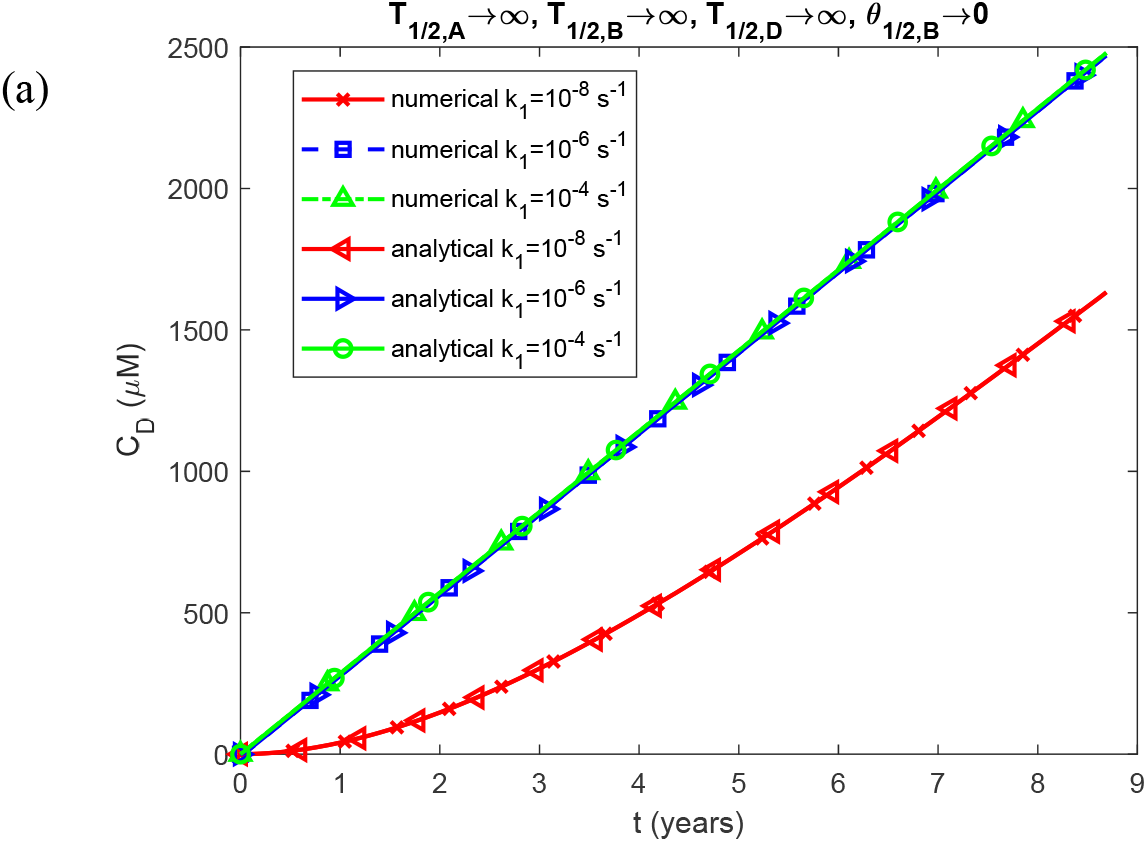
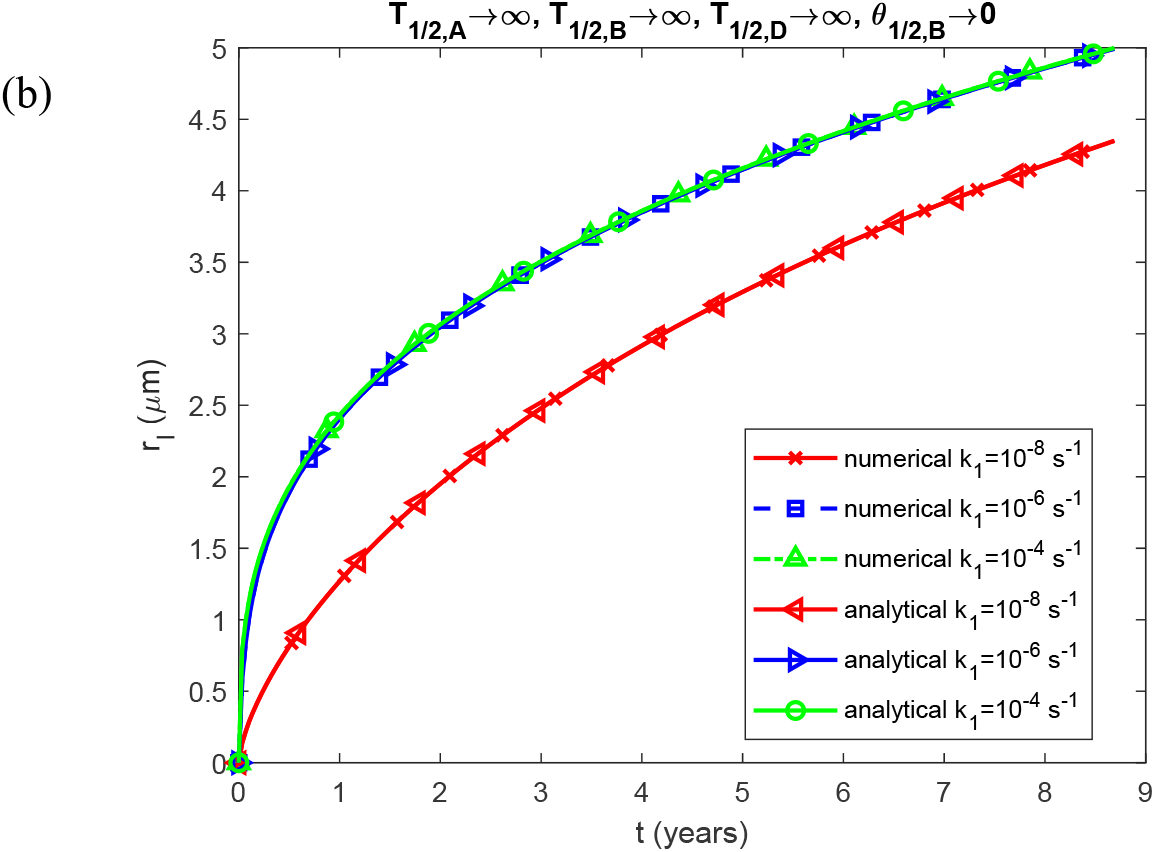
(a) The molar concentration of TAF15 aggregates deposited into TAF15 inclusions, *C*_*D*_ , plotted against time; and (b) the radius of a TAF15 inclusion, *r*_*I*_ , plotted against time, for various values of *k*_1_ . The scenario with *T*_1/2,*A*_ →∞ , *T*_1/2,*B*_ →∞ , *T*_1/2,*D*_ →∞, and *θ*_1/2,*B*_ →0 is presented. The analytical solution derived from Eqs. (12), (19), (24), and (36) is used to validate the numerical solution shown in Fig. S2. *k*_2_= 10^−6^ μM^-1^ s^-1^ and *q* = 3.79×10^−23^ mol s^-1^. Other parameters follow those specified in Table 2.

**Fig. S3.**
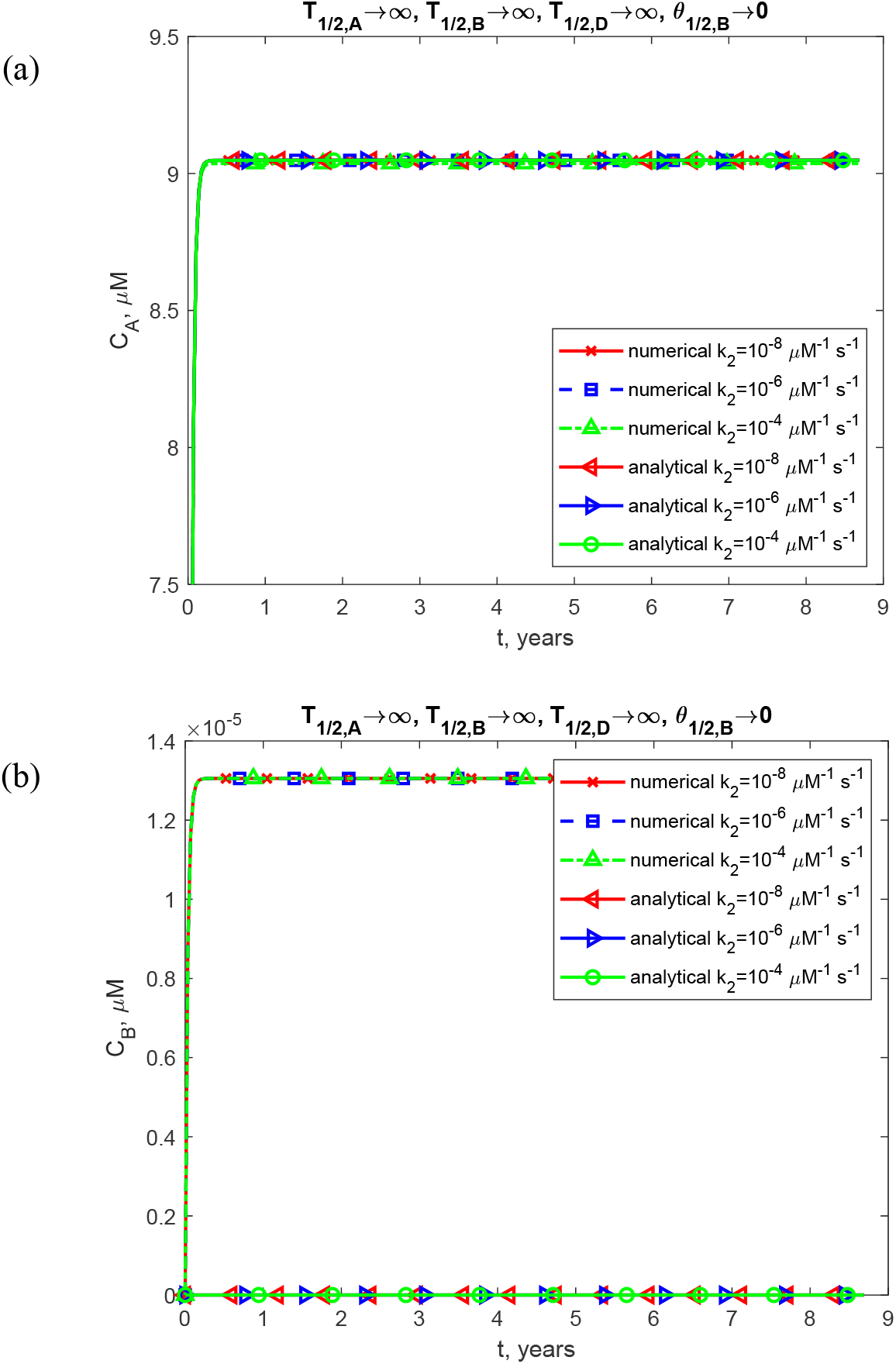
(a) The molar concentration of TAF15 monomers, *C*_*A*_ , plotted against time; and (b) the molar concentration of free TAF15 aggregates (not deposited into inclusions), *C*_*B*_ , plotted against time, for various values of *k*_2_ . The scenario with *T*_1/2,*A*_ →∞ , *T*_1/2,*B*_ →∞ , *T*_1/2,*D*_ →∞, and *θ*_1/2,*B*_ →0 is presented. The analytical solution derived from Eqs. (12), (19), (24), and (36) is used to validate the numerical solution shown in Fig. S3. *K*_1_ = 10 ^−6^ and *q* = 3.79×10^−23^ mol s^-1^. Other parameters follow those specified in Table 2.

**Fig. S4.**
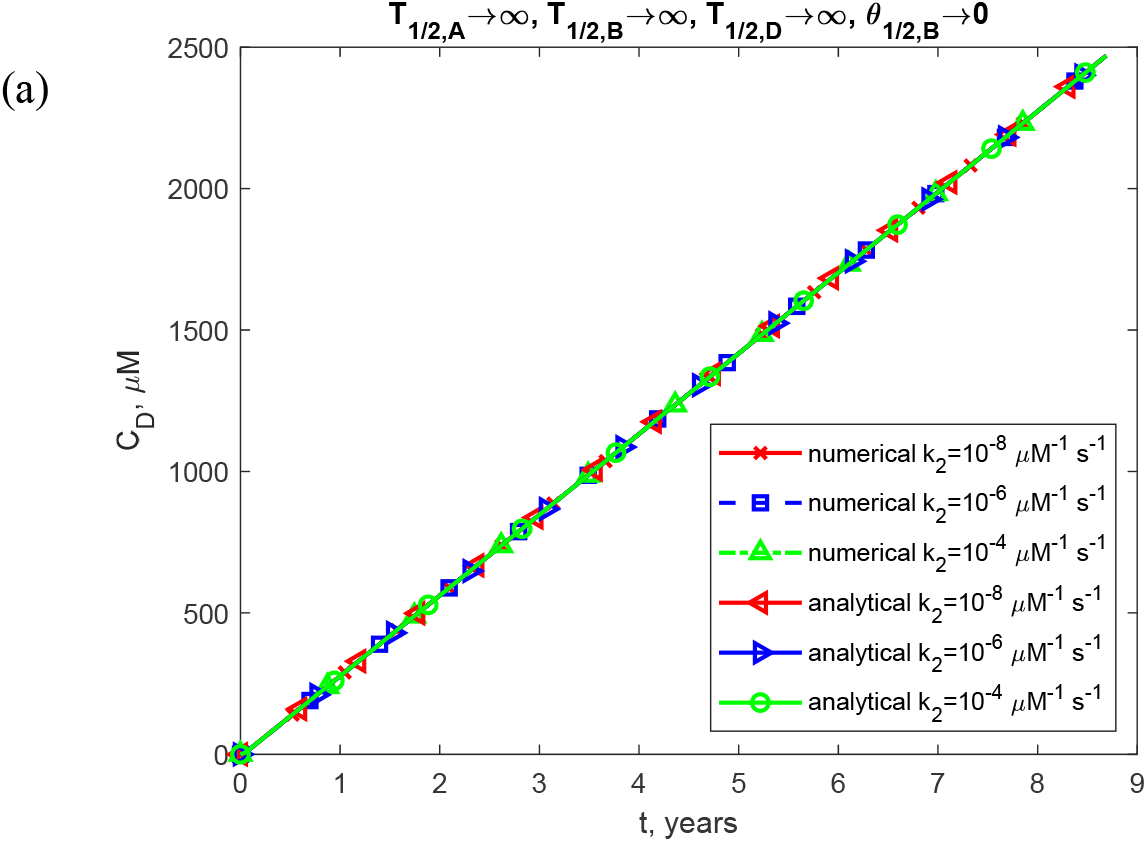
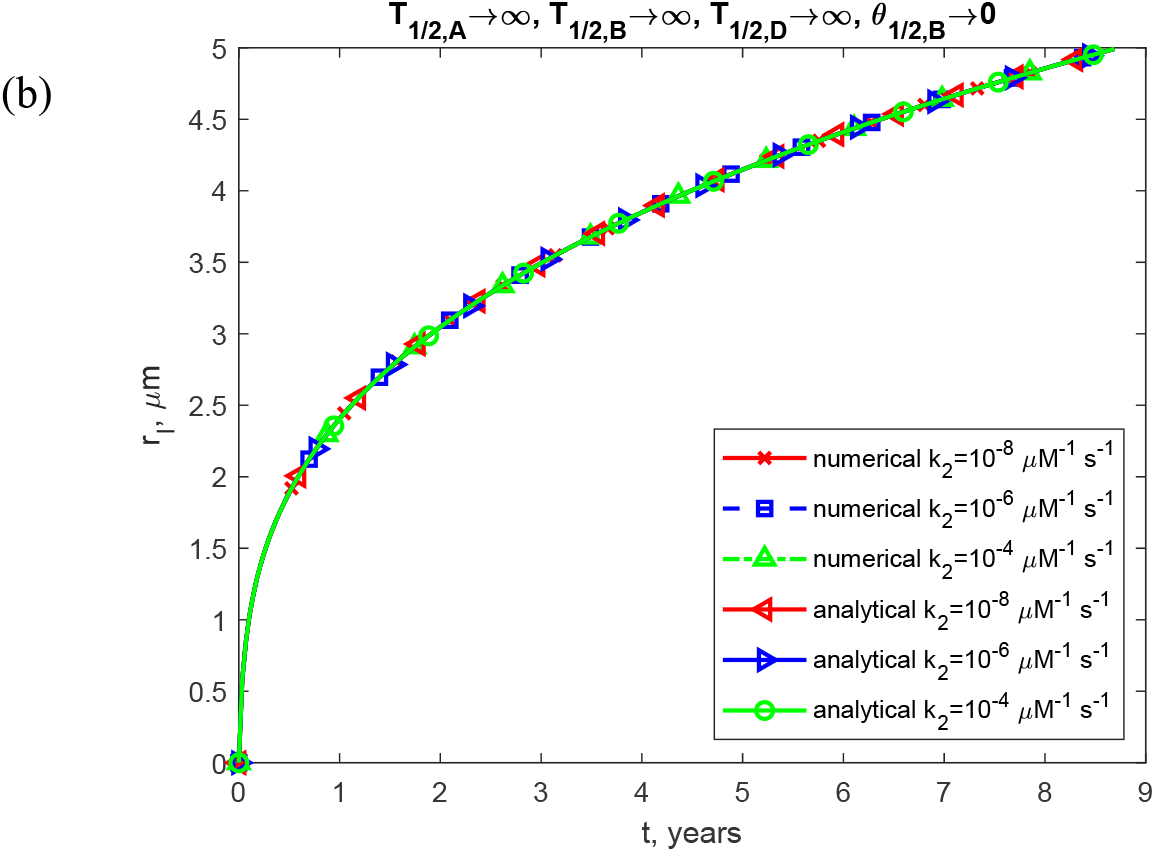
(a) The molar concentration of TAF15 aggregates deposited into TAF15 inclusions, *C*_*D*_ , plotted against time; and (b) the radius of a TAF15 inclusion, *r*_*I*_ , plotted against time, for various values of *k*_2_ . The scenario with *T*_1/2,*A*_ →∞ , *T*_1/2,*B*_ →∞ , *T*_1/2,*D*_ →∞, and *θ*_1/2,*B*_ →0 is presented. The analytical solution derived from Eqs. (12), (19), (24), and (36) is used to validate the numerical solution shown in Fig. S4.*k*_1_ = 10 ^−6^ s^-1^ and *q* = 3.79×10^−23^ mol s^-1^. Other parameters follow those specified in Table 2.

**Fig. S5.**
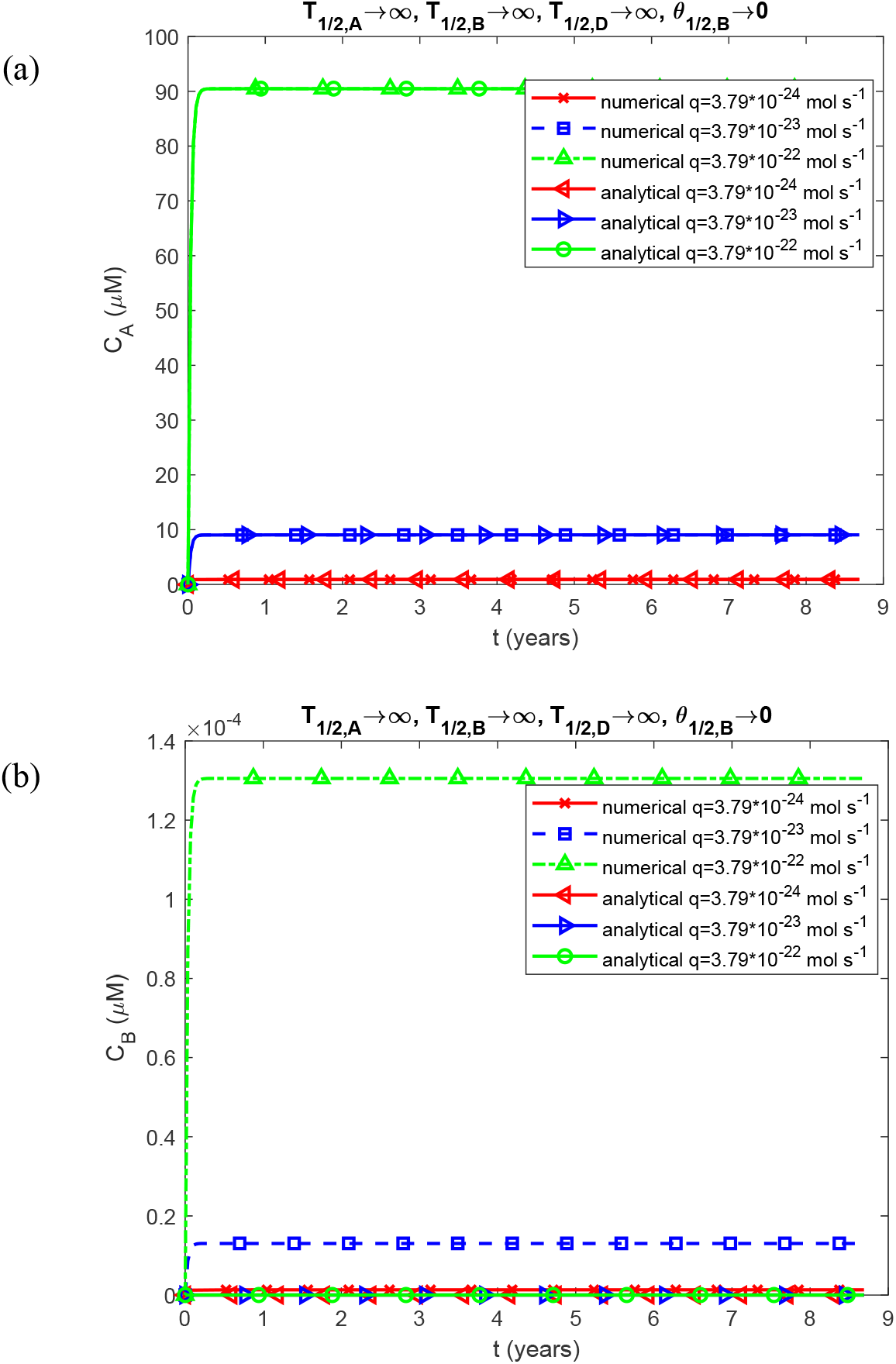
(a) The molar concentration of TAF15 monomers, *C*_*A*_ , plotted against time; and (b) the molar concentration of free TAF15 aggregates (not deposited into inclusions), *C*_*B*_ , plotted against time, for various values of *q* . The scenario with *T*_1/2,*A*_ →∞ , *T*_1/2,*B*_ →∞ , *T*_1/2,*D*_ →∞, and *θ*_1/2,*B*_ →0 is presented. The analytical solution derived from Eqs. (12), (19), (24), and (36) is used to validate the numerical solution shown in Fig. S5. *k*_1_ = 10 ^−6^ s^-1^ and *k*_2_= 10^−6^μM^-1^ s^-1^. Other parameters follow those specified in Table 2.

**Fig. S6.**
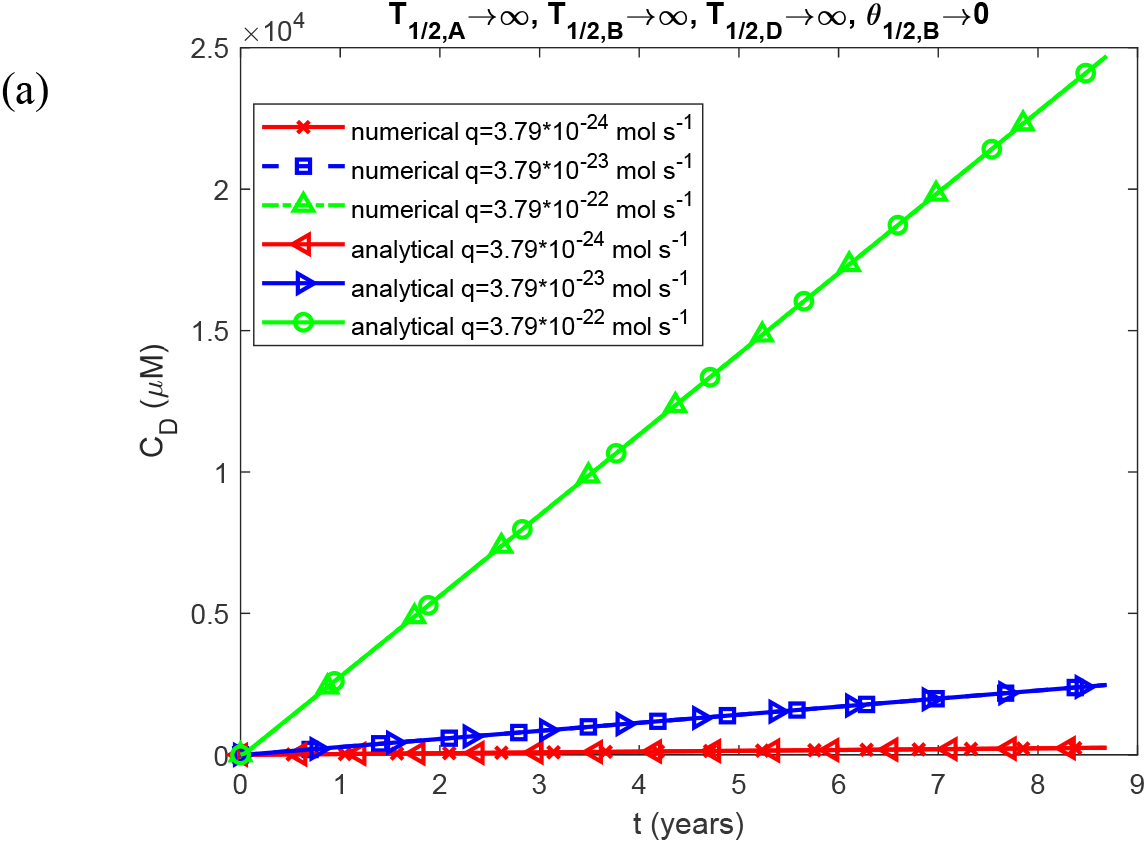
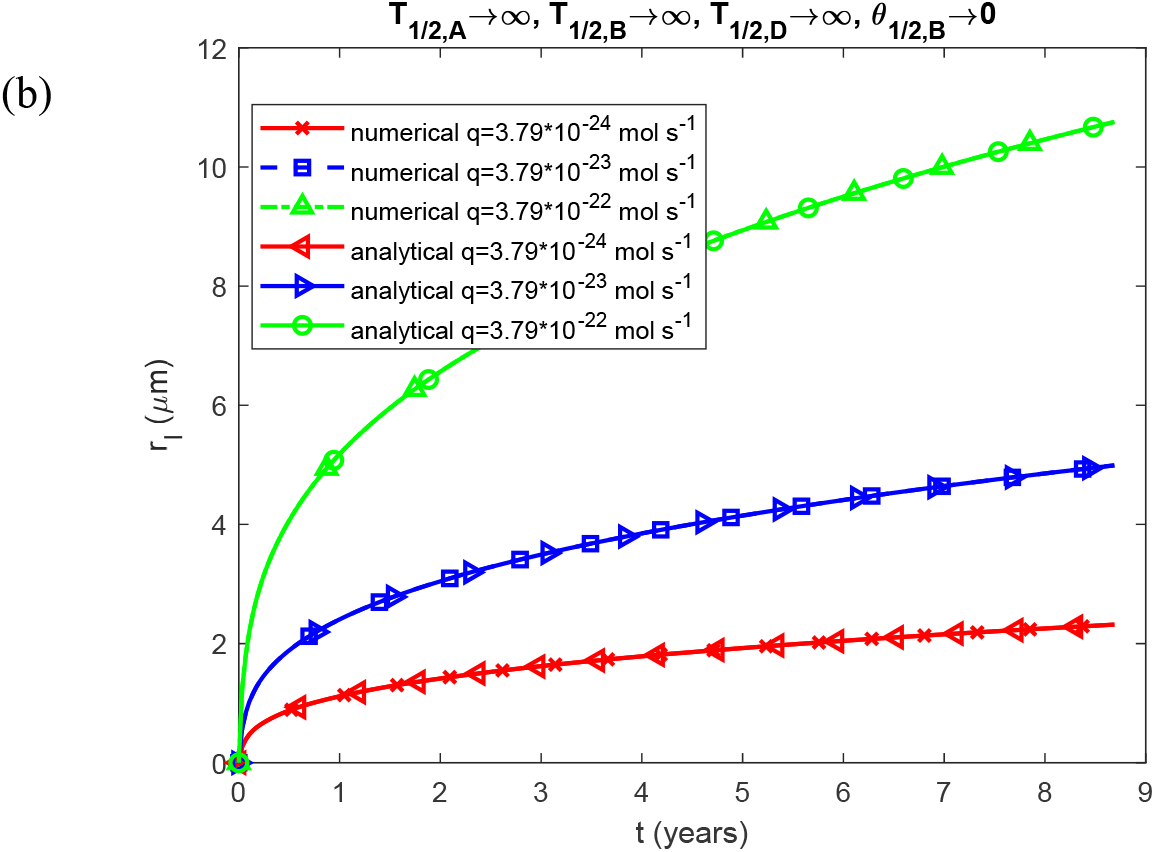
(a) The molar concentration of TAF15 aggregates deposited into TAF15 inclusions, *C*_*D*_ , plotted against time; and (b) the radius of a TAF15 inclusion, *r*_*I*_ , plotted against time, for various values of *q* . The scenario with *T*_1/2,*A*_ →∞ , *T*_1/2,*B*_ →∞ , *T*_1/2,*D*_ →∞, and *θ*_1/2,*B*_ →0 is presented. The analytical solution derived from Eqs. (12), (19), (24), and (36) is used to validate the numerical solution shown in Fig. S6. *k*__1__ = 10 ^−6^ s^-1^ and *k*_2_ = 10^−6^ μM^-1^ s^-1^. Other parameters follow those specified in Table 2.

**Fig. S7.**
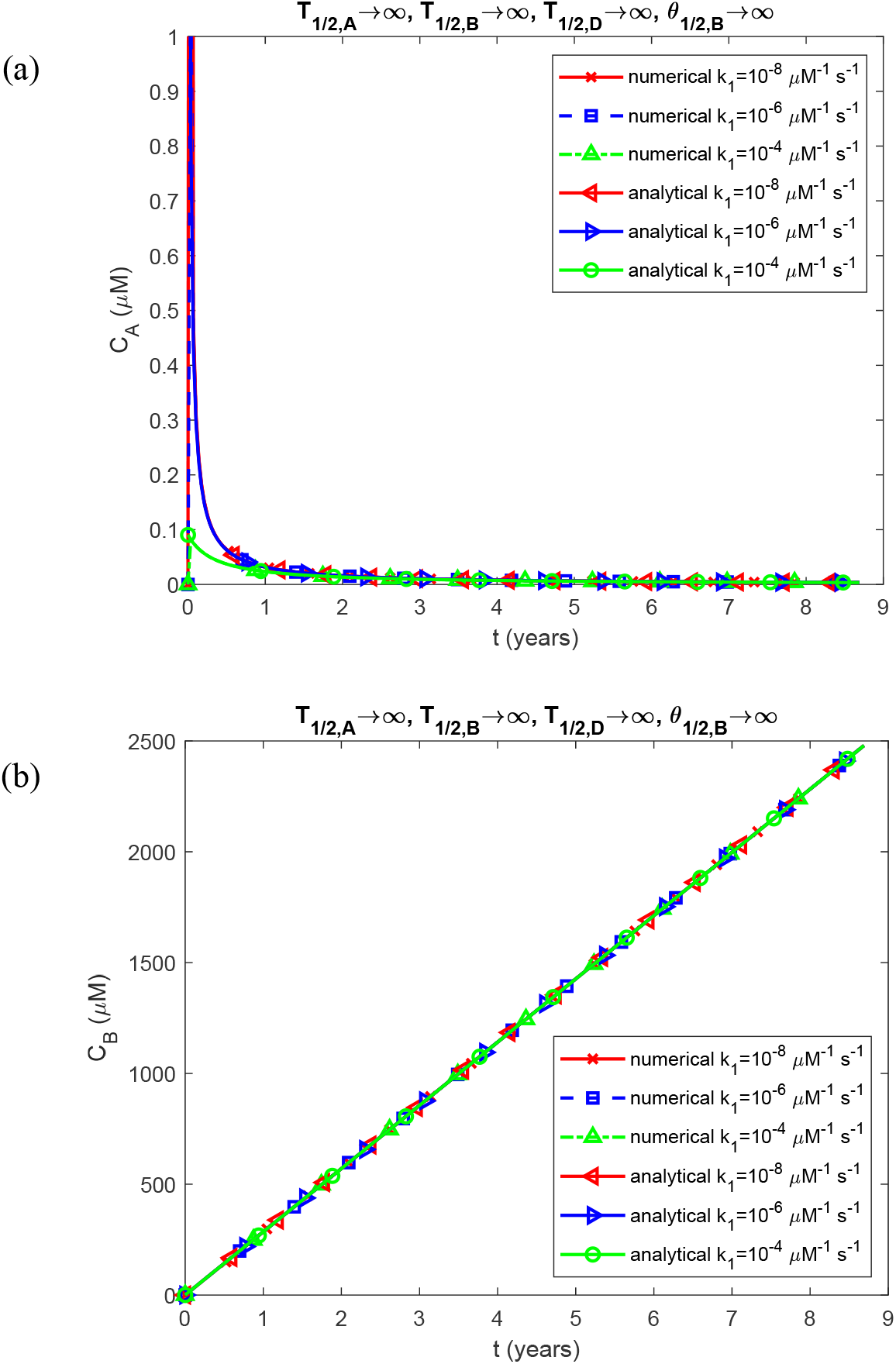
(a) The molar concentration of TAF15 monomers, *C*_*A*_ , plotted against time; and (b) the molar concentration of free TAF15 aggregates (not deposited into inclusions), *C*_*B*_ , plotted against time, for various values of *k*_1_ . The scenario with *T*_1/2,*A*_ →∞ , *T*_1/2,*B*_ →∞ , *T*_1/2,*D*_ →∞, and *θ*_1/2,*B*_ →∞ is presented. The approximate analytical solution derived from Eqs. (S8), (S9), (27), and (S10) is used to validate the numerical solution shown in Fig. S7. *K*_2_ = 10^−6^ μM^-1^ s^-1^ and *q* = 3.79×10^−23^ mol s^-1^. Other parameters follow those specified in Table 2.

**Fig. S8.**
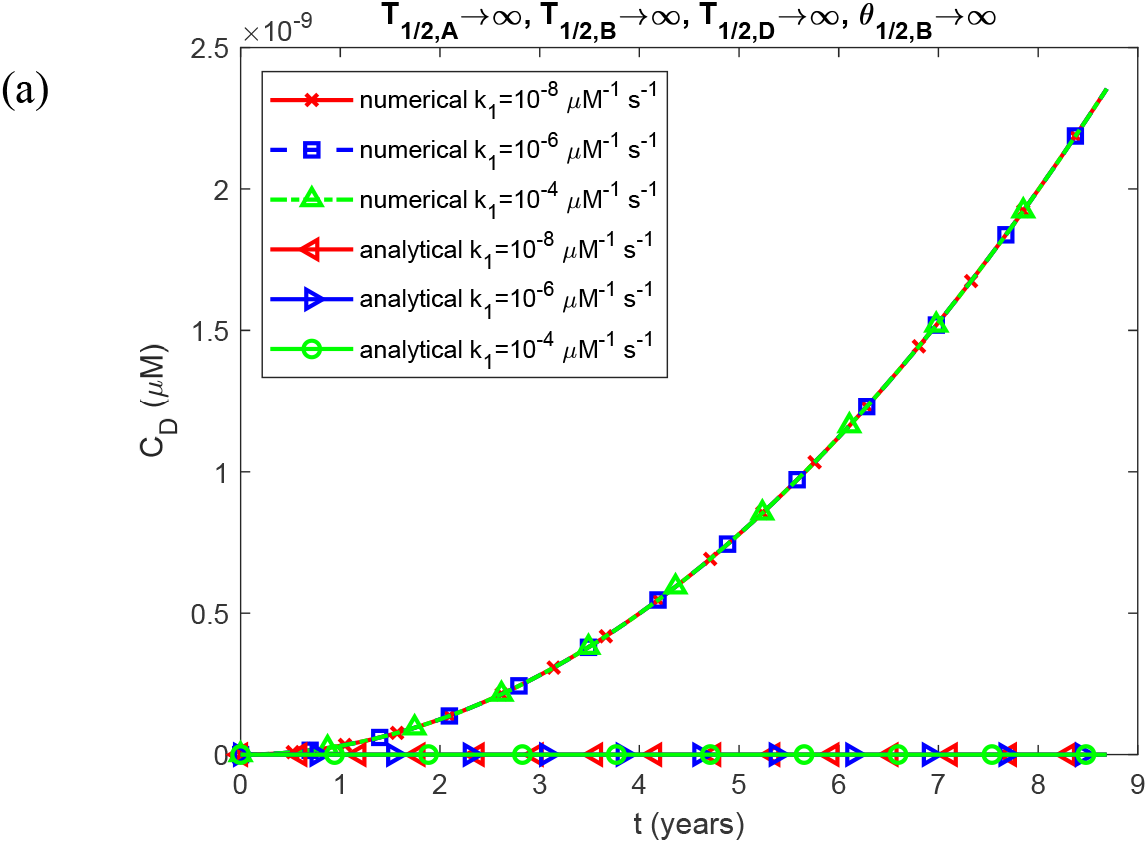
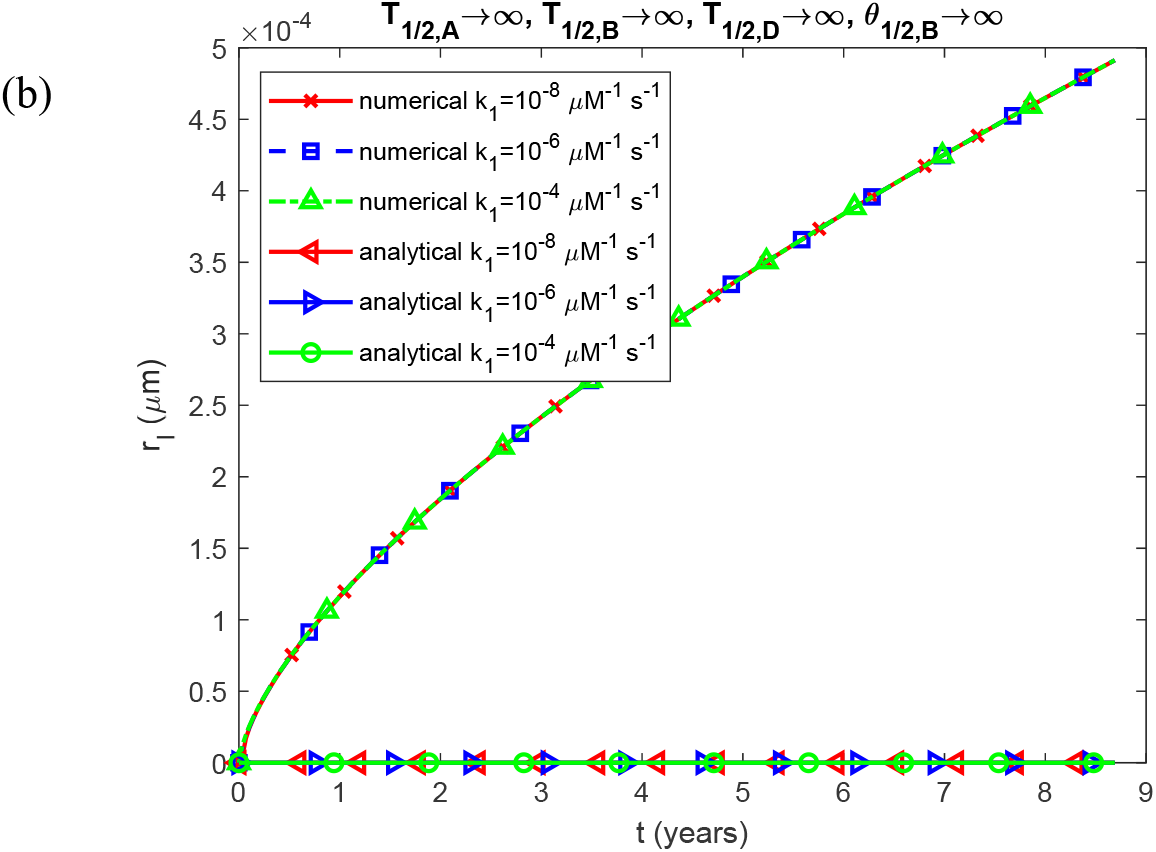
(a) The molar concentration of TAF15 aggregates deposited into TAF15 inclusions, *C*_*D*_ , plotted against time; and (b) the radius of a TAF15 inclusion, *r*_*I*_ , plotted against time, for various values of *k*_1_ . The scenario with *T*_1/2,*A*_ →∞ , *T*_1/2,*B*_ →∞ , *T*_1/2,*D*_ →∞, and *θ*_1/2,*B*_ →∞ is presented. The approximate analytical solution derived from Eqs. (S8), (S9), (27), and (S10) is used to validate the numerical solution shown in Fig. S8. *k*_2_= 10^−6^ μM^-1^ s^-1^ and *q* = 3.79×10^−23^ mol s^-1^. Other parameters follow those specified in Table 2.

**Fig. S9.**
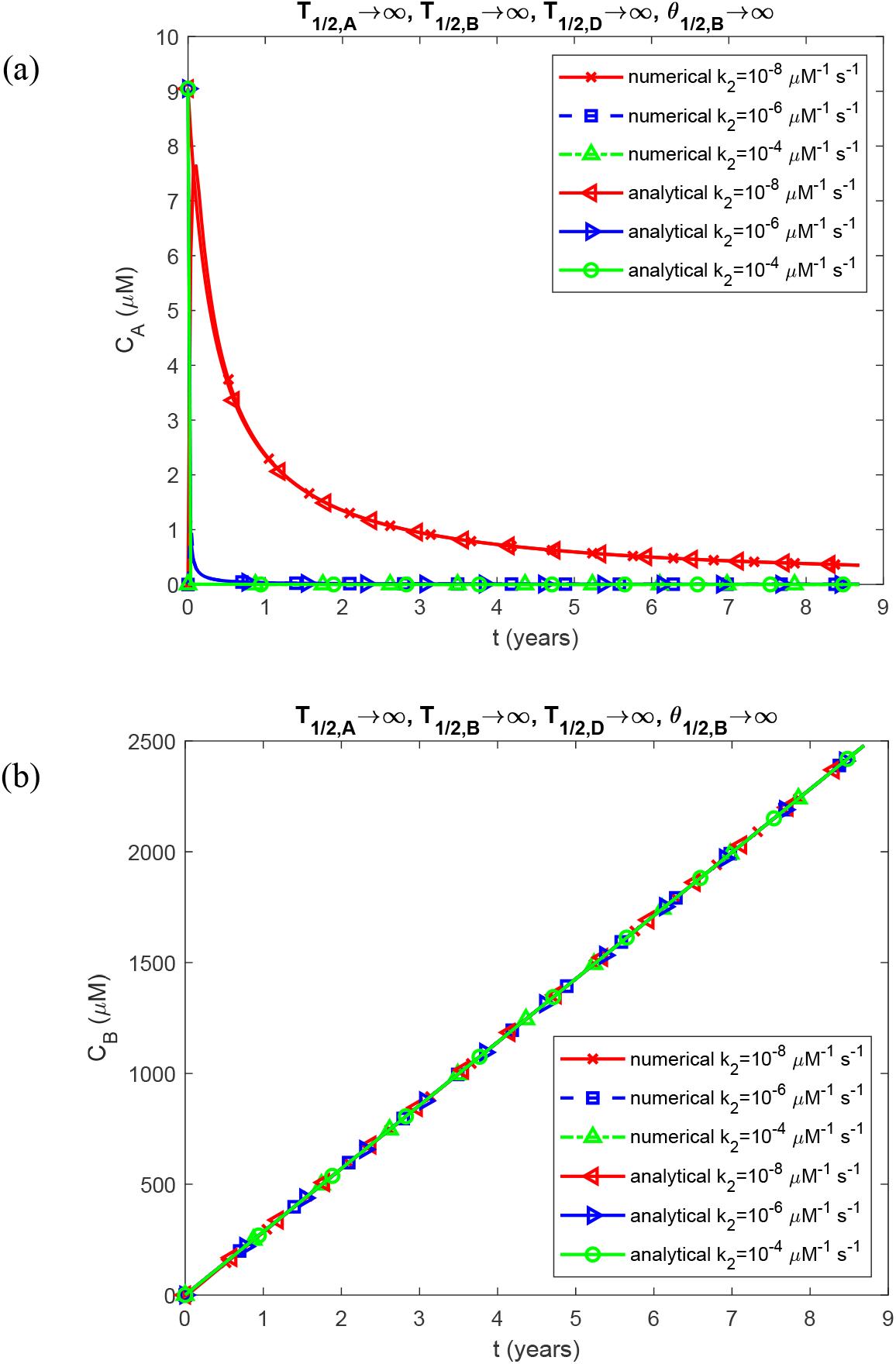
(a) The molar concentration of TAF15 monomers, *C*_*A*_ , plotted against time; and (b) the molar concentration of free TAF15 aggregates (not deposited into inclusions), *C*_*B*_ , plotted against time, for various values of *k*_2_ . The scenario with *T*_1/2,*A*_ →∞ , *T*_1/2,*B*_ →∞ , *T*_1/2,*D*_ →∞, and *θ*_1/2,*B*_ →∞ is presented. The approximate analytical solution from Eqs. (S8), (S9), (27), and (S10) is used to validate the numerical solution shown in Fig. S9. *K*_1_ = 10 ^−6^ s^-1^ and *q* = 3.79×10^−23^ mol s^-1^. Other parameters are as specified in Table 2.

**Fig. S10.**
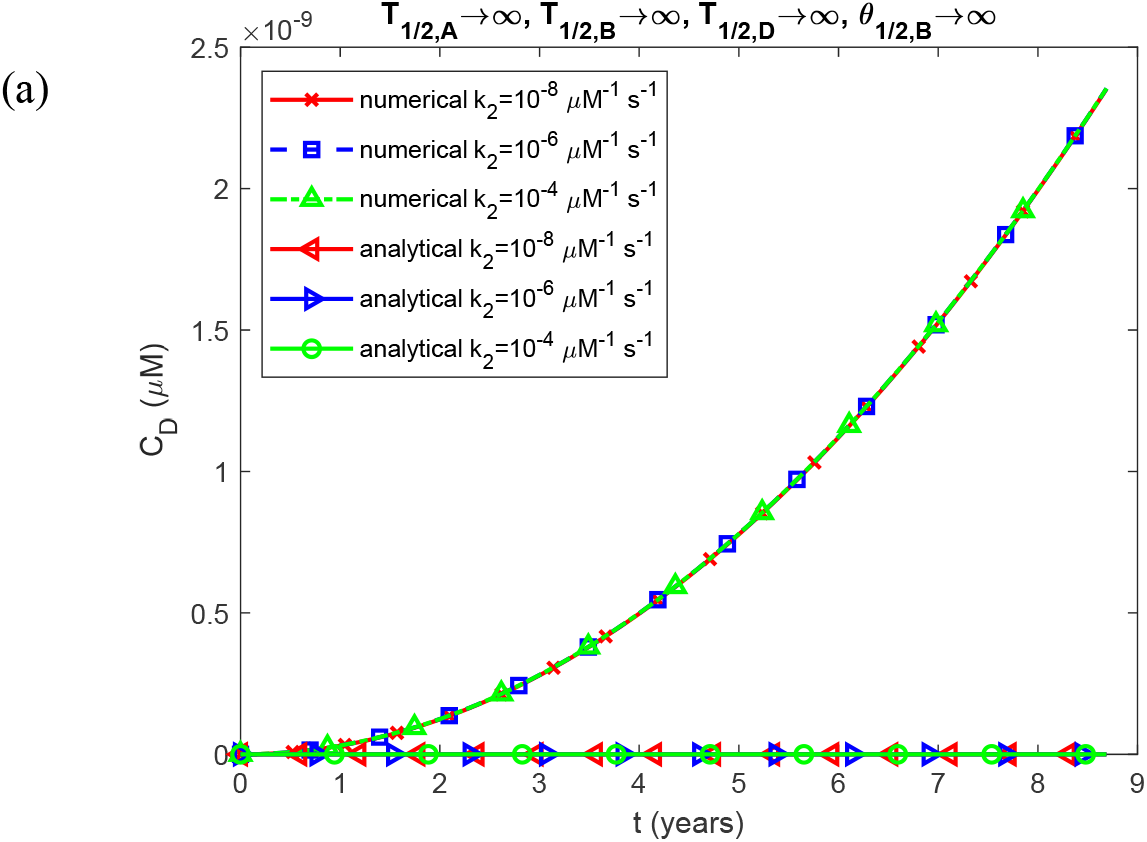
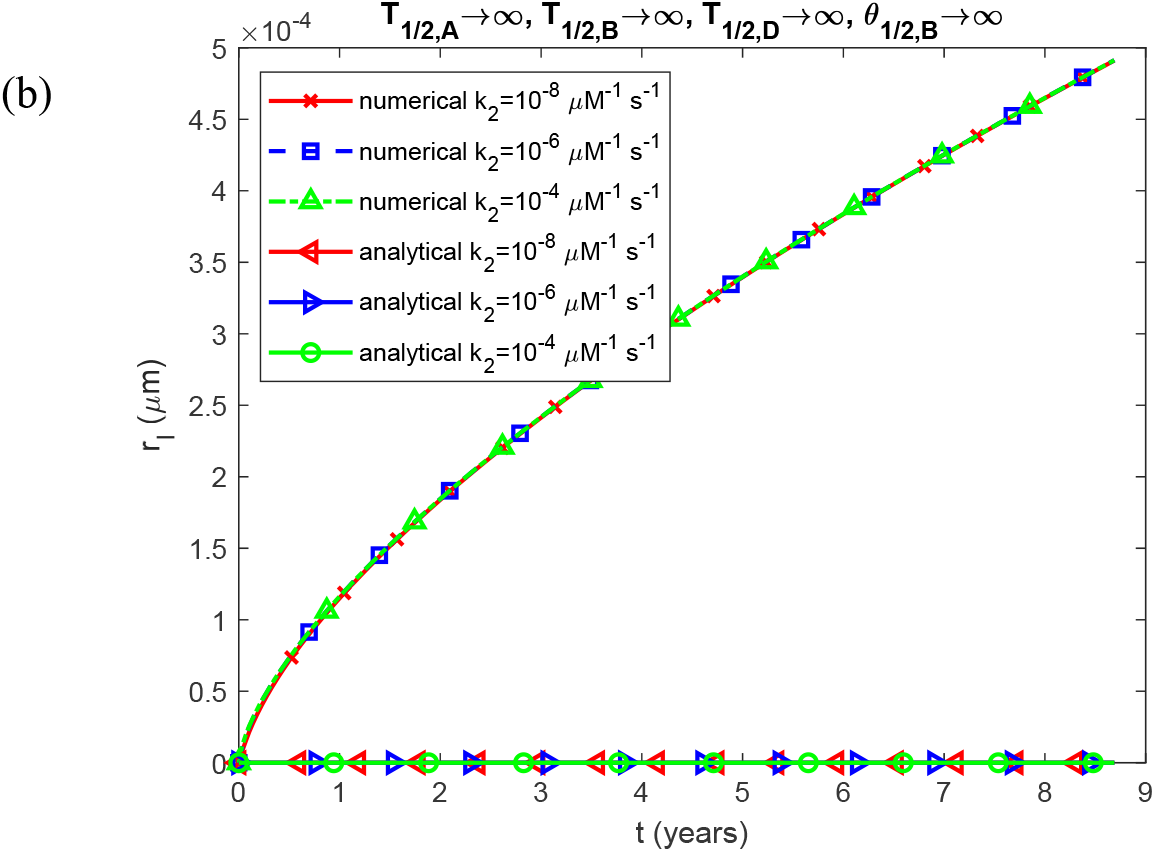
(a) The molar concentration of TAF15 aggregates deposited into TAF15 inclusions, *C*_*D*_ , plotted against time; and (b) the radius of a TAF15 inclusion, *r*_*I*_ , plotted against time, for various values of *k*_2_ .The scenario with *T*_1/2,*A*_ →∞ , *T*_1/2,*B*_ →∞ , *T*_1/2,*D*_ →∞, and *θ*_1/2,*B*_ →∞ is presented. The approximate analytical solution from Eqs. (S8), (S9), (27), and (S10) is used to validate the numerical solution shown in Fig. S10. *k*_1_ = 10 ^−6^ s^-1^ and *q* = 3.79×10^−23^ mol s^-1^. Other parameters are as specified in Table 2.

**Fig. S11.**
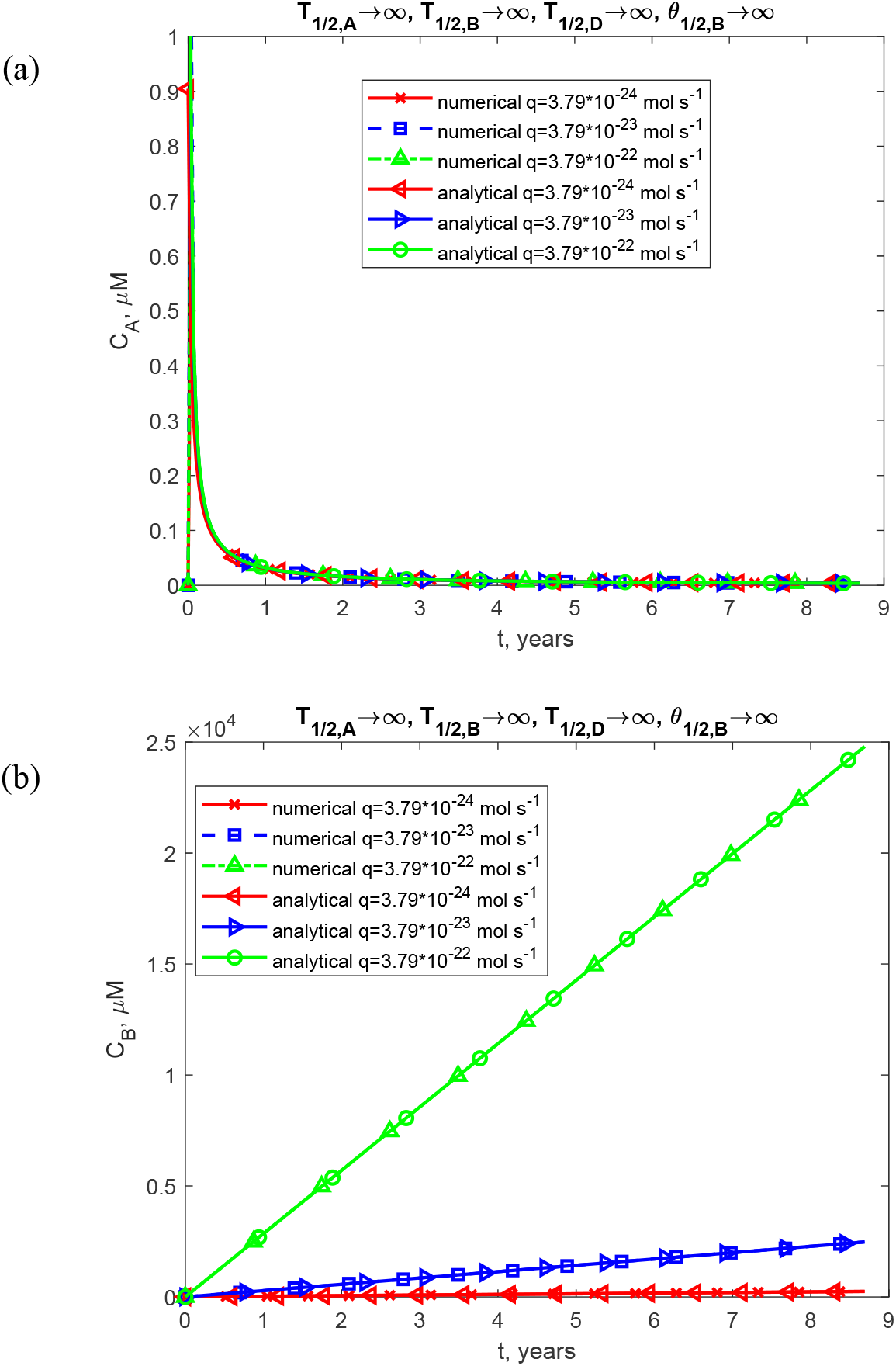
(a) The molar concentration of TAF15 monomers, *C*_*A*_ , plotted against time; and (b) the molar concentration of free TAF15 aggregates (not deposited into inclusions), *C*_*B*_ , plotted against time, for various values of *q* . The scenario with *T*_1/2,*A*_ →∞ , *T*_1/2,*B*_ →∞ , *T*_1/2,*D*_ →∞, and *θ*_1/2,*B*_ →∞ is presented. The approximate analytical solution from Eqs. (S8), (S9), (27), and (S10) is used to validate the numerical solution shown in Fig. S11. *K*_1_ = 10 ^−6^ s^-1^ and *k*_2_ = 10^−6^μM^-1^ s^-1^. Other parameters are as specified in Table 2.

**Fig. S12.**
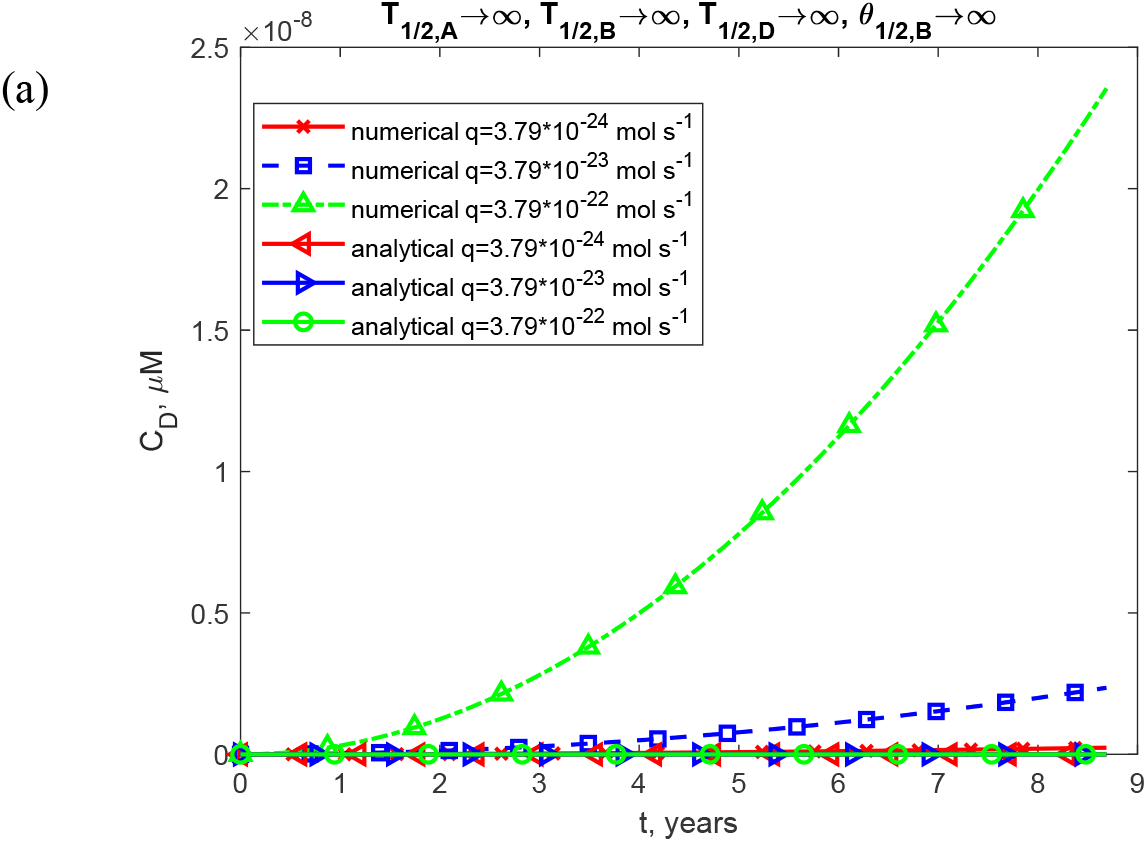
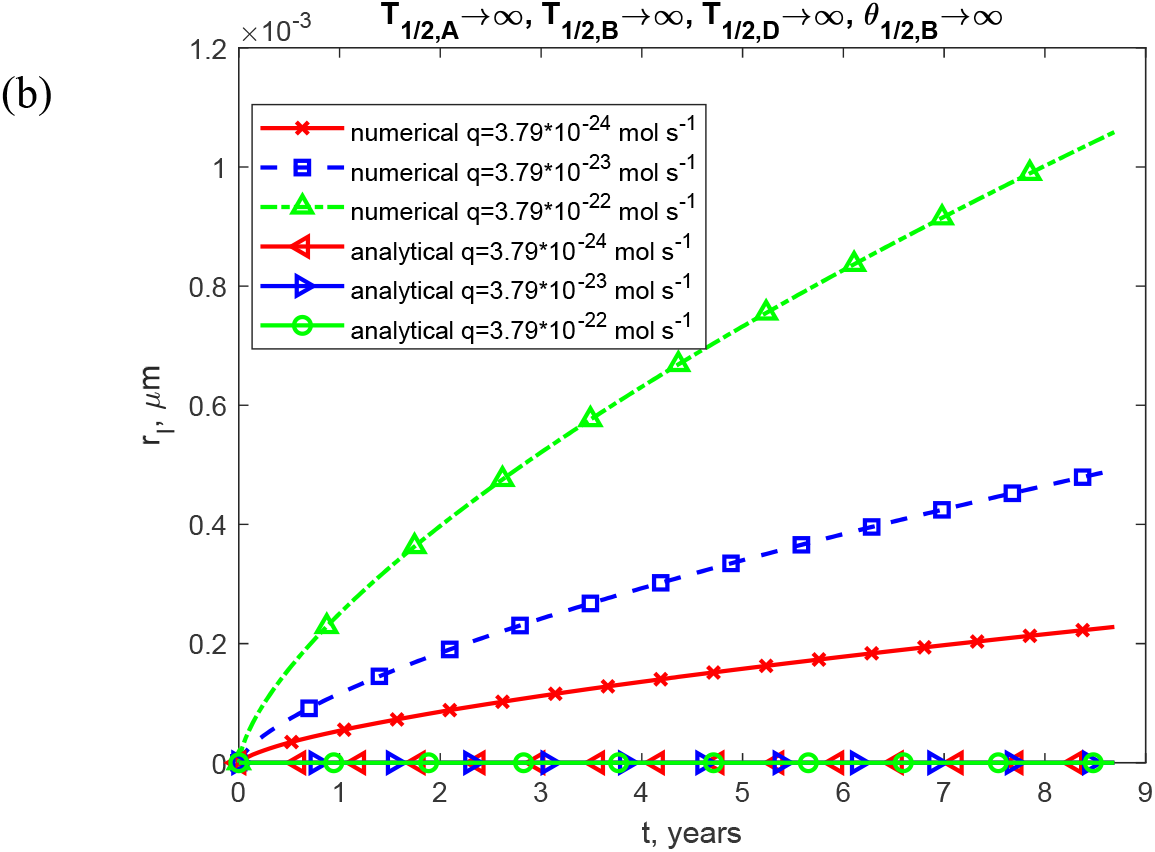
(a) The molar concentration of TAF15 aggregates deposited into TAF15 inclusions, *C*_*D*_ , plotted against time; and (b) the radius of a TAF15 inclusion, *r*_*I*_ , plotted against time, for various values of *q* . The scenario with *T*_1/2,*A*_ →∞ , *T*_1/2,*B*_ →∞ , *T*_1/2,*D*_ →∞, and *θ*_1/2,*B*_ →∞ is presented. The approximate analytical solution from Eqs. (S8), (S9), (27), and (S10) is used to verify the accuracy of the numerical solution. *K*_1_ = 10 ^−6^ s^-1^ and *k*_2_ = 10^−6^ μM^-1^ s^-1^. Other parameters are as specified in Table 2.

**Fig. S13.**
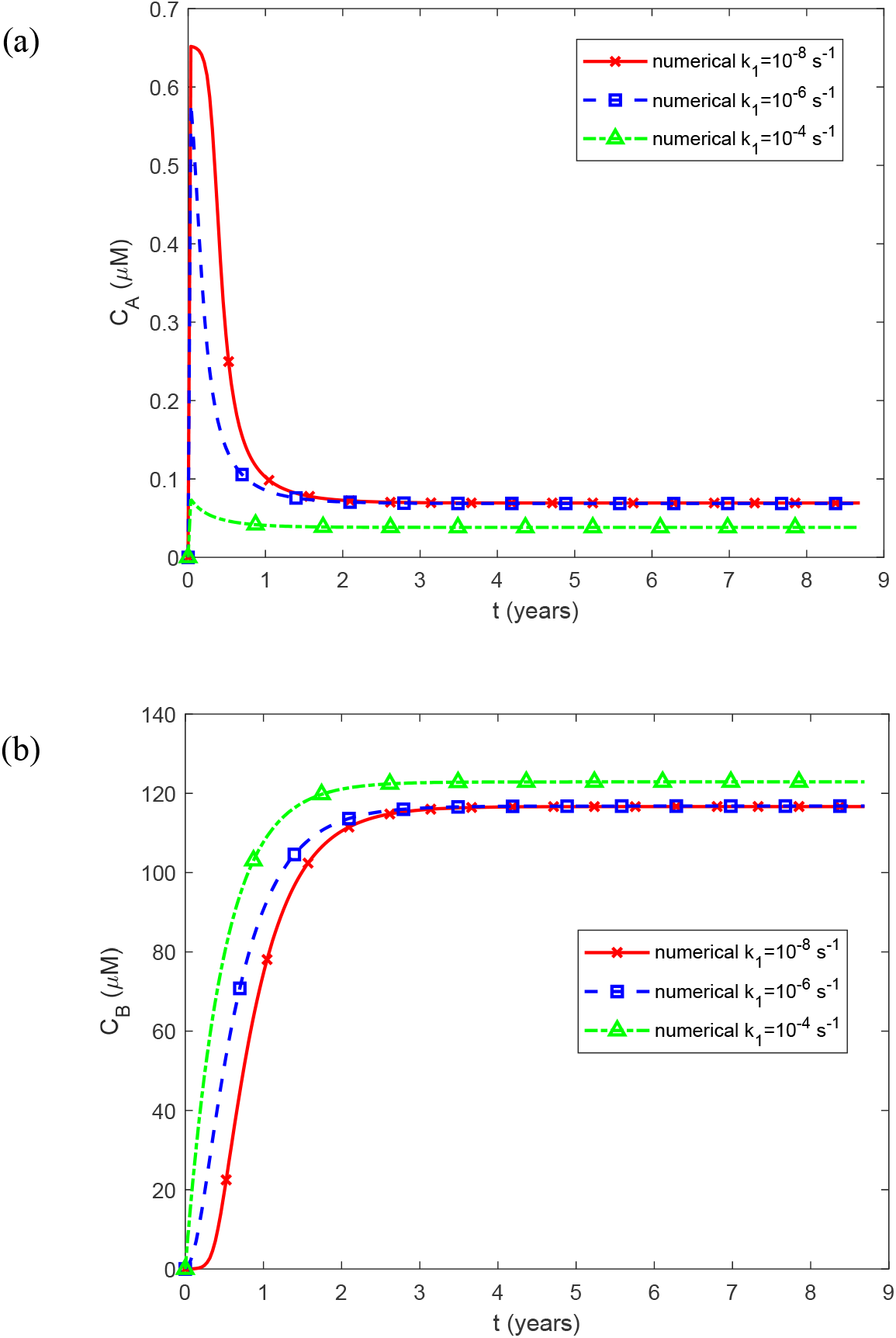
(a) The molar concentration of TAF15 monomers, *C*_*A*_ , plotted against time; and (b) the molar concentration of free TAF15 aggregates (not deposited into inclusions), *C*_*B*_ , plotted against time, for various values of *k*_1_ . Other than for *k*_1_ , physiologically relevant parameter values are utilized: *k*_2_ = 10^−6^ μM^-1^ s^-1^, *q* = 3.79×10^−23^ mol s^-1^, *T*_1/2,*A*_ = 5×10^4^ s, and *θ*_1/2,*B*_ =10^7^ s. Other parameters are as specified in Table 2.

**Fig. S14.**
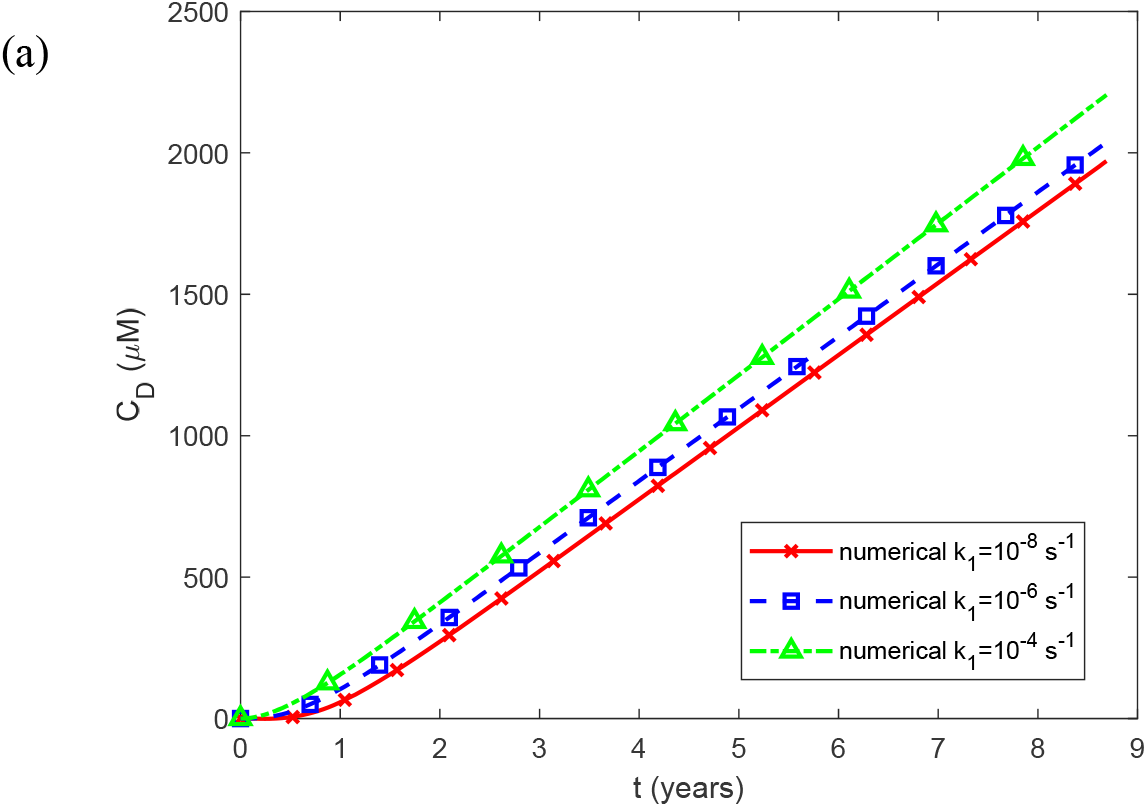
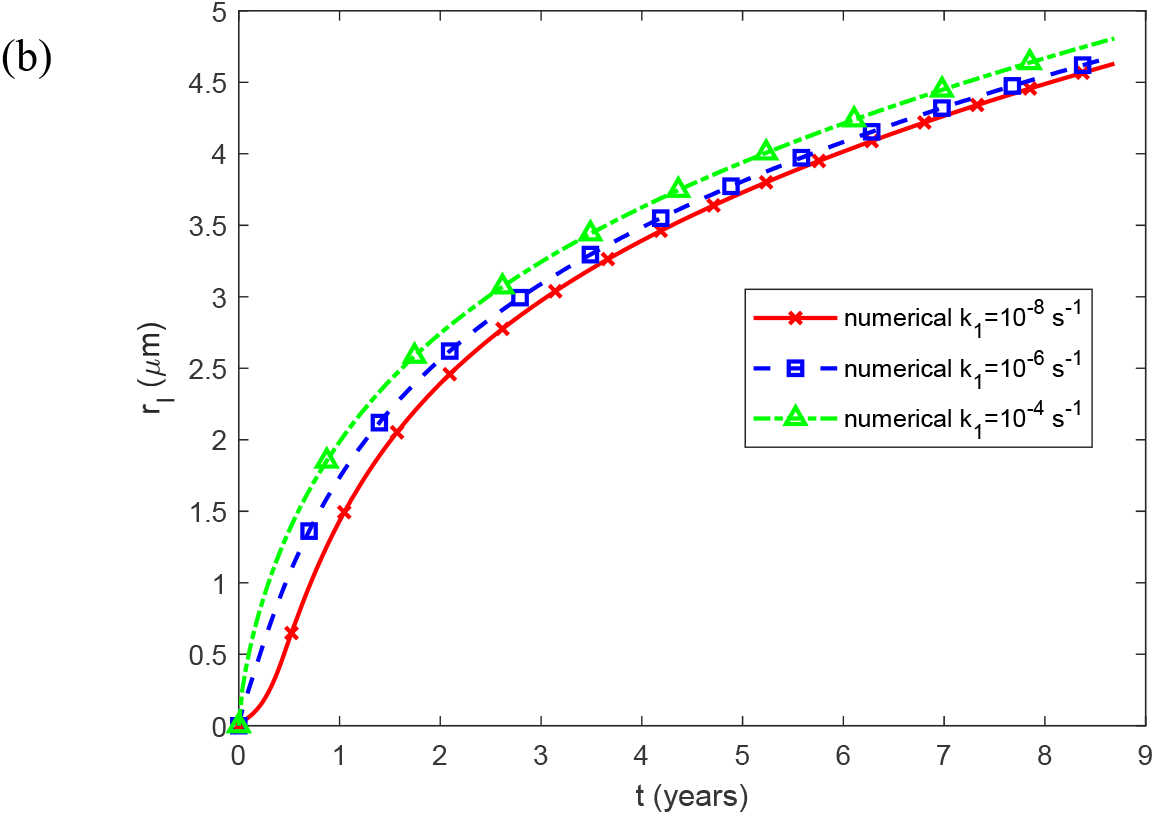
(a) The molar concentration of TAF15 aggregates deposited into TAF15 inclusions, *C*_*D*_ , plotted against time; and (b) the radius of a TAF15 inclusion, *r*_*I*_ , plotted against time, for various values of *k*_1_ . Other than for *k*_1_ , physiologically relevant parameter values are utilized: *k*_2_= 10^−6^ μM^-1^ s^-1^, *q* = 3.79×10^−23^ mol s^-1^, *T*_1/2, *A*_ = 5×10^4^ s, and *θ* _1/2,*B*_ =10^7^ s. Other parameters are as specified in Table 2.

**Fig. S15.**
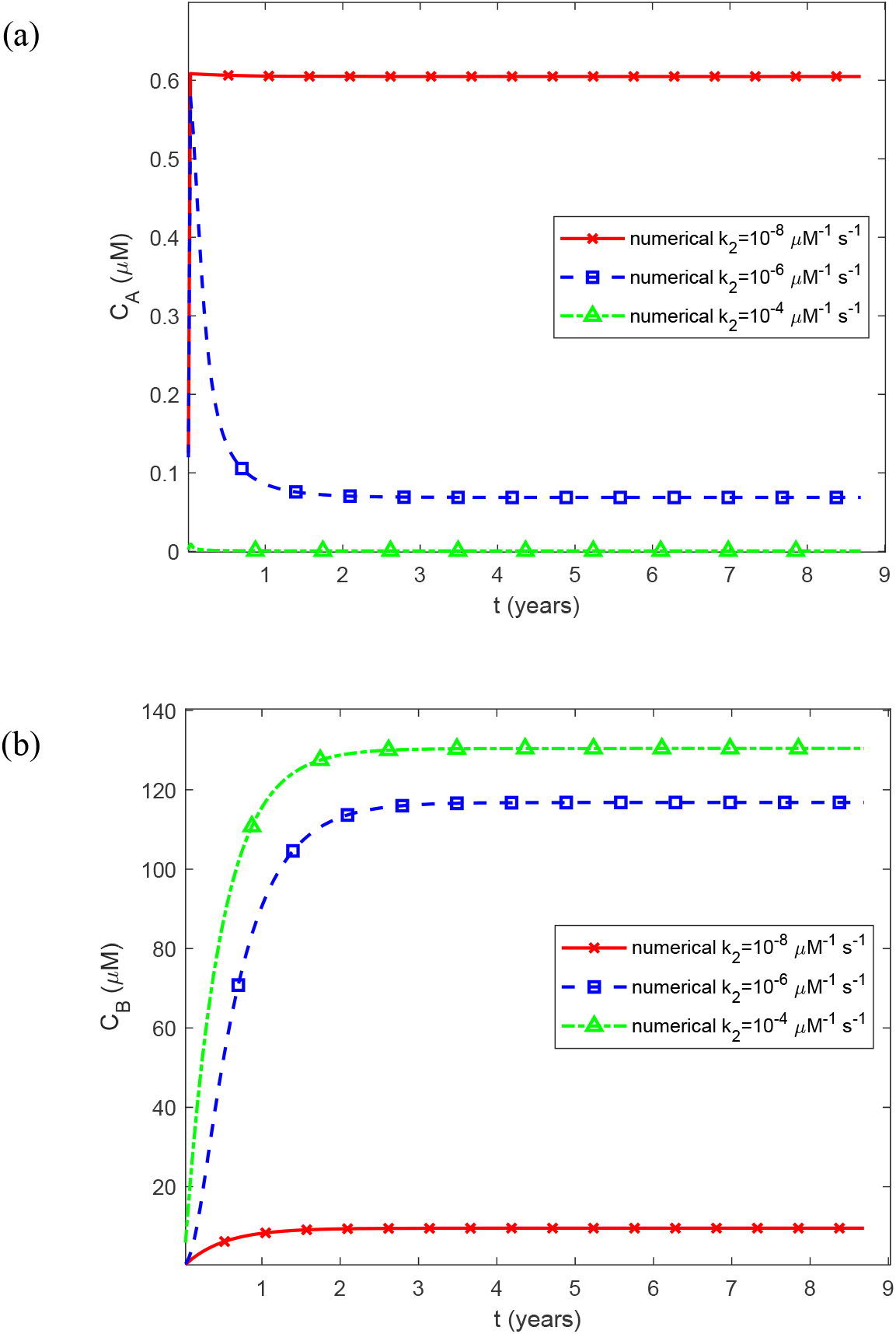
(a) The molar concentration of TAF15 monomers, *C*_*A*_ , plotted against time; and (b) the molar concentration of free TAF15 aggregates (not deposited into inclusions), *C*_*B*_ , plotted against time, for various values of *k*_2_ . Other than for *k*_2_ , physiologically relevant parameter values are utilized: *k*_1_= 10 ^−6^ s^-1^, *q* = 3.79×10^−23^ mol s^-1^, *T*_1/2, *A*_ = 5×10^4^ s, and *θ*_1/2,*B*_ =10^7^ s. Other parameters are as specified in Table 2.

**Fig. S16.**
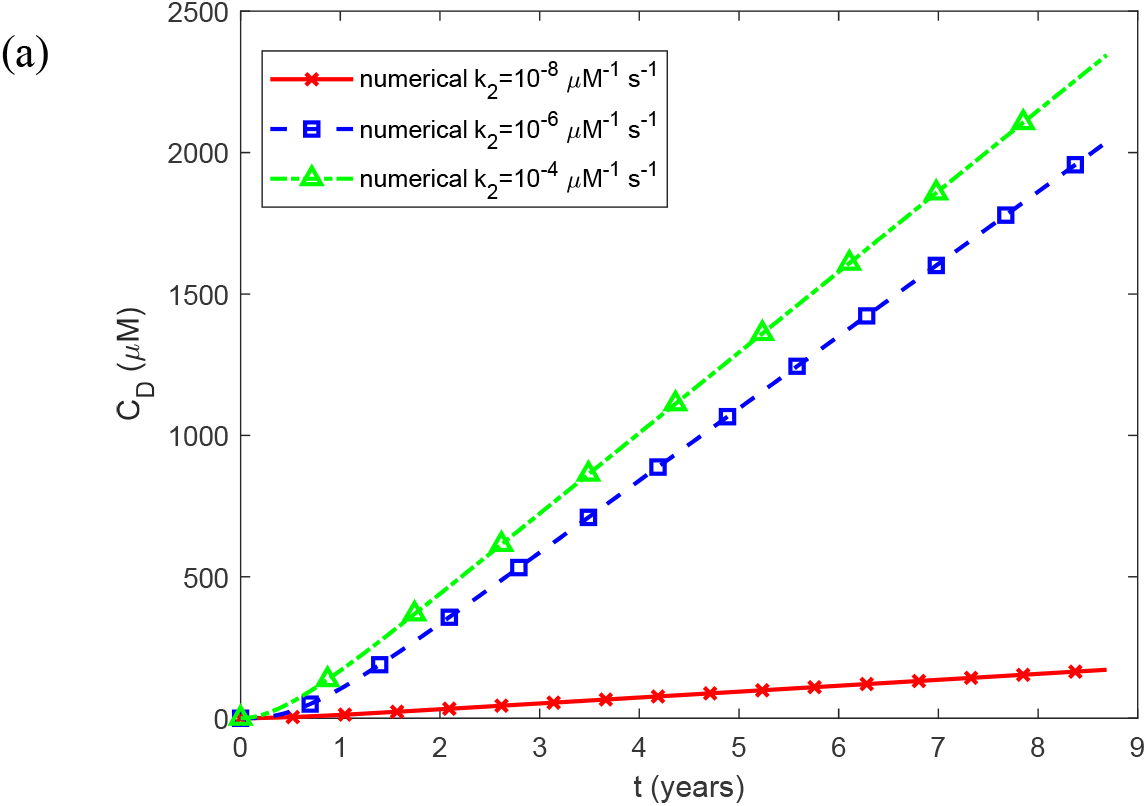
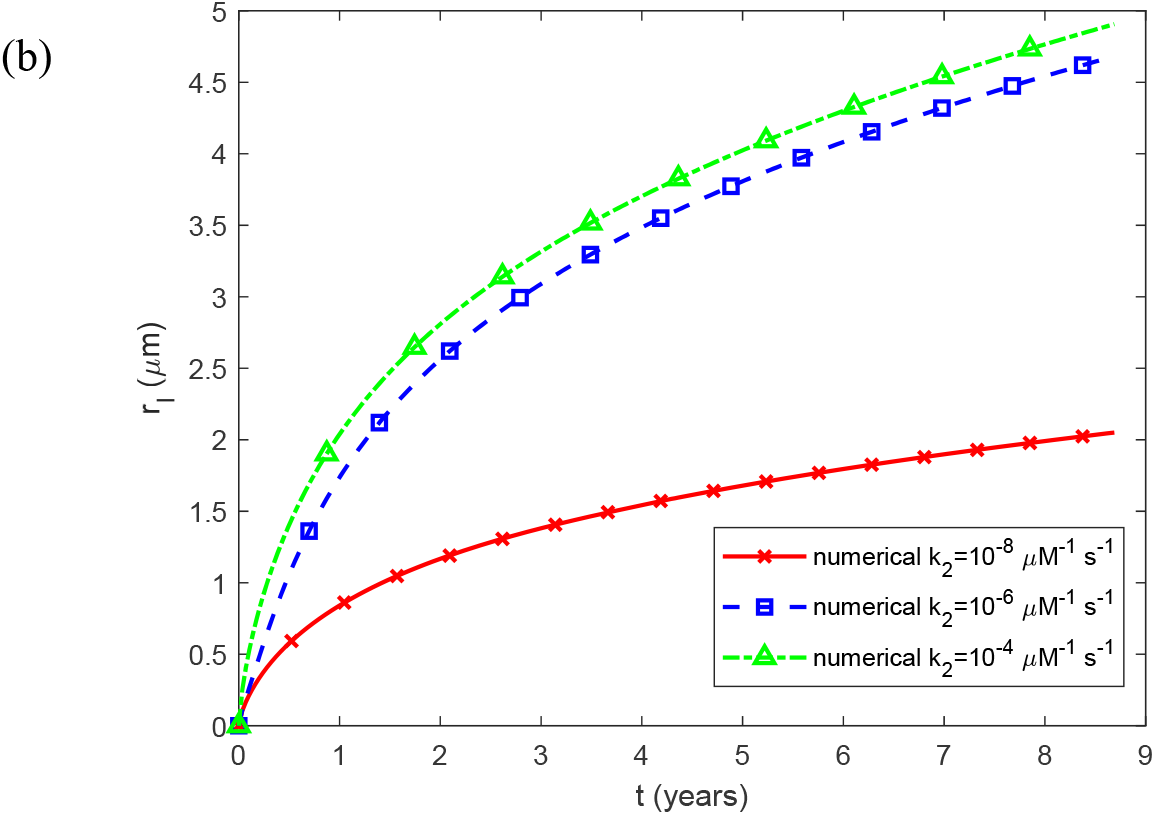
(a) The molar concentration of TAF15 aggregates deposited into TAF15 inclusions, *C*_*D*_ , plotted against time; and (b) the radius of a TAF15 inclusion, *r*_*I*_ , plotted against time, for various values of *k*_2_ .Other than for *k*_2_ , physiologically relevant parameter values are utilized: *k*_1_ = 10 ^−6^ s^-1^, *q* = 3.79×10^−23^ mol s^-1^, *T*_1/2, *A*_ = 5×10^4^ s, and *θ*_1/2,*B*_ =10^7^ s. Other parameters are as specified in Table 2.

**Fig. S17.**
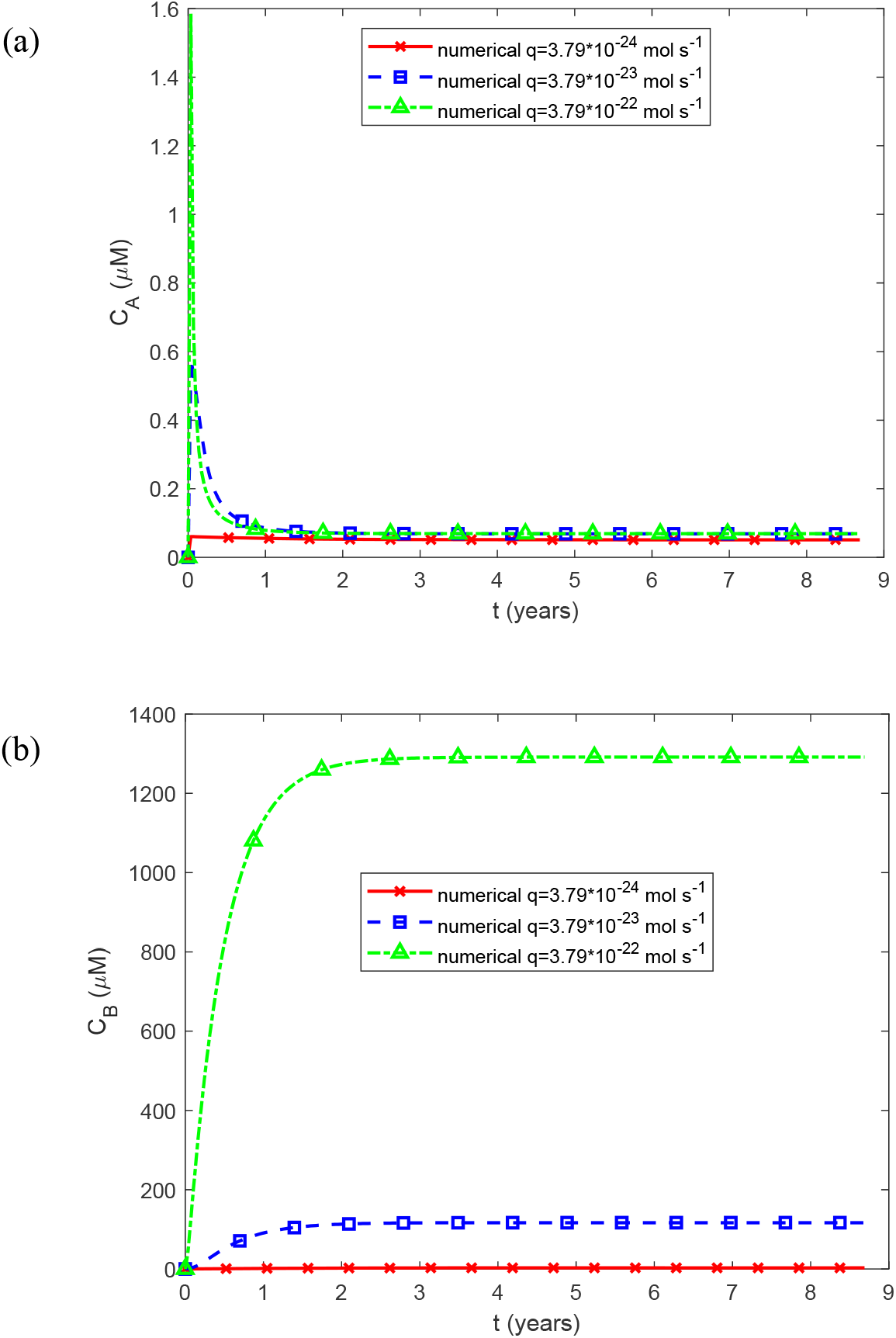
(a) The molar concentration of TAF15 monomers, *C*_*A*_ , plotted against time; and (b) the molar concentration of free TAF15 aggregates (not deposited into inclusions), *C*_*B*_ , plotted against time, for various values of *q* . Other than for *q* , physiologically relevant parameter values are utilized: *k*_1_ = 10 ^−6^ s^-1^, *k*_2_= 10^−6^ μM^-1^ s^-1^, *T*_1/2, *A*_ = 5×10^4^ s, and *θ* _1/2,*B*_ =10^7^ s. Other parameters are as specified in Table 2.

**Fig. S18.**
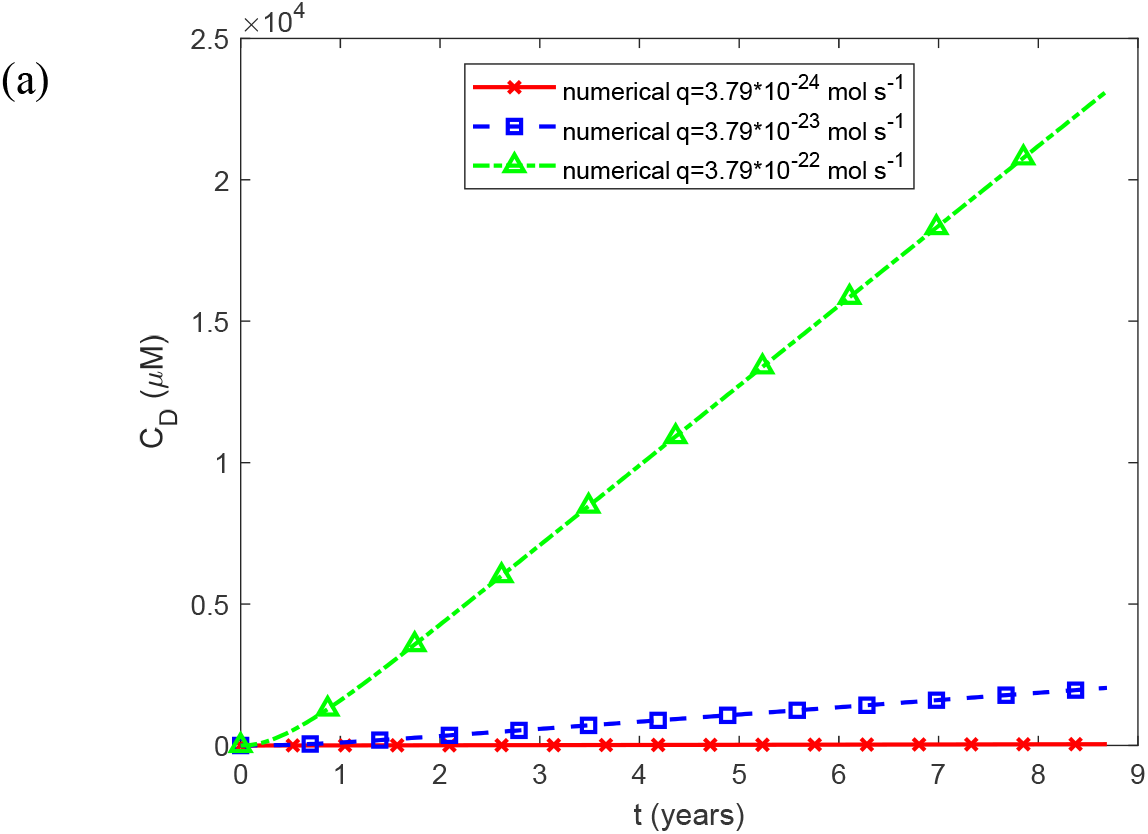
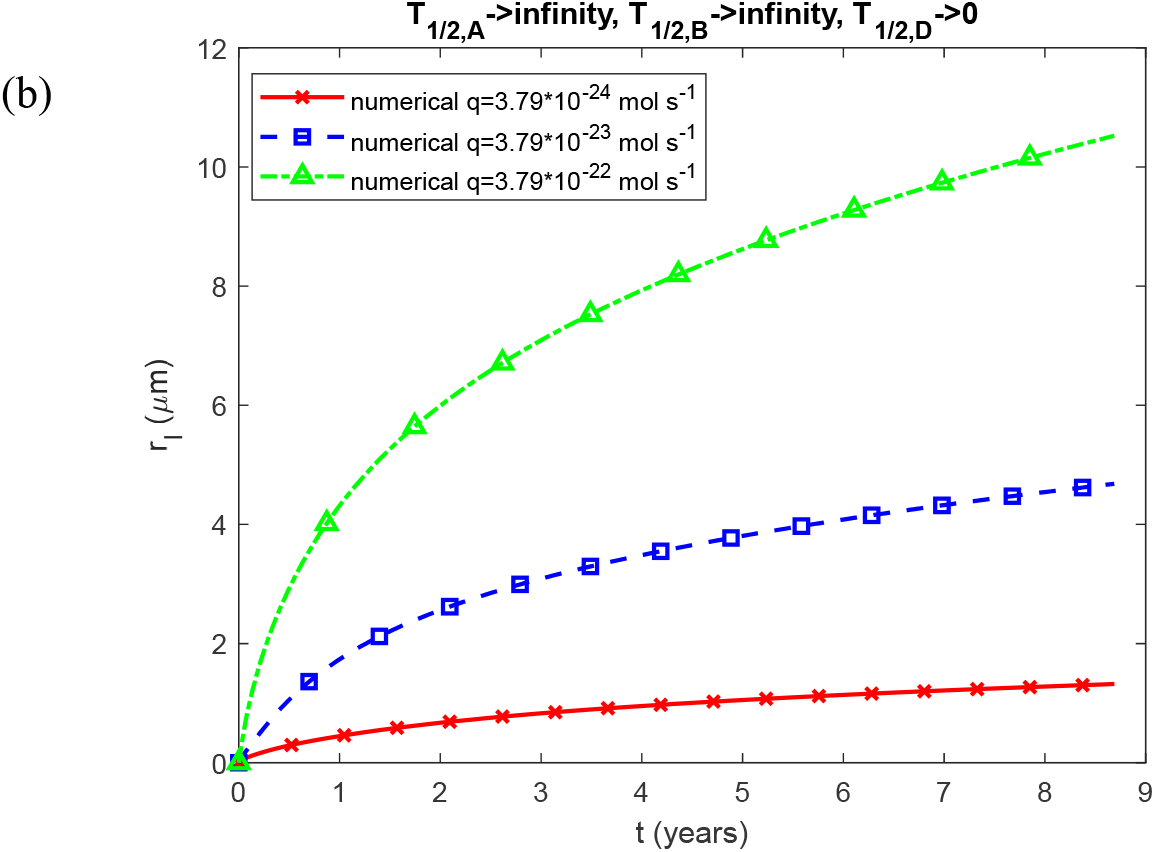
(a) The molar concentration of TAF15 aggregates deposited into TAF15 inclusions, *C*_*D*_ , plotted against time; and (b) the radius of a TAF15 inclusion, *r*_*I*_ , plotted against time, for various values *q* .Other than for *q* , physiologically relevant parameter values are utilized: *k*_1_ = 10 ^−6^ s^-1^ ,*k*_2_ = 10^−6^ μM^-1^ s^-1^, *T*_1/2, *A*_ = 5×10^4^ s, and *θ* _1/2,*B*_ =10^7^ s. Other parameters are as specified in Table 2.

**Fig. S19.**
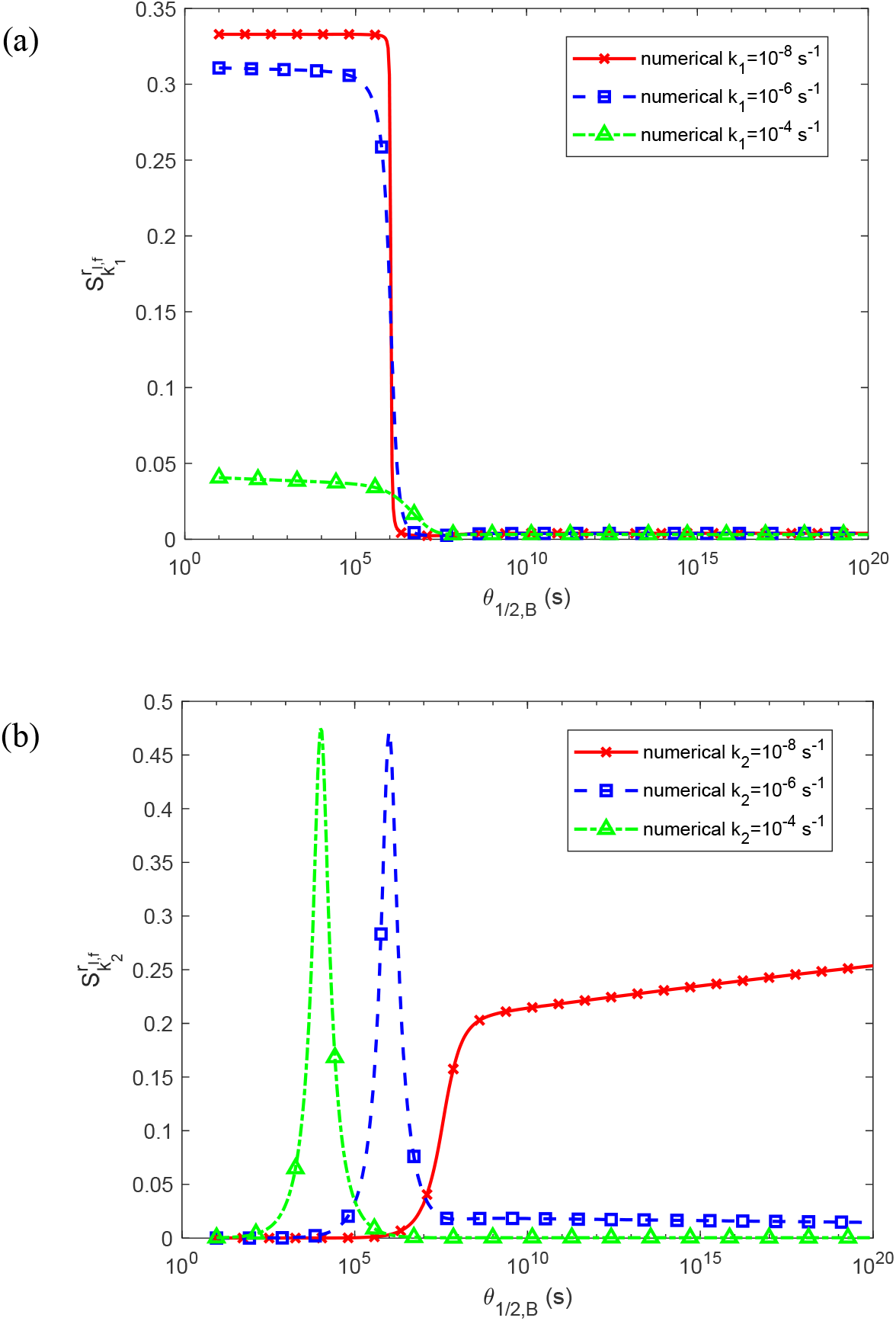
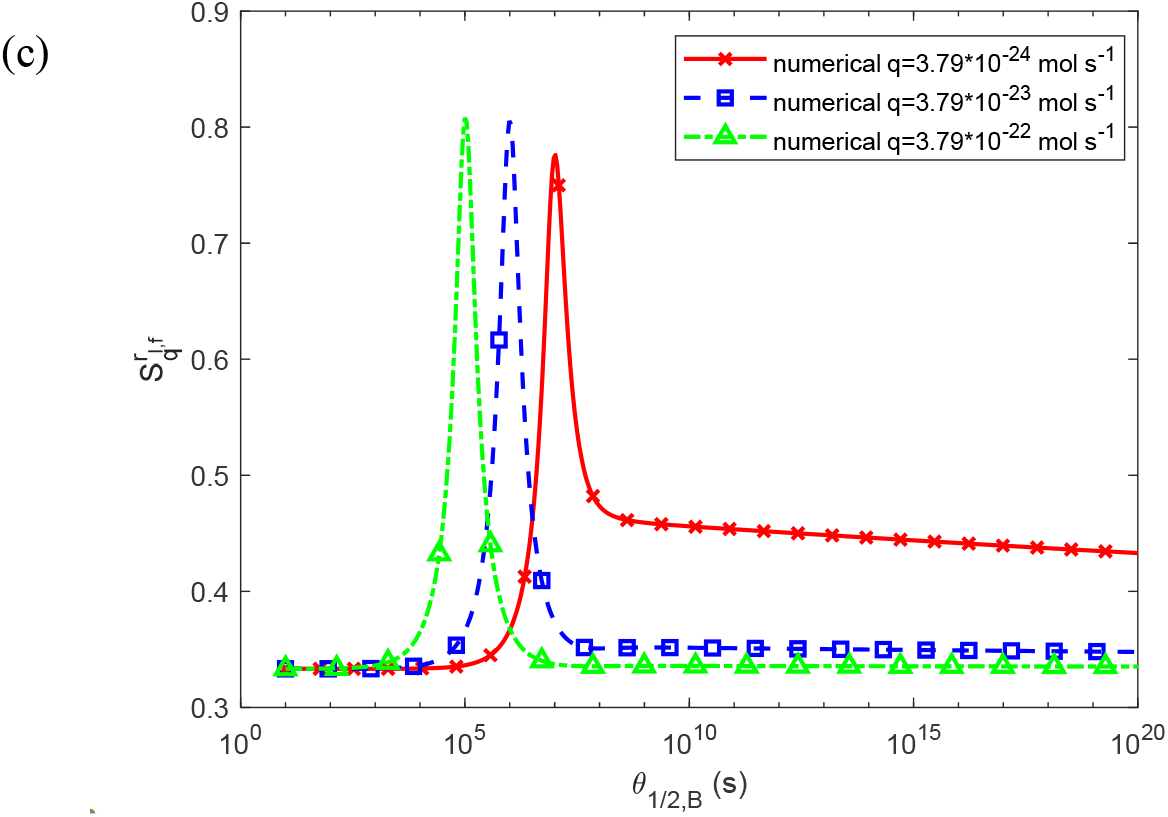
(a) The dimensionless sensitivity of the radius of a TAF15 inclusion at *t*_*f*_ , *r* _*I* , *f*_ , to 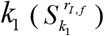 ,for various values of *k*_1_ . Except for *k*_1_ , physiologically relevant parameter values are used: *k*_2_ = 10^−6^ μM^-1^ s^-1^, *q* = 3.79×10^−23^ mol s^-1^, and *T*_1/2, *A*_ = 5×10^4^ s. Other parameter values are as specified in Table 2. (b) The dimensionless sensitivity of *r* _*I* , *f*_ to 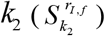 , for various values of *k*_2_ . Except for *k*_2_ , physiologically relevant parameter values are used *k*_1_ = 10^−6^ s^-1^, *q* = 3.79×10^−23^ mol s^-1^, and *T*_1/2, *A*_ = 5×10^4^ s. Other parameter values are as specified in Table 2. (c) The dimensionless sensitivity of *r* _*I* , *f*_ to 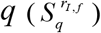 for various values of *q* . Except for *q* , physiologically relevant parameter values are used: *k*_1_ = 10^−6^ s^-1^ , *k*_2_ = 10^−6^ *μ*M s^-1^ , and *T*_1/2, *A*_ = 5×10 s. Other parameter values are as specified in Table 2.

